# A single-cell map of hypertension

**DOI:** 10.1101/2024.12.25.630332

**Authors:** Qiongzi Qiu, Yong Liu, Hong Xue, Rajan Pandey, Lishu He, Jing Liu, Pengyuan Liu, Bhavika Therani, Vinod Kumar, Jing Huang, Maya Guenther, Kristie Usa, Michael Grzybowski, Mark A. Vanden Avond, Andrew S. Greene, Allen W. Cowley, Sridhar Rao, Aron M. Geurts, Mingyu Liang

## Abstract

Hypertension is a leading risk factor for disease burden and death worldwide. Several organ systems are involved in the development of hypertension, which contributes to stroke, heart disease, and kidney disease. Despite the broad health relevance, our understanding of the molecular landscape in hypertension is limited and lags other major diseases. Here we report an extensive analysis of the molecular landscape in hypertension and its end-organ damage and uncover novel mechanisms linking human genetic variants to the development of these diseases. We obtained single-nucleus RNA-seq (612,984 nuclei), single-nucleus ATAC-seq (179,637 nuclei), or spatial transcriptome data from five organs (hypothalamus, kidney, heart, 3^rd^ order mesenteric artery, middle cerebral artery) in three mouse and rat models under twelve experimental conditions. More than one third of all hypertension research in animal models involves these three models. We identified both model-specific and convergent responses in cell types, genes, and pathways. By integrating our data with human genomic data, we partitioned the blood pressure and end-organ damage traits into cell type-specific transcriptional contributions and cell types common across multiple traits. Using genomic editing in animal models and human induced pluripotent stem cells, we extended key findings and identified new mechanisms linking human genetic variants to the development of hypertension and related renal injury. We anticipate that our rich data sets and findings will broadly drive forward the research of hypertension and hypertensive end-organ damage. Our approach of integrating multi-model and multi-tissue single-cell analysis with human genetic data and in vivo and in vitro genome editing can be applied to investigate other complex traits.

## Introduction

Hypertension is the leading global cause of disease burden and mortality, affecting around 33% of adults worldwide and contributing significantly to healthcare costs through complications such as heart disease, stroke, and kidney failure^1^. Most hypertensive patients lack a cure, and millions remain hypertensive despite multi-drug treatments^2^. Despite its widespread impact, our molecular understanding of hypertension and its associated end-organ damage lags behind other major diseases, limiting progress in therapeutic discovery and precision medicine.

Hypertension is a complex disorder involving multiple organ systems including the kidneys, cardiovascular system, central nervous system, and immune system^3^. The heterogeneity, including diverse causes and organ-specific effects, complicates the identification of shared molecular mechanisms. Additionally, the cellular complexity within each affected organ makes it challenging, yet crucial, to pinpoint the specific cell types driving disease progression. A systems-level approach across multiple animal models and relevant tissues is needed to address these challenges and reveal the molecular landscape underlying the development of hypertension and its end-organ damage.

To address this gap, we developed an extensive single-cell gene expression atlas across three widely used animal models of hypertension: angiotensin II-treated mice, Dahl salt-sensitive (SS) rats, and spontaneously hypertensive rats (SHR)^4^. These models account for over 35% of hypertensive animal model studies based on PubMed searches. Our multi-tissue analysis encompasses key organs involved in the development of hypertension and its end-organ damage, including the hypothalamus, kidney, 3^rd^ order mesenteric artery, heart, and middle cerebral artery. We generated high-resolution molecular profiles of these organs during hypertension progression using single-nucleus RNA-sequencing (snRNA-seq), supplemented with snMultiome-seq and spatial transcriptomics in select tissues, revealing a rich landscape of gene expression changes and cellular dynamics across models. By incorporating normotensive control strains (Sprague-Dawley and Wistar-Kyoto rats), we differentiated hypertension-specific molecular changes from those induced by genetic or environmental factors.

Our single-cell approach enabled the identification of both model-specific and convergent cellular responses, as well as key genes and pathways. Comparing hypertensive and normotensive strains revealed baseline molecular differences that correlate with hypertension susceptibility and disease progression. The inclusion of progressive time-point data further enabled us to track dynamic molecular changes as hypertension developed. By integrating our single-cell data with human genomic datasets, we identified cell type-specific transcriptional contributions to blood pressure and end-organ damage traits, identifying both known and novel vulnerable cell populations. This integrative analysis further nominated key SNP-gene pairs associated with hypertension or its end-organ damage. Using genomic editing in animal models and human induced pluripotent stem cells, we discovered novel mechanisms by which these SNPs drive the development of hypertension and related organ damage.

## RESULTS

### Study design and cell type characterization in hypertension-relevant tissues

This study utilized three widely used animal models of hypertension: angiotensin II-treated C57BL/6 mice, salt-sensitive (SS) rats subjected to a high-salt (4% NaCl) diet, and spontaneously hypertensive rats (SHR), along with their respective controls, Sprague-Dawley (SD) and Wistar-Kyoto (WKY) rats (Fig. 1a, Supplementary Table 1). In total, 96 animals were used, divided into 12 treatment groups (N=8 per group). As expected, hypertensive models developed elevated blood pressure (BP) over the course of the experiments, and showed end-organ damage, including albuminuria and fibrosis in the kidney, cardiomyocyte hypertrophy and fibrosis in the heart, and media thickening in the mesenteric artery. In contrast, BP of the controls remained stable or, in the case of aged WKY rats, showed a slight increase (Supplementary Fig. 1-6).

**Fig. 1.**
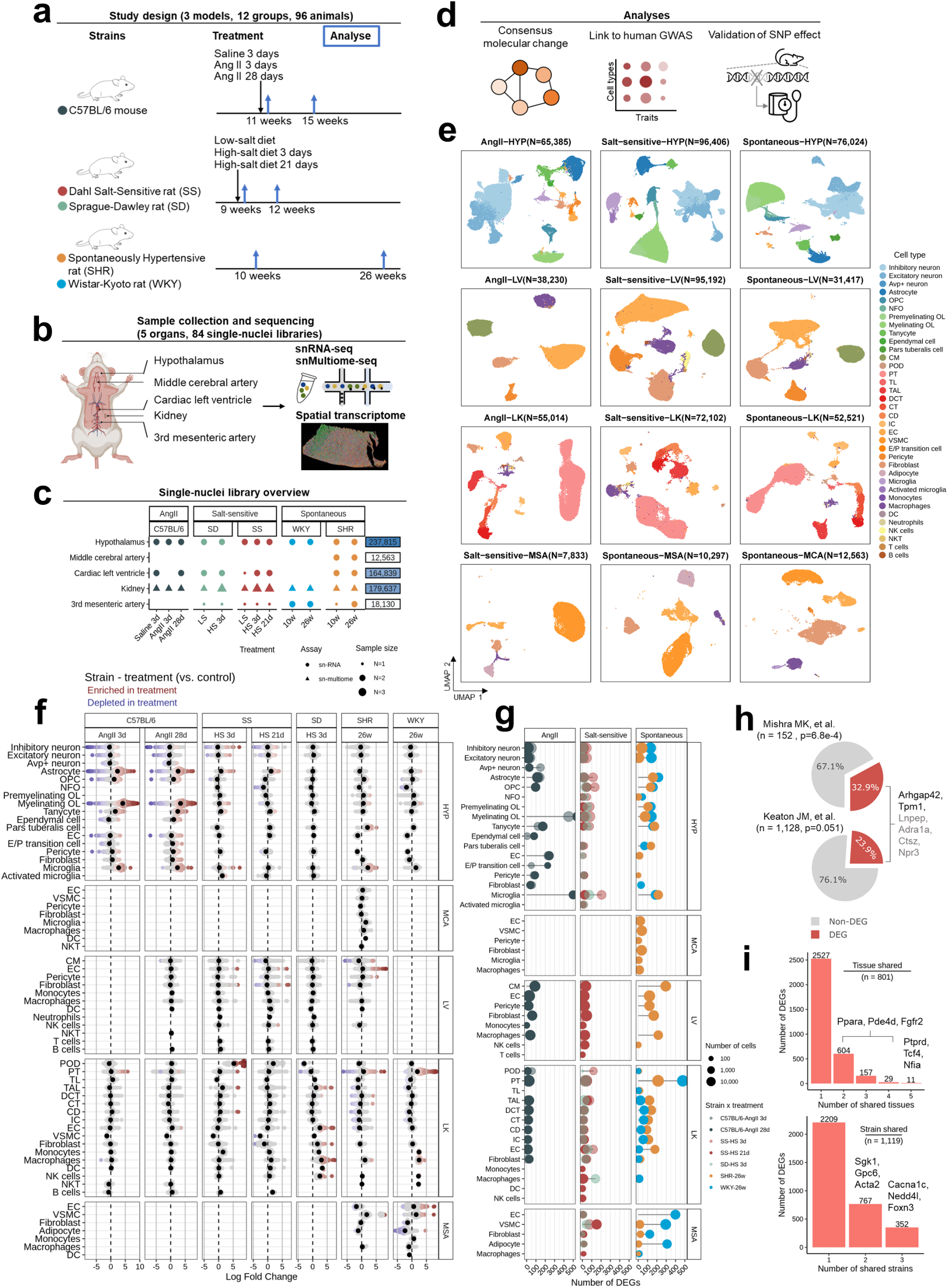
Study design, cell type characterization, and global cellular and molecular changes in hypertension. (a) Schematic of the study design using three animal models to study hypertension and hypertensive end-organ damage. (b) Overview of tissue collection and sequencing methods used for hypertension-related organs. (c) Summary of single-nuclei libraries generated from five tissues. N refers to the number of libraries with each library prepared from a median of 4 animals. (d) Overview of the main analyses performed in the study. (e) UMAP plots showing major cell types identified in each tissue and model. (f) Beeswarm plots displaying log fold change in cell neighborhood abundance for each treatment group versus controls. Colored spots represent neighborhoods that are differentially abundant with spatial FDR < 0.1. (g) Lollipop plot showing the number of differential expressed genes (DEGs) identified in each treatment group versus controls across cell types, tissues and models. The AngII column includes data from angiotensin II (AngII)-treated C57BL/6 mice compared to saline 3d. The salt-sensitive column includes data from Dahl salt-sensitive (SS) rats and Sprague-Dawley (SD) rats on high-salt diets compared to their respective low-salt group. The spontaneous column includes data from 26-week-old spontaneously hypertensive (SHR) rats and Wistar-Kyoto (WKY) rats compared to their respective 10-week-old control. (h) Pie chart displaying the proportion of DEGs overlapping with curated blood pressure-relevant gene lists. DEGs are typically defined by an absolute log2 fold change greater than 0.5 and a Bonferroni-adjusted p-value less than 0.05 (Wilcoxon rank sum test). Genes with a fold change greater than 0.25 are also included for broader visualization in panel (h) and are highlighted in grey. For each tested gene list, numbers represent genes expressed in our snRNA-seq data. P-values were derived from a one-tailed hypergeometric test, comparing the proportion of DEGs in the tested gene list to the proportion of DEGs across all detected genes. (i) Bar plots categorizing DEGs based on the number of tissues and strains they are shared across. Tissue abbreviations: HYP (hypothalamus), MCA (middle cerebral artery), LV (left ventricle), LK (left kidney), and MSA (3^rd^ mesenteric artery). Cell type abbreviations: OPC (oligodendrocytes progenitor cell), NFO (newly formed oligodendrocyte), OL (oligodendrocyte), CM (cardiomyocyte), POD (podocyte), PT (proximal tubule), TL (thin limb), TAL (thick ascending limb), DCT (distal convoluted tubule), CT (connecting tubule), CD (collecting duct), IC (intercalated cell), EC (endothelial cell), VSMC (vascular smooth muscle cell), E/P transition cell, DC (endothelial/pericyte transition cell), NK cells (natural killer cell), NKT (natural killer T cell).

Tissues from the hypothalamus, middle cerebral artery, cardiac left ventricle, kidney, and the 3^rd^ order mesenteric artery were collected at multiple time points (Fig. 1a and b). From these, nuclei were isolated and pooled from a median of 4 animals per library to generate 57 snRNA-seq libraries, with 27 additional snMultiome-seq libraries created specifically for kidney samples (Fig. 1c, Supplementary Table 1). Spatial transcriptomics were performed on two selected kidney samples using Stereo-seq (Fig. 1b, Supplementary Table 1). Bulk RNA-seq was performed on the same cohort by pooling tissues from two animals per library for transcriptional validation (Supplementary Table 1).

We retained RNA-seq data from a total of 612,984 high-quality nuclei for downstream analysis (Fig. 1d, Supplementary Table 2, Methods). Clustering and annotation were performed separately for each tissue and model to preserve model-specific characteristics (Fig. 1e, Supplementary Fig. 7, Supplementary Table 3 and 4). In total, 36 major cell types were identified across the datasets using well-curated canonical markers (Supplementary Fig. 7 and 8).

### Global cellular and gene expression shifts in hypertension

We assessed changes in cell type abundances by comparing hypertensive or treatment groups with their corresponding baseline controls using Milo (Fig. 1f). Milo detects differential abundance by grouping cells into neighborhoods and estimating their proportions^5^. The most pronounced cellular shifts occurred in the hypothalamus of Ang II-treated mice, where we observed great reductions in neuronal proportion in treatment groups, coupled with a predominant increase of glial cells, including astrocytes, myelinating oligodendrocytes, microglia, and tanycytes. Within astrocytes and myelinating oligodendrocytes, we also detected shifts in subpopulations, suggesting potential cell state transitions. A similar increase in astrocytes was observed in SS rats in response to the high-salt diet but was absent in the salt-resistant SD strain. Enrichment of microglia was consistently seen across all three models following hypertension onset. Notably, a distinct cluster of *Cd74*+ activated microglia emerged specifically in the salt-sensitive model, indicating a unique immune activation response (Supplementary Fig. 9a). In the kidney, enrichment of proximal tubular (PT) cell groups was observed across multiple strains following treatment. In the SHR model, additional decrease in PT subpopulations, as well as in various epithelial cell types, was identified. In the cardiac left ventricle, a small subset of endothelial cell (EC) subpopulations enriched in hypertensive animals was observed in both SS and SHR rats.

Next, we identified differentially expressed genes (DEGs) in each cell type in each group compared to baseline control (Fig. 1g, Supplementary Table 5). Consistent with the observed changes in cell proportions, transcriptional alterations in the mouse hypothalamus were highly dynamic, with the highest number of DEGs detected in myelinating oligodendrocytes, microglia, and ECs. The salt-sensitive model showed relatively modest transcriptional changes across tissues, despite pronounced phenotypic alterations. In salt-induced hypertension, the most transcriptionally active cell types were microglia in the hypothalamus, ECs in the kidney, and vascular smooth muscle cells (VSMCs) in the mesenteric artery. Notably, in the hypothalamus, more DEGs were identified at the early stage of hypertension (high-salt diet, 3 days) compared to later stages (high-salt diet, 21 days), suggesting an acute transcriptional response. In the spontaneous hypertension model, widespread gene expression dysregulation was observed across multiple tissues, including the hypothalamus, cardiac left ventricle, and kidney. In particular, the kidney showed a higher number of DEGs in the SHR model compared to WKY controls across most cell types. These findings revealed shared and distinct patterns of cellular and gene expression changes across models and tissues, providing a foundation for further exploration of underlying biological processes.

We found that 32.9% and 23.9% of BP-relevant genes curated by Mishra et al and Keaton et al^6,7^, respectively, were differentially expressed in our dataset, significantly higher than the overall DEG rate (Fig. 1h). Here, we highlighted three well-known BP-regulating genes: *Arhgap42*, *Npr3*, and *Adra1a*.

*Adra1a* mediates vasoconstriction through G-protein-coupled adrenergic signaling and is a target gene for BP medications alpha blockers^8^. *Arhgap42* regulates vascular smooth muscle contraction via RhoA signaling, critical for maintaining vascular tone^9^. *Npr3* modulates natriuretic peptide clearance and promotes vasodilation, both important for BP regulation^10,11^. We then investigated DEGs shared across tissues and strains, and uncovered several known BP relevant genes, including *Nedd4l*^12^, *Sgk1*^12^, *Cacna1c*^13^, *Ppara*^14,15^, *Pde4d*^16^, and *Ptprd*^15^ (Fig. 1i). For example, *Nedd4l* and *Sgk1* have well-established roles in regulating BP and sodium homeostasis^12,17^. Additionally, we observed that DEGs shared across multiple tissues or strains were more likely to overlap with curated BP-relevant gene sets, suggesting they may represent conserved and robust regulatory mechanisms in hypertension across different biological contexts (Supplementary Fig. 9b).

### Consensus molecular changes in the hypothalamus

Given the hypothalamus’ important role in BP regulation and the substantial cellular and transcriptional changes observed, we then focused on consensus changes in this region to identify biological processes consistently altered across models and cell types in hypertension. A total of 208 genes were differentially expressed in more than 10 cell types, including several associated with lipid metabolism and inflammation, such as *Rora*, *Apoe*, *Ptgds*, *Csmd1*, and *Zbtb16* (Fig. 2a, Supplementary Fig. 10a). *Apoe* was notably among the most upregulated genes in astrocytes, OPCs, and activated microglia in the salt-sensitive model (Supplementary Fig. 10b). Genes linked to oxidative stress, such as *Fth1* and *Hsp90ab1*, also showed recurrent differential expression. *Hsp90ab1*, a heat shock protein family member, showed upregulation in multiple cell clusters in both Ang II and salt-sensitive models (Fig. 2b). Bulk RNA-seq from the same animal cohort validated many of these pan-cell-type gene expression changes, including the strong upregulation of *Zbtb16* and *Lars2* in mouse hypothalamus after Ang II treatment (Fig. 2c).

**Fig. 2.**
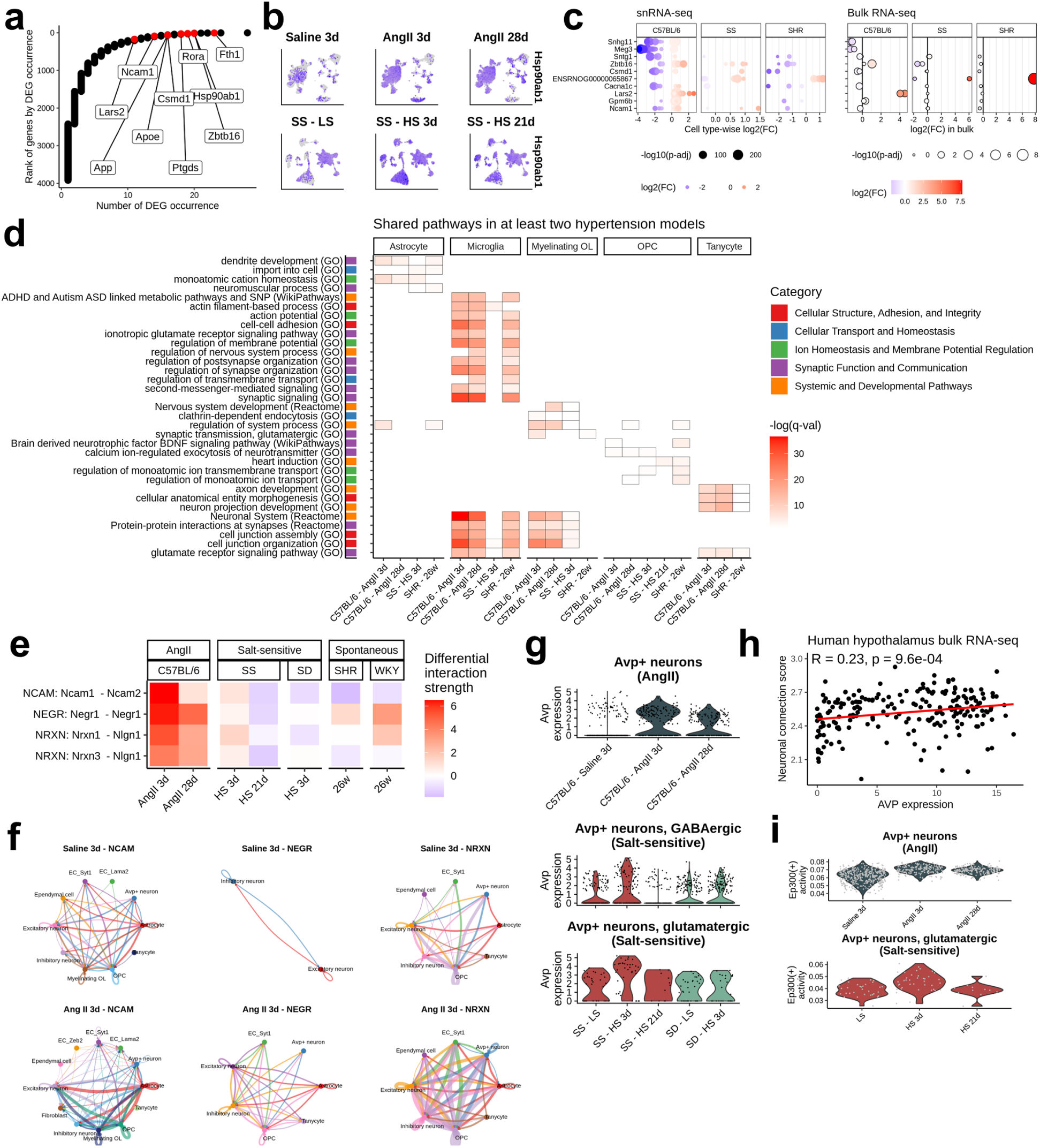
Consensus molecular changes in the hypothalamus. (a) Rank plot displaying the rank of DEGs by the number of occurrences across cell types and models in hypothalamus. (b) UMAP plots showing *Hsp90ab1* expression changes across conditions in AngII-treated mice and SS rats on a high-salt diet. (c) Top 10 genes with consistent expression changes between snRNA-seq (left) and bulk RNA-seq (right) across models. In the snRNA-seq panel, each row represents a gene, with dots indicating log2 fold changes for significant DEGs across all cell types in each comparison (Bonferroni-adjusted p-value<0.05, Wilcoxon rank sum test). In the bulk RNA-seq panel, each dot represents the log2 fold change for significant DEGs in any comparison (BH-adjusted p-value<0.05, DESeq2’s Wald test). Genes are ordered by DEG occurrence in the snRNA-seq data. (d) Pathway and GO terms enriched in DEGs of each cell type within each comparison, showing pathways identified in at least two models. (e) Heatmap of differential cellular interaction strength between treatment groups and controls across models, with signaling pathways displaying consistent changes in at least two models. (f) Circle plots illustrating ligand-receptor interactions from pathways in panel (e) in mice treated with saline 3d and AngII 3d, with edge width representing communication strength. (g) Violin plot showing *Avp* expression changes across conditions in neuron subtypes specifically expressing *Avp*. (h) Dot plot showing Pearson correlation between neuronal connection score and *AVP* expression in human hypothalamus bulk RNA-seq data from Genotype-Tissue Expression (GTEx). (i) Violin plot displaying Ep300 activity changes in *Avp*+ neurons across conditions.

As mentioned above, a distinct cluster of *Cd74*+ activated microglia was specifically identified in the salt-sensitive model (Supplementary Fig. 9a). Furthermore, immune-related genes, such as *B2m*, *C1qa*, *C1qb*, and *C1qc*, were exclusively upregulated in microglia after 3 days of high-salt diet in SS rats, but not in SD rats, or animals from other models (Supplementary Fig. 10c), indicating a unique early immune response in microglia during high-salt diet-induced hypertension.

Consensus pathways and biological processes identified in at least two models were prevalent across various glial cells, including astrocytes, OPCs, myelinating oligodendrocytes, tanycytes, and microglia (Fig. 2d, Supplementary Fig. 10d). These changes were broadly related to synaptic function and communication, ion homeostasis and membrane potential regulation, and cellular adhesion and junction integrity, suggesting a coordinated effort among glial cells to maintain neural stability under hypertensive conditions. Moreover, cell-cell communication analysis across three models demonstrated recurrent upregulation in the interaction strength of gene pairs within neuronal communication pathways, including NCAM, NEGR, and NRXN, particularly during the early stages of hypertensive treatment (Ang II 3 days or high-salt 3 days) (Fig. 2e, Supplementary Fig. 11). These elevated signaling networks involved multiple glial cells, excitatory and inhibitory neurons, and magnocellular *Avp*+ neurons (Fig. 2f, Supplementary Fig. 11).

Concurrently, early-stage elevation of *Avp* expression in *Avp*+ neurons was observed in both Ang II and salt-sensitive models (Fig. 2g). We hypothesized that increased neuronal communication, quantified by a neuronal communication score based on genes from the NCAM, NEGR, and NRXN pathways (Methods), might correlate with *Avp* expression. This hypothesis was supported by a significant correlation between *AVP* expression and the neuronal communication score in human hypothalamus bulk RNA-seq data from The Genotype-Tissue Expression project (GTEx)^18^ (Fig. 2h). To further explore direct regulators of *Avp* expression during hypertension, we used SCENIC analysis and identified *Ep300* as a strong candidate^19^. Elevated *Ep300* activity was observed in magnocellular neurons within the hypothalamus in both mice treated with Ang II and SS rats on a high-salt diet, especially at the early stage of hypertension progression (Fig. 2i).

### Mecom+ endothelial cells linking baseline deficiency to hypertension response in the kidney

The kidney both drives and is affected by hypertension. We conducted pathway enrichment analysis on DEGs in the kidney identified from hypertensive animals compared to controls, as well as baseline comparisons between hypertensive and normotensive strains (Fig. 3a). Notably, baseline differences in biological processes were primarily observed in kidney endothelial cells, with several processes, including circulatory system processes, cell junction organization, and intracellular signal transduction, also showing alterations after hypertension onset in SS and SHR rats (Fig. 3b). A consistent gene expression pattern was observed within these processes, where most genes were less abundant in SS rats compared to SD rats at baseline, followed by an increase post-hypertension onset.

**Fig 3.**
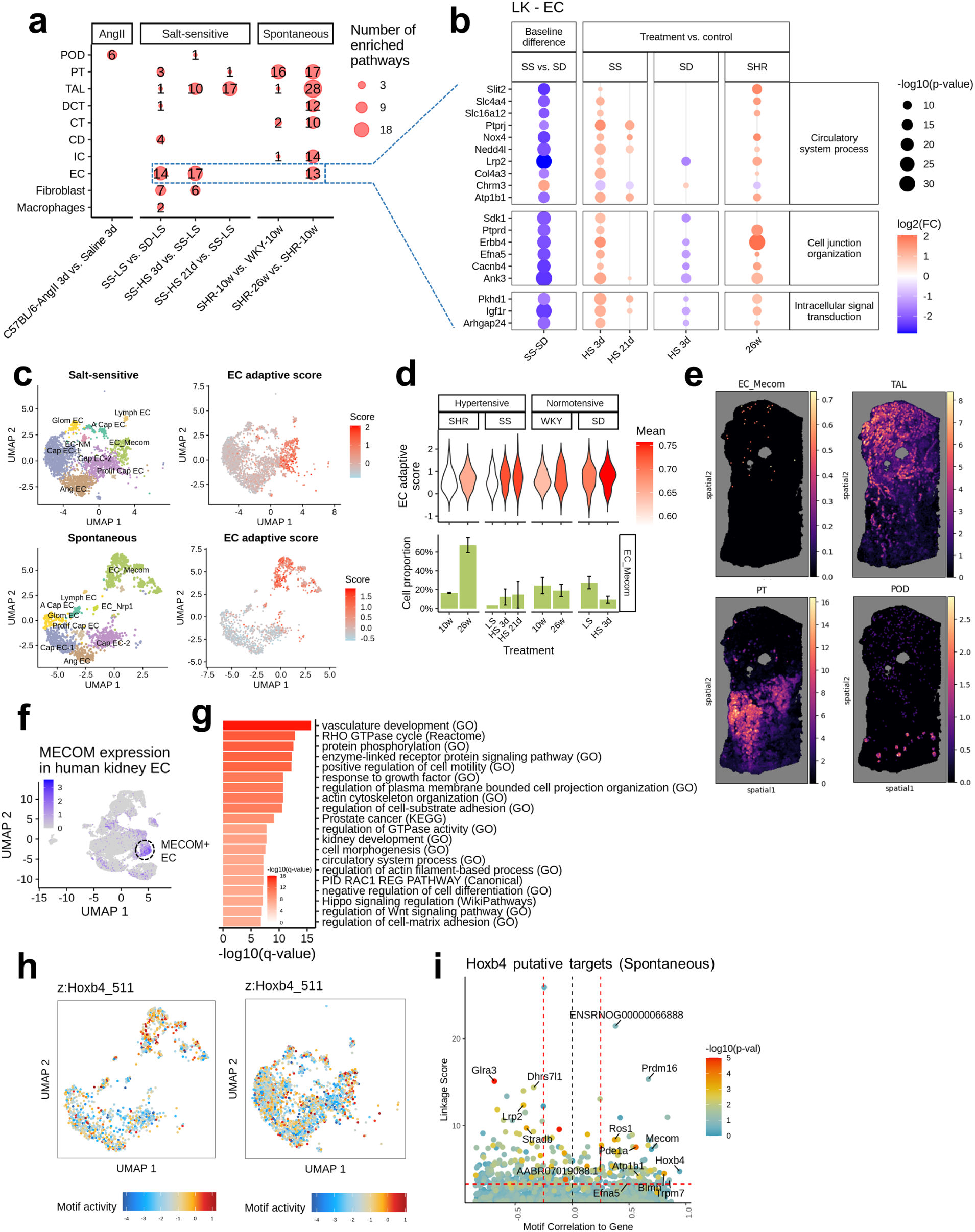
*Mecom*+ endothelial cell (EC) as a baseline-deficient and hypertension-responsive cell type in the kidney. (a) Dot plot showing the number of pathways enriched in cell-type-specific DEGs in the kidney across comparisons, including treatment vs. control within each strain and hypertensive vs. normotensive strains at baseline. (b) Dot plot displaying expression changes of genes from selected pathways in panel (a) in kidney EC, comparing treatment vs. control within each strain and hypertensive vs. normotensive strains at baseline. (c) UMAP plots of kidney EC subtypes (top left) and EC adaptive score (top right) in SS and SD, and EC subtypes (bottom left) and EC adaptive score (bottom right) in SHR and WKY. (d) Violin plot showing EC adaptive score (top) and bar plot showing the proportional changes of *Mecom*+ EC (bottom) across conditions in salt-sensitive and spontaneous models and their normotensive controls. Bar plots are shown as mean ± s.e.m. (e) Spatial distribution of estimated abundance of *Mecom*+ EC, TAL, PT, and POD in kidney sections from a salt-sensitive rat on a high-salt diet, as inferred using cell2location with spatial transcriptomics data. (f) UMAP plots showing *MECOM* expression in ECs from human kidney scRNA-seq data. (g) Bar plot of pathway enrichment using genes specifically expressed in the *MECOM*+ EC subtype compared to other ECs from human kidney data (BH-adjusted p-value<0.05, Wilcoxon rank sum test). Enrichment p-values were calculated using the hypergeometric test. (h) UMAP plots showing TF motif activity deviation z-score of Hoxb4_511 across ECs in the spontaneous (left) and salt-sensitive hypertension models (right). (i) Prioritization of gene targets for *Hoxb4* in spontaneous model. The x-axis represents the Pearson correlation between TF motif activity and integrated gene expression across ECs. The y-axis shows the TF linkage score, calculated as the sum of scaled motif scores for all linked peaks based on peak-to-gene link correlation. The color of the points reflects the hypergeometric enrichment of the TF motif in linked peaks for each gene. The horizontal dashed line marks the 80th percentile of the linkage score, while the vertical dashed lines represent motif-to-gene expression correlations of -0.25 and 0.25, respectively.

We developed an ‘EC adaptive score’ based on the DEGs in Figure 2b and found a specific enrichment of the score in *Mecom*+ ECs (Fig. 3c, Supplementary Fig. 12a, Methods). The elevated EC adaptive score during hypertension reflected both increased gene expression and the expanded proportion of *Mecom*+ ECs. At baseline, the proportion of *Mecom*+ ECs was lower in hypertensive strains compared to normotensive strains, but increased significantly following hypertension onset, particularly in SHR rats (Fig. 3d). Given the milder kidney damage in 26-week-old SHR rats compared to more severe damage in SS rats following a high-salt diet, these findings suggest a potential protective role for *Mecom*+ ECs in maintaining kidney function under hypertensive stress (Supplementary Fig. 6). Spatial transcriptomics localized these cells to the medulla, within the region enriched with thick ascending limbs (Fig. 3d). Parallel analysis in human kidney samples^20^ revealed that *MECOM*+ ECs were enriched for vasculature development and circulatory system processes, functionally overlapping with findings from animal models (Fig. 3f and g). This suggests that *Mecom*+ ECs have a conserved role in vascular remodeling across species, which may have heretofore unrecognized relevance to kidney function.

To investigate the transcriptional regulators driving these changes, we integrated our kidney snATAC-seq and snRNA-seq data. We first performed peak-to-gene linkage analysis at both the general EC level and the EC subtype level to capture a full regulatory relationship. Peaks associated with EC subtype marker genes were analyzed for TF motif enrichment (Supplementary Fig. 12b). For motifs significantly enriched in both SS and SHR models, we calculated z-score differences to identify TFs specific to *Mecom*+ ECs, with several Hox family motifs emerging as key candidates (Fig. 3h, Supplementary Fig. 12c). To further refine the identification of potential TF drivers, we applied the pipeline from Ober-Reynolds et al. Briefly, this pipeline correlates motif activity with gene expression, calculating a ‘linkage score’ for each gene based on linked peaks containing specific TF motifs. For each candidate TF, potential regulatory targets were identified as genes that showed strong correlations between expression and global motif activity (absolute Pearson correlation coefficient > 0.25) and a high linkage score (> 80th percentile). Hoxb4 emerged as a potential regulator as it linked to *Mecom* and multiple genes in the signature of the EC adaptive score, including *Efna4*, *Atp1b1*, *Slc16a12*, and *Lrp2* (Fig. 3i, Supplementary Fig. 12d).

### Cell type linkages to hypertension and end-organ damage through integration with human GWAS data

To better understand the cellular contributions in genetic susceptibility of hypertension and hypertensive end-organ damage, we integrated disease progression transcriptome data from our animal models with human GWAS data and examined trait-cell type and trait-trait relationships (Fig. 4a). We collected 10,253 SNPs associated with nine traits related to BP and hypertensive end-organ damage, including systolic BP, diastolic BP, essential hypertension, pulse pressure, stroke, coronary artery disease (CAD), estimated glomerular filtration rate (eGFR), albuminuria and blood urea nitrogen (BUN), and mapped them to 10,041 human genes using proximal, expressional, and regulatory evidence (Fig. 4a, Methods). A median of 488 SNP-related orthologous genes for each trait were significantly differentially expressed during hypertension progression in our animal models, corresponding to a median of 63.04% of SNPs across traits, which was significantly higher than the proportion observed in traits not directly related to BP or hypertensive end-organ damage (Fig. 4b, Supplementary Fig. 13a).

**Fig 4.**
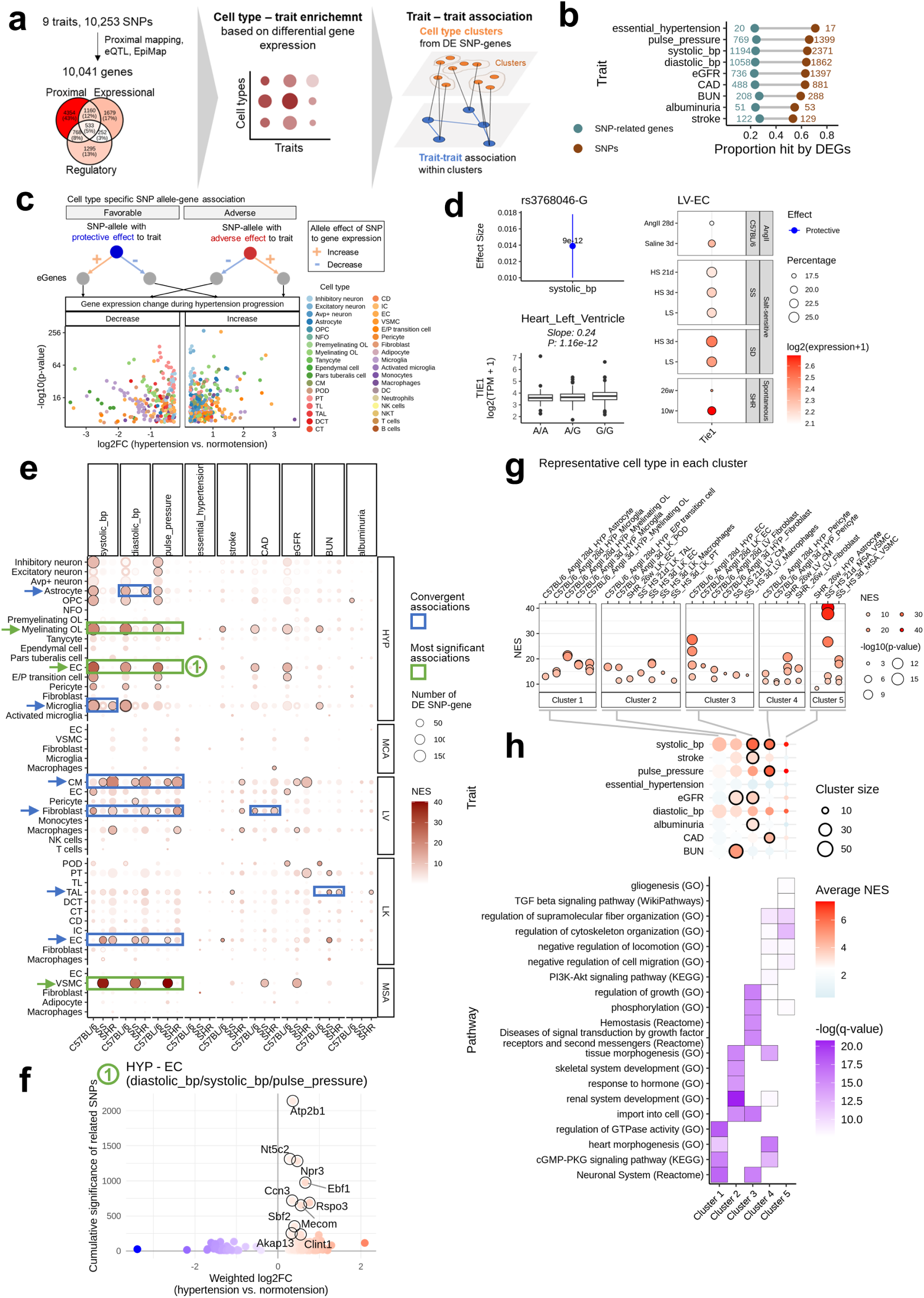
Cell type enrichment for genes involved in human genetic susceptibility to hypertension and end-organ damage. (a) Schematic of the analysis pipeline for integrating snRNA-seq data with human genome-wide association study (GWAS) data on hypertension and end-organ damage traits to perform cell type-trait enrichment and trait-trait association analysis. (b) Lollipop plot displaying the number and proportion of SNP-related genes identified as DEGs from our dataset, along with their corresponding SNPs. (c) Volcano plot displaying consistent relationships among allelic effect of SNPs on BP, allelic effect on eQTL gene expression in humans, and DEG in specific cell types in our animal models. (d) Relationship between SNP rs3768046-G and *TIE1* expression. The figure includes: the effect size (95% CI) of rs3768046-G on systolic BP (top left), box plots showing associations between rs3768046 genotypes and *TIE1* expression from the GTEx heart left ventricle dataset (bottom left), and the expression of *Tie1* in heart left ventricle ECs across conditions from our snRNA-seq data (right). In the box plot, the central line represents the median, the box edges indicate the interquartile range (IQR) from the 25th to the 75th percentile, and the whiskers extend to the minimum and maximum values within 1.5 times the IQR. Data points outside this range are shown as outliers. (e) Cell type-trait association plot across models based on the enrichment of SNP-related genes in DEGs within each cell type. Significant associations (BH-adjusted p-value < 0.05, permutation test) are highlighted with black borders. (f) Scatter plot showing the association of genes to multiple blood pressure traits via cumulative significance of related SNPs (y-axis) and their weighted expression change in hypothalamus endothelial cells during hypertension (x-axis). The top 10 genes with the highest cumulative significance are highlighted. (g) Top 10 cell type-trait associations in each cell type cluster, derived from the presence-absence matrix of differentially expressed SNP-related genes across various models and conditions. (h) Enrichment of traits in cell type clusters. Significant enrichments (p-value < 0.05, Wilcoxon test) are highlighted with black borders (top), along with pathway enrichment of differentially expressed SNP-related genes within each cluster.

To better understand hypertensive SNP allele-gene association at the cell type-specific level, we integrated allele-specific GWAS results, eQTL data from GTEx, and cell type-specific gene expression from our dataset. We classified SNP allele-gene pairs into two categories: adverse or favorable (Fig. 4c). In adverse SNP allele-gene pairs, the allele associated with higher BP or greater hypertensive end-organ damage (identified by GWAS) is linked to higher gene expression based on GTEx and the orthologous gene up-regulated during hypertension in our animal models, or lower gene expression in GTEx with the orthologous gene down-regulated in hypertensive animal models. Conversely, favorable SNP allele-gene pairs involve the allele associated with lower BP or less hypertensive end-organ damage, linked to higher gene expression in GTEx but with the orthologous gene down-regulated in hypertensive animal models or lower gene expression in GTEx but with the gene up-regulated in hypertensive animal models. In total, we identified 804 adverse and 780 favorable SNP allele-gene associations, each showing cell-type specificity in our snRNA-seq data (Supplementary Fig. 13b-g, Supplementary Table 6). For example, the systolic BP-protective variant rs3768046-G is linked to higher *TIE1* expression, a key endothelial receptor for vessel integrity, in GTEx, and Tie1 expression was notably lower in SS compared to SD, and significantly downregulated in hypertensive mice and SHR, specifically in left ventricle endothelial cells (Fig. 4d).

To partition the genetic susceptibility of BP changes and hypertensive end organ damage into transcriptional contribution of various cell types, we assessed cell type-trait associations by calculating the enrichment of SNP-related genes within the differential expression patterns of each cell type (Fig. 4e). This analysis revealed expected associations, such as MSA VSMCs and endothelial cells from the hypothalamus and kidney linked to multiple BP traits, and kidney thick ascending limbs linked to BUN. Additionally, we discovered novel associations, including myelinating oligodendrocytes, astrocytes, and microglia from the hypothalamus linked to multiple BP traits, and left ventricle cardiomyocytes and fibroblasts linked to BP. Several of these associations suggest the involvement of inflammatory responses in BP regulation and hypertensive end-organ damage. These findings highlight the underappreciated roles of several cell types in the central nervous system and the heart in BP regulation, potentially mediated by the coordinated regulation of genes related to GWAS-identified loci. To further explore these associations within key cell types, we focused on genetic hotspot genes linked to multiple SNPs, ranking them by cumulative significance from GWAS analysis (Fig. 4f, Supplementary Fig. 14a). A notable example is *NPR3*, which was upregulated after AngII treatment in mouse hypothalamus endothelial cells and associated with multiple SNPs, including rs1173771 (Supplementary Fig. 14b). The association between *NPR3* and rs1173771 was previously investigated, including in our study of iPSC-derived endothelial cells^21,22^.

We explored the connections among BP and end-organ-damage traits by analyzing cell type cluster co-association patterns and specific SNP-gene transcriptional relationships. Cell types were clustered based on the presence-absence matrix of differentially expressed SNP-related genes across various models and conditions (Supplementary Fig. 15a). These clusters exhibited co-association patterns with specific traits, such as kidney cell-major Cluster 2 linking BUN and eGFR, AngII-responsive EC-rich Cluster 3 linking stroke and systolic BP, and cardiomyocyte-dominated Cluster 4 linking CAD and BP traits (Fig. 4g and h). These co-associations were further supported by functional enrichment analyses of SNP-gene subsets within each cluster, identifying renal system development in Cluster 2 and heart morphogenesis and PI3K-Akt signaling in Cluster 4 (Fig. 4h). Additionally, we identified distinct SNP-gene expression relationships linking BP and end-organ damage, including: 1) BP SNPs related genes with expression changes in end-organ cell types; 2) end-organ-damage SNPs related genes with hypertension-responsive expression changes; and 3) SNPs associated with multiple traits, displaying cell type- or tissue-specific changes of their related genes (Supplementary Fig. 15b-d). These findings highlight substantial overlaps of cellular and molecular involvements across traits related to BP and hypertensive end-organ damage.

### rs28451064 orthologous noncoding genomic region influences pulse pressure in SS rats

As proof of principle, we sought to experimentally investigate select mechanisms revealed by the above analysis. To nominate a SNP-gene pair for further investigation, we applied a filtering strategy that integrates genetic, evolutionary, transcriptional, and technical feasibility criteria (Supplementary Fig. 16, Methods). Furthermore, we focused on SNPs located at least 100 kbp from any protein-coding gene to test if our dataset and approach could help to reveal the physiological role and mechanisms of action for these difficult-to-study SNPs. This approach led us to prioritize SNP rs28451064 for further investigation. GWAS have shown that rs28451064-A, compared with rs28451064-G, is associated with higher diastolic BP, smaller pulse pressure^6,23,24^ , and greater incidence of coronary artery disease^25–28^, myocardial infarction^28^, and angina pectoris^24,29^. A search using TOP-LD did not identify any SNPs that were in linkage disequilibrium (LD) with rs28451064 at r^2^>0.8. rs28451064 is approximately 143 kbp upstream of the transcription start site (TSS) of *KCNE2* and 148 kbp downstream from the shared TSS of *SLC5A3* and *MRPS6*. Other neighboring protein-coding genes include *SMIM11*, *SMIM34*, and *RCAN1* (Fig. 5a). In our single-nucleus dataset, *Slc5a3* was prominently expressed in the kidney cell types, and *Kcne2* and *Rcan1* were broadly expressed across cell types from various tissues (Supplementary Fig. 17). Although differential expression reached significance only in certain cell types, *Slc5a3* showed an overall lower expression in hypertensive strains after the onset of hypertension, in contrast to normotensive strains. And *Kcne2* showed an opposite expression pattern (Supplementary Fig. 16 and 17). Like most noncoding genomic loci associated with human traits, the effect of rs28451064 has not been examined with genome editing in vivo.

**Fig 5.**
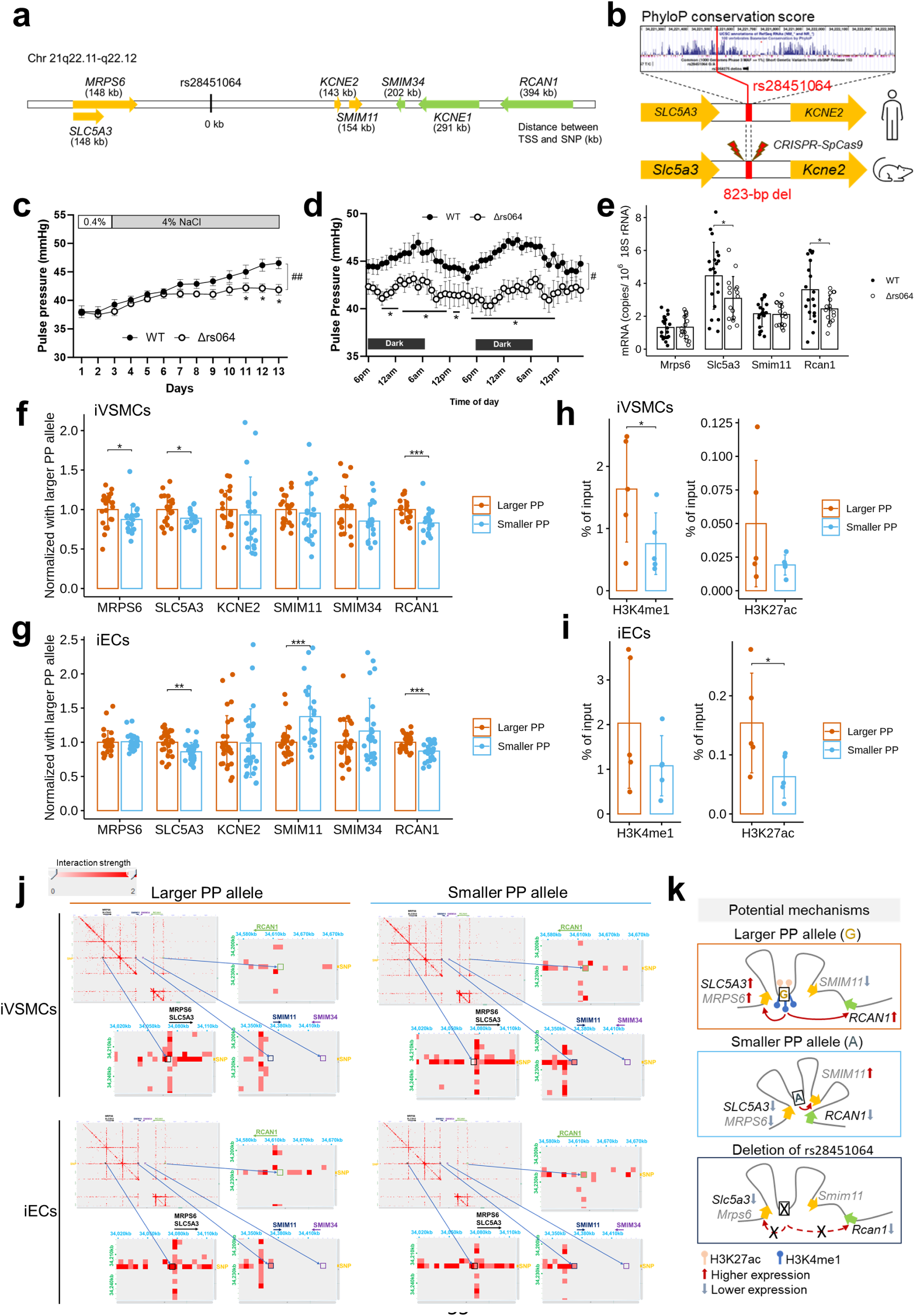
Noncoding SNP rs28451064 regulates *SLC5A3*, *RCAN1* and other local genes and influences blood pressure. (a) Genomic location of noncoding SNP rs28451064 and local genes. (b) Generation of the SS- Δrs28451064 rat. rs28451064 lies within semiconserved sequences between humans and rodents. A ∼800-bp noncoding segment in the SS rat genome containing the orthologous site for rs28451064 was deleted using CRISPR/SpCas9 to create the SS-Δrs28451064 rat. The deleted segment is more than 100 kbp from the transcription start site of the closest protein-coding gene. (c) Pulse pressure was significantly smaller in SS-Δrs28451064^−/-^ (Δrs064) rats compared to wild-type littermate SS rats (WT). N=11 WT and 14 Δrs064. ##, p<0.01 between WT and Δrs064; *, p<0.05 vs. WT; two-way repeated measure ANOVA followed by Holm-Sidak test. (d) Hourly pulse pressure on days 9-10 on a 4% NaCl diet. N=11 WT and 14 Δrs064. #, p<0.05 WT vs. Δrs064, two-way repeated measure ANOVA; *, p<0.05 vs. WT, Holm-Sidak test. (e) *Slc5a3* and *Rcan1* expression in mesenteric tissues was down-regulated in SS- Δrs28451064^−/-^ (Δrs064) rats. N=19 to 20 for WT and 16 for Δrs064. Data shown as mean ± s.e.m. *, p<0.05, unpaired t-test. (f) *SLC5A3*, *RCAN1* and *MRPS6* expression was lower in isogenic hiPSC-derived vascular smooth muscle cells (iVSMCs) with homozygous smaller PP allele of rs28451064 compared with the larger PP allele. N=21. *, p<0.05, unpaired t-test. (g) *SLC5A3* and *RCAN1* expression was lower, and *SMIM11* was higher, in isogenic hiPSC-derived endothelial cells (iECs) with homozygous smaller PP allele of rs28451064 compared with the larger PP allele. N=27. *, p<0.05, unpaired t-test. (h) H3K4me1 binding at the rs28451064 region was decreased in isogenic iVSMCs with homozygous smaller PP allele of rs28451064 compared with the larger PP allele. N=5. *, p<0.05, paired t-test. (i) H3K27ac binding at the rs28451064 region was decreased in isogenic iECs with homozygous smaller PP allele of rs28451064 compared with the larger PP allele. N=5. *, p<0.05, paired t-test. (j) Chromatin contact maps at rs28451064 locus and the surrounding region based on a region capture Micro-C analysis, were shown for iVSMCs and iECs with larger PP allele of rs28451064 (left)or smaller PP allele of rs28451064 (right). (k) Schematic showing the potential mechanisms of allelic effects of rs28451064 on the expression of *SLC5A3*, *RCAN1*, *MRPS6* and *SMIM11*. Black-colored genes showed changes common to both iEC and iVSMC, while grey-colored genes showed changes specific to either iECs or iVSMCs. The proximity of the SNP and genes indicates chromatin contacts. Curved red arrows indicate stimulatory effects of the SNP, which involve direct chromatin contact in some cases. Bar plots are shown as mean ± s.e.m.

We mapped the human rs28451064 to a noncoding genomic region in rat (Supplementary Fig. 18a). The local gene organization in the rat genome is similar to human. We used CRISPR/Cas9 to successfully delete an 823 bp noncoding segment in the SS rat genome encompassing the orthologous site for human rs28451064 (Fig. 5b, Supplementary Fig. 18b and c). The mutant strain was designated SS-Δrs28451064. Mean arterial pressure, systolic BP, diastolic BP, and heart rate were not significantly different between SS-Δrs28451064^−/-^ rats and wild-type (WT) SS littermates on the 0.4% or 4% NaCl diet (Supplementary Fig. 19). However, pulse pressure was significantly reduced in SS-Δrs28451064-/- rats by up to 6 mmHg over several days on the 4% NaCl diet (Fig. 5c). The reduction of pulse pressure in male rats appeared to be due to higher diastolic pressure (Supplementary Fig. 19b). Pulse pressure was lower in both light and dark phases of the day (Fig. 5d) and in both male and female SS-Δrs28451064^−/-^ rats compared with the WT SS rats (Supplementary Fig. 20). *Slc5a3* and *Rcan1* expression in the mesenteric tissue was downregulated in SS-Δrs28451064^−/-^ rats (Fig. 5e).

### Allelic effects of rs28451064 on local gene expression in human vascular cells

Next, we examined the allelic effect of rs28451064 on local gene expression in human vascular cells. We generated isogenic hiPSCs precisely edited to contain homozygous rs28451064-G (low DBP and larger pulse pressure allele) or rs28451064-A (high DBP and smaller pulse pressure allele) (Supplementary Fig. 21-23). The edited hiPSCs, three clones each, were differentiated to endothelial cells (iECs) and vascular smooth muscle cells (iVSMCs) (Supplementary Fig. 24). In both iVSMCs and iECs, the abundance of *SLC5A3* and *RCAN1* was significantly lower in cells with the smaller PP allele of rs28451064 compared those with the larger PP allele (Fig. 5f and g). The lower abundance of *MRPS6* in smaller PP allele was specific to iVSMCs, and higher abundance of *SMIM11* in smaller PP allele was specific to iECs.

In the genomic region containing the smaller PP allele of rs28451064, a reduction in H3K4me1 was detected in iVSMCs, and a reduction in H3K27ac was found in iECs, compared to the larger PP allele (Fig. 5h and i). Region-capture Micro-C analysis with probes targeting the rs28451064 region and the promoter regions of neighboring protein coding genes revealed a chromatin interaction between rs28451064 and the *RCAN1* promoter specific to the smaller PP allele in both iVSMCs and iECs (Fig. 5j). The interaction between rs28451064 and *SMIM11* promoter was specifically absent in iVSMCs with the larger PP allele. These findings suggest a consensus regulatory role of rs28451064 on *SLC5A3* and *RCAN1* expression through allelic-specific chromatin modification or chromatin interaction changes, alongside cell type-specific mechanisms regulating gene expression of *MRPS6* and *SMIM11* (Fig. 5k).

### rs1173771-NPR3 influences podocyte injury in hypertension

We recently reported that the rs1173771 noncoding haplotype influenced *NPR3* (natriuretic peptide receptor 3) expression in hiPSC-derived vascular cells and that salt-induced hypertension was attenuated and vasoreactivity was improved via an *NPR3*-dependent mechanism in SS rats with the rs1173771 orthologous noncoding genomic region deleted (SS-Δrs1173771LD^−/-^ rats)^22^. *NPR3* encodes natriuretic peptide receptor C (NPRC). The current snRNA-seq dataset showed a more prominent expression of *Npr3* in kidney podocytes than all other cell types (Supplementary Fig. 25). RNAScope analysis in SS rat kidneys confirmed expression of *Npr3* in podocytes (Supplementary Fig. 26).

These findings led us to hypothesize that rs1173771 and NRPC influenced podocyte function and the closely related albuminuria in hypertension. Despite significant attenuation of hypertension in SS- Δrs1173771LD^−/-^ rats fed the high-salt diet, albuminuria was not significantly different between SS- Δrs1173771LD^−/-^ rats and WT littermate SS rats (Supplementary Fig. 27a-27d). In SS-Δrs1173771LD^−/-^ rats, Npr3 expression in glomeruli and specifically podocytes was lower than in WT rats (Supplementary Fig. 27e, 27f). These findings suggest that the deletion of the rs1173771 haplotype orthologous region decreased NPRC expression in podocytes and increased the sensitivity of podocytes to hypertension-induced injury. In podocytes derived from hiPSCs (Supplementary Fig. 28a), treatment with tumor necrosis factor α or angiotensin II significantly increased the formation of stress fibers, which was attenuated by C-type natriuretic peptide (CNP), a ligand for NPRC (Supplementary Fig. 28b-S28d). The effect of CNP was blocked by AP811, an antagonist for NPRC (Supplementary Fig. 28b-S28d). Together, these findings indicate that NPRC protects podocytes from injury induced by hypertension and hypertension-relevant stressors. It suggests that, instead of the recognized role of renal NPRC in clearing natriuretic peptides, NPRC in podocytes under hypertensive conditions may mediate the signaling of CNP as NPRC does in blood vessels^30,31^.

## Discussion

The current study has provided an unprecedented view of the molecular landscape underlying the development of hypertension and hypertensive end-organ damage. The findings point to several conceptual advances for the understanding of these disorders, which may also be relevant to the understanding of other complex disorders. First, we found extensive, disease-associated molecular changes that were shared across tissues, models, or cell types. For example, approximately one third of DEGs were shared by two or more of the three models, and close to a quarter of the DEGs were shared by two or more of the five tissues. These findings highlight a greater degree of molecular coordination and commonality in the regulation of complex traits than generally recognized. The findings also illustrate the power of our multi-model, multi-tissue study design for understanding complex traits.

Second, we confirmed the known involvement of many genes and pathways in hypertension but also made several unexpected discoveries. Examples of the latter include the consensus elevated cellular communication within the hypothalamus and *Mecom*+ ECs in the kidney. The anatomical location of *Mecom*+ ECs in the medulla suggests that they may contribute to the regulation of renal medullary blood flow and tubular reabsorption, which are known to be important in BP regulation^32^. These findings point to new directions of future research and demonstrate the unique value of agnostic and systems-level analyses.

Third, we substantially elevated and expanded the utility of our large dataset from animal models by integration with human data, especially human GWAS data. We carried out this integration in innovative ways including considerations of clusters of traits and cell types as well as allelic effects, discovering novel, convergent connections between genetic susceptibility, disease traits, genes, and cell types. These approaches can be applied to integrate animal model data with human GWAS data for other complex traits.

One of the greatest challenges in human genetics is that most SNPs associated with complex traits are noncoding and many are located far from any protein-coding gene. Targeted, mechanistic, and physiological investigation of these SNPs is difficult and largely lacking. By integrating human GWAS data with rich animal model data, we developed a filtering process that effectively prioritized SNP-gene-disease relationships for targeted experimental investigation. Using genome editing in vivo and in vitro, we have shown that the BP-associated SNP rs28451064 may regulate the expression of *SLC5A3*, *RCAN1*, and other local genes, in part through long-range chromatin interactions, and influence BP. Lower or loss of *SLC5A3* or *RCAN1* expression is known to exacerbate vasoconstriction. *SLC5A3* encodes sodium/myo-inositol cotransporter 1 (SMIT1)^33^. Knockdown of SMIT1 increased the contraction of renal arteries induced by α1-adrenergic receptors agonist methoxamine and impaired the response to an activator of potassium channels Kv7.2–Kv7.5^34^. *RCAN1* (regulator of calcineurin 1) encodes calcipressin-1, which may inhibit or activate calcineurin or act through mechanisms independent of calcineurin. Deletion of *Rcan1* in vascular smooth muscle cells or endothelial cells in mice induces a hypercontractile phenotype, predisposing the mice to several hypertension-induced vascular abnormalities^35^.

Our studies of rs28451064 and its local genes as well as rs1173771 and podocyte NRPC illustrate how the rich data from the current omics study can be integrated with human data to drive novel, mechanistic studies of hypertension and hypertensive end-organ damage and bridge the critical gap between genetic discoveries and physiological understanding. Furthermore, our study of rs28451064 supports the notion that GWAS noncoding variants may have greater effects on associated traits in groups of humans with permissive genomic backgrounds and exposed to certain stressors than the minute effects reported by GWAS. The study also suggests that long-range chromatin interactions may play important roles not only in genetics and development but also in the regulation of physiological function.

The snRNA-seq analysis in the current study is limited in its representation of vascular tissues with just over 30,000 nuclei from middle cerebral artery and the 3^rd^ order mesenteric artery. We initially included arterial tissues from additional samples and treatment groups in our study. Unfortunately, the data from several samples did not meet our quality criteria to be included in the subsequent analysis. We suspect this was primarily due to the greater challenge in isolating high-quality nuclei from arteries than other tissues. It would be important to investigate vascular tissues in greater depth in future studies when the methods improve.

Sex difference is well-recognized in the development of hypertension and hypertensive end-organ damage^36,37^. The current study focused on male animals to ensure sufficient depth for the large-scale study. It would be highly valuable to perform similar studies in female animals in the future and compare the findings with our findings from the male animals. Indeed, several studies using single cell approaches to compare males and females in other research contexts have been highly informative^38,39^.

## Methods

### Treatment and phenotyping of C57BL/6 mice treated with Angiotensin II

All experimental procedures involving animals were approved by the Institutional Care and Use Committee of the Medical College of Wisconsin. The protocols for mice were approved by the Institutional Animal Care and Use Committee at the Medical College of Wisconsin (AUA1381). Male C57BL/6 mice were obtained from the Jackson Laboratory and were randomly assigned to one of three groups, saline-infused for 3 days, Angiotensin II (Ang II, Sigma-Aldrich, A9525)-infused for 3 days, and Ang II-infused for 28 days (n = 8 per group). Each mouse was implanted with a 28 Day (Alzet 1004; Durect Corporation) micro-osmotic pump subcutaneously containing saline or 490 ng/kg/min Ang II at 11 weeks of age as described previously^40^. Systolic BP was measured using tail-cuff plethysmograph (Visitech Systems BP-2000 Series II Blood Pressure Analysis System™, Apex, NC)^41^. BP was measured at baseline (before micro-osmotic pump implantation) and on the day before tissue collection. After BP measurement, each mouse was moved into a metabolic cage to collect 24h urine.

### Treatment and phenotyping of Dahl salt-sensitive rat, SD rat, SHR and WKY rat

The protocols for rats were approved by the Institutional Animal Care and Use Committee at the Medical College of Wisconsin (AUA206). Inbred Dahl salt-sensitive rats maintained at Medical College of Wisconsin (SS/JrHsdMcwi), SD rats obtained from ENVIGO, SHR and WKY rats obtained from Charles River were used in current study. Telemeter (model HD-S10, DSI, Harvard Bioscience, Inc.) was implanted in the right carotid artery for measuring arterial BP^42,43^. Rats were returned to the home cages with free access to water and chow after surgery. The mean of BP recorded from 9am to 12pm was used in the current study. After BP measurement, rats were moved into a metabolic cage to collect urines.

Dahl SS rats and SD rats were fed an AIN-76A diet containing 0.4% NaCl after weaning or upon arrival, respectively. The LS groups were maintained on the 0.4% NaCl until tissue collection when rats were 9-10 weeks of age (n=8 per group). The HS3d groups and HS21d group were fed an AIN-76A diet containing 4% NaCl for 3 days or 21 days, respectively, starting when rats were 9-10 weeks of age.

SHR and WKY rats were maintained on 5L79 diet. BP was recorded when rats were 10 weeks or 6 months of age. Mean BP of three days was used in the current study.

### Tissue collection – mouse

Plasma was collected through right ventricle puncture, aliquot and snap frozen. Kidneys were cut from the cross section. The upper and lower half of the left kidney and upper half of the right kidney were snap frozen. The front half of lower right kidney was frozen in O. C. T compound (Tissue-Tek, REF 4583). The back half of lower right kidney was stored in 10% formalin. Cardiac left ventricle was divided into 5 pieces. One piece was frozen in O. C. T compound. One piece was stored in 10% formalin. The remaining 3 pieces were snap frozen. Mesentery tissue, hypothalamus, and middle cerebral arteries were collected and snap frozen. Frozen tissues were stored at -80°C until analysis.

### Tissue collection – rat

At the termination, rats were anesthetized with a mixture of ketamine (75 mg/kg), xylazine (10 mg/kg), and acepromazine (2.5 mg/kg). Kidneys were flushed with 30 ml saline. To avoid affecting other organs, ascending aorta, mesenteric artery and abdominal aorta were tied before flushing. Next, both sides of kidneys were removed and processed in the same way as for mice. Third order of mesenteric arteries were dissected, and then snap frozen in liquid nitrogen, slowly frozen in O. C. T compound, or stored in 10% formalin. Cardiac left ventricle, hypothalamus and middle cerebral arteries were collected and processed in the same way as for mice.

### Morphological analyses

Cross-sectional area of cardiomyocytes, as an index of hypertrophy, was measured within transverse cardiac sections^44,45^. Wall thickness ratio of mesenteric artery was determined by artery media to inner lumen ratio^46,47^. Left ventricle and kidney sections were stained with Masson trichrome for fibrosis analysis. The positive-stained area was quantified as we described previously^40,47,48^. FIJI software (NIH; https://imagej.net) was used for the quantification.

### Urine analysis

Urinary albumin was quantified using a fluorescent assay utilizing albumin blue 580 dye (Biosynth). Urine creatinine was measured by an ACE Alera Autoanalyzer Clinical Chemistry System (Alfa Wassermann). The autoanalyzer utilizes the Jaffé Reaction for creatinine measurement. The measurements were performed by Biochemical Assay Core at Medical College of Wisconsin.

### Single-nucleus RNA-seq

The same type of -80°C frozen tissue from up to 4 animals was pooled together and ground into powder in liquid nitrogen. Nuclei were extracted using the ‘Frankenstein’ protocol for nuclei isolation from fresh and frozen tissue. Briefly, the ground tissue powder was resuspended in 4 ml ice-cold Nuclei EZ Lysis buffer (Sigma-Aldrich, St. Louis, MO) and incubated for 5 minutes on ice before being filtered through a 70 µm strainer mesh. The extracted nuclei were washed once with Nuclei Wash and Suspension Buffer. The nuclei were then stained with 7-AAD and subjected to sorting using a BD FACSMelody cell sorter (BD Biosciences, Franklin Lakes, NJ). Single-nucleus RNA-seq libraries were prepared using Chromium Single Cell 3’ Reagent Kits V3 (10x Genomics, Pleasanton, CA), following the manufacturer’s user guide. Up to 16,000 nuclei were loaded for each library. snRNA-seq libraries were sequenced using the Illumina NovaSeq sequencers to generate approximately 30 GB/100 million paired-end reads of data per library.

### Combined single-nucleus RNA-seq and ATAC-seq

Nuclei were isolated from frozen tissue pooled from up to 4 animals following the Nuclei Isolation from Complex Tissues protocol for Single Cell Multiome ATAC + Gene Expression Sequencing by 10x Genomics. The nuclei were then stained with 7-AAD and subjected to sorting using a BD FACSMelody cell sorter (BD Biosciences, Franklin Lakes, NJ). Up to 16,000 nuclei were used with the Chromium Next GEM Single Cell Multiome ATAC + Gene Expression kit (10x Genomics, Pleasanton, CA), following the user guide to create the libraries. snRNA-seq and snATAC-seq libraries were sequenced using Illumina NovaSeq sequencers to generate approximately 30 GB/100 million paired-end reads or 10 GB/33 million paired-end reads of data per library, respectively.

### Poly(A)-dependent RNA-seq and data analysis

Poly(A)-dependent RNA-seq experiment and data analysis were performed as described^49,50^.

### Preprocessing and quality control of snRNA-seq data

Raw fastq files were processed using CellRanger (v7.0.0)^51^ to generate feature matrices, with either the 10x Genomics reference genome assembly mm10 (refdata-gex-mm10-2020-A) or rn7, created via CellRanger mkref using mRatBN7.2 from Ensembl release 108. To minimize systematic background noise, CellBender (v0.2.0)^52^ was applied with the remove-background module. The parameters -- expected-cells and --total-droplets-included were set based on manual inspection of each sample’s CellRanger barcode rank plot. DoubletFinder R package (v2.0.3)^53^ was used to eliminate potential doublets, with the doublet rate set to max(0.3, #cells*8*1e-6). Further quality control steps included removing cells with fewer than 200 genes, more than 20,000 unique molecular identifiers (UMIs), or a mitochondrial gene content exceeding 20%.

### Preprocessing and quality control of snMultiome data

CellRanger-ARC (v2.0.2)^54^ was used to process raw fastq files, generating both feature and peak matrices. The reference genome assemblies used were either mm10 (refdata-cellranger-arc-mm10-2020-A-2.0.0) or rn7, created with cellranger-arc mkref and mRatBN7.2 from Ensembl release 108. Potential doublets were removed using DoubletFinder R package (v2.0.3)^53^. Cells were further excluded if they had fewer than 200 genes, more than 20,000 UMIs, a mitochondrial content greater than 20%, TSS enrichment <2, peak counts less than 400 or greater than 100,000.

### Clustering and annotation

Cells passing quality control were merged for each model-tissue, and subsequent analyses were performed using Seurat R package (v4.3.0.1)^55^. For snRNA-seq data, the gene count matrix was log-transformed and scaled, with mitochondrial percentage regressed out. We identified the top 2000 most variable genes using the FindVariableFeatures function, which were then used for principal component analysis (PCA). Batch effects were then corrected using Harmony R package (v1.2.0)^56^. Clustering was based on the top 30 principal components using the Louvain algorithm, with resolution adjusted from 0.1 to 1 in increments of 0.1. For snATAC-seq data, the peak matrix was normalized via Term Frequency-Inverse Document Frequency (TF-IDF) and processed using partial singular value decomposition (SVD). Batch correction was again performed with Harmony. For the snMultiome-seq dataset, we integrated the top 30 dimensions from snRNA-seq and the 2nd to 30th from snATAC-seq to construct a weighted nearest neighbor graph, and clustering was performed using the SLM algorithm, with resolution optimized between 0.1 and 1. The best resolution was selected based on Clustree (v0.5.1) analysis^57^.

To annotate identified clusters, differentially expressed genes (DEGs) between clusters were identified by Wilcoxon rank sum test using FindAllMarkers function (logfc.threshold = 0.25, min.pct = 0.25, only.pos = TRUE). Statistical significance was determined by adjusted p-value <0.05. Reference markers from PanglaoDB^58^ and CellMarker^59^ were used to annotate major cell types (Supplementary Fig.

7a, Supplementary Table 3). Endothelial cells were extracted, and subtype clustering was performed using the previously described pipeline within each model-tissue. Marker genes for each subcluster were cross-referenced across models to ensure consistent annotation (Supplementary Table 4). For immune cells, Cd45+ clusters were extracted from MCA, LV, and MSA samples and merged to create a reference project, following the previously described clustering pipeline. The *Cd45*+ RNA assay from the kidney sample was then integrated into this reference using FindTransferAnchors and MapQuery, resulting in a merged dataset from four tissues. This dataset was subsequently reclustered using previously described pipeline, and immune cell types were annotated using SingleR R package (v2.4.1)^60^ with the ImmGen reference^61^ (Supplementary Fig. 7b). Clusters with mixed marker expression were removed.

### Differential abundance analysis using Milo

To assess differential abundance of cell populations across different experimental conditions, miloR R package (v1.6.0)^5^ was applied separately for each strain-tissue. For each strain-tissue, neighborhood graphs were generated using the top 30 principal components from Harmony reduction. Neighborhoods were defined to include 20% of the graph vertices. Differential abundance testing was performed for the following comparisons: AngII-treated mice at 3 days versus saline-treated mice at 3 days, AngII-treated mice at 28 days versus saline-treated mice at 3 days, SS rats on a high-salt (HS) diet for 3 days versus low-salt (LS) diet, SS rats on a HS diet for 21 days versus LS diet, SD rats on a HS diet for 3 days versus LS diet, SHR rats at 26 weeks versus 10 weeks, and WKY rats at 26 weeks versus 10 weeks.

### Differential gene expression analysis

Differential gene expression analysis was processed within each cell type and tissue across comparison groups using the Wilcoxon rank sum test, as implemented in the Seurat FindMarkers function. The comparisons included: 1) treatment versus control within each strain: AngII-treated mice at 3 days versus saline-treated mice at 3 days, AngII-treated mice at 28 days versus saline-treated mice at 3 days, SS rats on a HS diet for 3 days versus LS diet, SS rats on HS diet for 21 days versus LS diet, SD rats on HS diet for 3 days versus LS diet, SHR rats at 26 weeks versus 10 weeks, and WKY rats at 26 weeks versus 10 weeks; and 2) hypertensive versus normotensive strains at baseline: SS rats on LS diet versus SD rats on LS diet, and SHR rats at 10 weeks versus WKY rats at 10 weeks. DEGs were considered significant if they met the criteria of a Bonferroni-corrected p-value less than 0.05 and an absolute log2 fold change greater than 0.5.

### Functional annotation based on DEGs

Pathway enrichment analysis for each cell type and tissue was performed using Metascape (https://metascape.org/)^62^, based on the DEGs generated in the previous step. This analysis integrated pathways from Gene Ontology (GO), KEGG, Reactome, and WikiPathways. Metascape evaluated pathway similarity based on shared genes, constructed a pathway network, and nominated representative pathways within the network. The significance threshold was set at a false discovery rate (FDR) of 0.05. For enriched pathways or GO terms derived from DEGs in treatment versus control comparisons, we focused on pathways specifically enriched in hypertensive strains by excluding those enriched in normotensive control strains, such as SD and WKY rats.

### Cell-cell communication analysis

Cell-cell communication analysis was conducted within each strain-treatment-tissue group using CellChat R package (v1.6.1)^63^. For each ligand-receptor pair, the communication probability (or strength) across cell types was summarized for each condition and then compared between relevant conditions to calculate the differential interaction strength. The comparisons included AngII-treated mice at 3 days versus saline-treated mice at 3 days, AngII-treated mice at 28 days versus saline-treated mice at 3 days, SS rats on a HS diet for 3 days versus a LS diet, SS rats on HS diet for 21 days versus LS diet, SD rats on HS diet for 3 days versus LS diet, SHR rats at 26 weeks versus 10 weeks, and WKY rats at 26 weeks versus 10 weeks.

### Transcription factor analysis

We applied pySCENIC (v0.12.1)^19^ to identify potential transcription factors (TFs) regulating gene expression in our snRNA-seq data. The workflow involved gene co-expression module inference, cis-regulatory motif enrichment, and regulon activity scoring in individual cells. For mouse data, we utilized the cisTarget databases (v1.24.0), generated using the SCENIC+ motif collection (2022) as a reference. For the rat snRNA-seq data, we generated a custom cisTarget database using the create_cistarget_databases tool (https://github.com/aertslab/create_cisTarget_databases). To adapt the motif information from the RcisTarget mouse database for rats, we mapped orthologous genes based on Ensembl annotations. Regulatory regions around gene start sites in the rat genome (mRatBN7.2, Ensembl Release 112) were extracted. These regions included sequences spanning 500 bp upstream and 100 bp downstream, 2 kbp upstream and 200 bp downstream, 8 kbp upstream and 1 kbp downstream, and 10 kbp upstream and downstream. The workflow for pySCENIC is the same as mouse.

### GTEx data analysis

To calculate the correlation between the neuronal connection score and *AVP* expression within the human hypothalamus, gene TPMs was accessed from GTEx Analysis V8^64^ and log2-transformed. The neuronal connection score was calculated using the ssGSEA method from the GSVA R package (v1.46.0)^65^, based on genes involved in the NCAM, NEGR, and NRXN pathways, including *NCAM1, NCAM2, FGFR1, L1CAM, NEGR1, NRXN1, NRXN2, NRXN3, NLGN1, NLGN2, and NLGN3.* Pearson correlation was applied to assess the relationship between the neuronal connection score and *AVP* expression.

### Kidney endothelial cell data analysis

The endothelial cell adaptive score was calculated on a cell-wise basis using the AddModuleScore function from the Seurat R package, based on the gene set shown in Figure 2b, including *Slit2, Slc4a4, Slc16a12, Ptprj, Nox4, Nedd4l, Lrp2, Col4a3, Atp1b1, Sdk1, Ptprd, Erbb4, Efna5, Cacnb4, Ank3, Pkhd1, Igf1r, and Arhgap24. Chrm3* was excluded due to its opposite expression trend compared to the other genes.

Single-cell RNA-seq data from human kidney samples were downloaded from the Kidney Precision Medicine Project (https://atlas.kpmp.org/), merged, and normalized using the variance-stabilizing transformation. Clusters were identified following the previously described pipeline. Cell labels were transferred from the original KPMP reference^20^. Endothelial cells (ECs) were extracted for further subtype clustering and annotation. DEGs specific to the *MECOM*+ EC subtype was identified by comparing them to all other ECs using the Seurat FindAllMarkers function. DEGs with a Bonferroni-corrected p-value < 0.05 and an absolute log2 fold change > 0.25 were used for pathway enrichment analysis with Metascape^62^.

### Stereo-seq data analysis

Raw sequencing files from Stereo-seq were processed using the SAW pipeline (v6.1.0)^66^. Spots were processed at a bin size of 50, and low-quality spots containing fewer than 200 genes per spot were filtered out for downstream analysis. To estimate cell type proportions in each spot, Cell2location (v0.1.4)^67^ was employed, using cell type annotations from our kidney snRNA-seq dataset. The number of cells per location was set to N_cells_per_location = 5 during estimation.

### Integrative analysis of snRNA-seq and snATAC-seq data using ArchR

ATAC-seq fragment data were further processed using the ArchR R package (v1.0.2)^68^. Peak calling was performed using MACS2^69^, and peak-gene linkage analysis was conducted using the addPeak2GeneLinks function, which integrates snRNA-seq gene expression data to link regulatory peaks with nearby genes at both the major cell type and EC subtype levels. Peaks associated with marker genes of each EC subtype, at both major EC and EC subtype levels, were then extracted for motif enrichment analysis.

Motif enrichment analysis was performed using a hypergeometric test after adding CisBP motif information^70^ using the addMotifAnnotations function and generating background peaks matched for genomic accessibility using the addBgdPeaks function. Motif deviation z-scores for each motif were calculated using the addDeviationsMatrix function. Cell type-specific motifs were identified by comparing z-scores in one cell type to those in all other cell types using the Wilcoxon rank sum test, followed by Benjamini-Hochberg (BH) adjustment for p-values. For *Mecom*+ EC-specific motifs, significant motifs were identified with an FDR < 0.05 and a z-score difference > 0.35 (*Mecom*+ ECs compared to other EC subtypes). These motifs were cross-referenced between the salt-sensitive and spontaneous hypertensive models to identify consistent *Mecom*+ EC-specific motifs.

To identify putative target genes for selected transcription factors (TFs), we followed the pipeline described by Ober-Reynolds et al^71^. Briefly, this approach correlates motif activity with gene expression, calculating a linkage score for each gene based on linked peaks containing specific TF motifs. For each candidate TF, genes showing strong correlations between expression and global motif activity (absolute Pearson correlation > 0.25) and with high linkage scores (> 80th percentile) were considered potential regulatory targets.

### BP SNP-related gene identification

GWAS summary statistics for BP regulation and hypertension end-organ-damage traits were downloaded from the GWAS Catalog (https://www.ebi.ac.uk/gwas/), including systolic BP, diastolic BP, essential hypertension, pulse pressure, stroke, coronary artery disease (CAD), estimated glomerular filtration rate (eGFR), albuminuria, and blood urea nitrogen (BUN). Only SNPs with a p-value < 5e-8 were retained for further analysis. BP SNP-related genes were identified using the following approaches: 1) proximal information, where the nearest protein-coding gene to each target SNP was considered; 2) cis-eQTL information, obtained from BP-relevant tissues in the GTEx Analysis V8^64^, including artery aorta, artery coronary, artery tibial, brain hypothalamus, heart left ventricle, and kidney cortex. Significant SNP-gene pairs associated eGenes (q-value ≤ 0.05) were retained, where variants with a nominal P value below the gene-level threshold were considered significant; 3) regulatory information, where gene-regulatory region linkages covering target SNPs were derived from EpiMap^72^ using BP-relevant tissues including brain, endothelial, heart, and kidney.

To account for potential loss of DEGs during cross-species integration due to orthologous mismatches, a relatively lenient DEG threshold was applied (Bonferroni-corrected p-value < 0.05 and absolute log2 fold change > 0.25) to increase statistical power for subsequent analyses.

To test whether BP SNP-related genes are more likely to be DEGs in our snRNA-seq data, we selected non-BP-relevant traits (including both physiological and disease traits) and calculated their SNP-associated genes using the previously described pipeline. The traits included birth weight, bone density, breast cancer, colorectal cancer, Alzheimer’s disease, Parkinson’s disease, multiple sclerosis, rheumatoid arthritis, asthma, psoriasis, and chronic obstructive pulmonary disease. To eliminate confounding effects from overlapping genes, any genes previously identified as BP SNP-related were excluded from the control gene list. The proportion of DEGs was then compared between BP-relevant genes and the control gene list using the Wilcoxon rank sum test. Comparisons were made across different DEG thresholds, including Bonferroni-corrected p-value < 0.05, and absolute log2 fold change > 0, 0.25, 0.5, or 1.

### Cell type-trait enrichment analysis

To evaluate the enrichment of BP SNP-related genes within cell type-specific DEGs, Fisher’s exact test was used to assess the overlap between BP SNP-related genes and DEGs, compared against a background set of cross-species expressed genes. To account for variability in DEG numbers across cell types, 1,000 permutation tests were performed by randomly selecting genes to replace SNP-related genes, followed by Fisher’s exact test on the permuted sets. A normalized enrichment score (NES) was then calculated as -log10(p-value) of the observed test divided by the mean of the -log10(p-values) from the shuffled gene sets.

### Cell type cluster and trait-trait association analysis

Cell types across all conditions were clustered based on the presence-absence matrix of differentially expressed SNP-related genes using Leiden’s clustering algorithm on a Jaccard similarity distance matrix. To eliminate the impact of data sparseness, consensus clustering was applied to 1,000 independent runs of Leiden’s clustering to generate robust cell type clusters. The top 10 cell types with the largest NES for any trait were selected to characterize the clusters. For each trait, the NES of cell types within a cluster was compared to the NES of all remaining cell types using the Wilcoxon rank sum test. Significant results (p-value < 0.05) were used to establish relationships between clusters and traits. The relationship between multiple traits were inferred based on their co-enrichment in the cluster. Functional annotation of each cell type cluster was performed by conducting pathway enrichment analysis using Metascape^62^ on all differentially expressed SNP-related genes in each cell type within the clusters.

### SNP-gene pair nomination

To identify and prioritize SNP-gene pairs with potential biological significance in BP regulation, we applied a series of filtration steps combining genetic, evolutionary, and transcriptional criteria. First, we filtered the dataset to include only SNPs associated with BP traits, including diastolic BP, systolic BP, and pulse pressure. We used TopLD^73^ to identify linkage disequilibrium (LD) regions, and selected SNPs located within LD regions containing a single SNP, ensuring that the genetic association was not confounded by closely linked variants. These SNPs were then lifted over to the mouse (mm10) and rat (rn7) genomes using rtracklayer (v1.58.0)^74^ to identify SNPs conserved across species, focusing on those successfully mapped in both species. For each remaining SNP-gene pair, a weighted fold change was calculated separately for hypertensive and normotensive strains. Specifically, log2 fold changes from the DEG analysis were scaled across tissues and strains, and a weighted fold change z-score was generated by adjusting for the log10(p-value). We considered only SNP-gene pairs where the gene exhibited opposite weighted fold change z-scores between hypertensive and normotensive strains. Additionally, we restricted the analysis to SNP-gene pairs where the gene was linked to a single filtered SNP, eliminating genes associated with multiple SNPs to reduce ambiguity. Finally, we focused on SNPs located more than 10,000 base pairs away from the transcription start site (TSS) of their associated genes, thereby emphasizing distal regulatory elements. These criteria were used to prioritize SNP-gene pairs that may play crucial roles in BP regulation but are also more challenging to study.

### Bulk RNA-seq data analysis

Raw RNA-seq reads were first preprocessed to remove low-quality bases and adapter sequences using Trim Galore (v0.6.8) (https://github.com/FelixKrueger/TrimGalore). The cleaned reads were aligned to the reference genome (mm10 or rn7) using HISAT2 (v2.1.0) with default parameters^75^. The resulting alignment files in BAM format were then processed for transcript assembly and quantification using StringTie (v2.2.1)^76^. Differential gene expression analysis was performed using DESeq2 (v1.38.3)^77^ using raw read counts obtained from StringTie. After DESeq2 normalization, differential expression analysis was carried out between comparison groups consistent with those used in the snRNA-seq analysis. Genes with an FDR < 0.05 were considered significantly differentially expressed.

### SNP editing in hiPSCs

The generation of isogenic iPSCs with homozygous BP-elevating or BP-lowering allele for the SNP rs28451064 was achieved using an efficient two-step approach as we described^22^. Briefly, two synthetic sgRNAs targeting genomic regions flanking the SNP rs28451064 (Supplementary Table 7) and spCas9-2NLS Nuclease protein were used to delete a segment of DNA around rs28451064. After 48 hrs of transfection, cells underwent clonal expansion. iPSC clones showing the intended deletion in quick extraction of DNA and PCR (Supplementary Table 8) were further characterized for the deletion by PCR and sanger sequencing. iPSC line with the deletion of the region around rs28451064 was tested for their genetic stability by molecular karyotyping using hPSC Genetic Analysis Kit (STEMCELL). For the second step of 2-step SNP editing, homology dependent knock-in approach was used with two gRNAs targeting, the junction of the rs28451064 region deletion site (Supplementary Table 7) and ssDNA donor fragment for smaller PP rs28451064-A or larger PP rs28451064-G alleles with approximately 100bp long homology arms at the 3’ and 5’ sides of the SNP rs28451064 (Supplementary Table 9). Isogenic iPSC lines with the reconstitution of homozygous smaller PP rs28451064-A and larger PP rs28451064-G alleles were selected, confirmed, and assessed for their pluripotency, differentiation potential, and genetic stability by molecular karyotyping using hPSC Genetic Analysis Kit and Giemsa-based karyotyping.

### Differentiation of hiPSCs to endothelial cells (iECs) and vascular smooth muscle cells (iVSMCs)

The differentiation was performed as we described (Supplementary Fig. 24a)^22,78,79^. Briefly, hiPSCs were treated with mTeSR™ plus medium with 10 µM Rock inhibitor Y-27632 (STEMCELL Technologies) for 24 h and then N2B27 medium (Life Technologies) plus 8 µM CHIR99021 (Selleck Chemicals) and 25 ng/ml BMP4 (PeproTech) for 3 days to generate mesoderm cells. For EC induction cells were further induced with StemPro-34 SFM medium (STEMCELL Technologies) supplemented with 200 ng/ml VEGF (PeproTech) and 2 µM forskolin (Abcam) for 2 days, purified with CD144 (VE-Cadherin) magnetic beads (Miltenyi Biotec) on day 6, and cultured in StemPro-34 SFM medium supplemented with 50 ng/ml VEGF for 5 more days before harvest. For VSMC induction, mesoderm cells were grown with N2B27 medium supplemented with 10 ng/ml PDGF-BB (PeproTech) and 2 ng/ml Activin A (PeproTech) for 2 days, and N2B27 supplemented with 2 ng/ml Activin A and 2 µg/ml Heparin (STEMCELL Technologies) for 5 days. VSMCs were enriched by removing CD144 + cells using CD144 magnetic beads.

### Real-time PCR

RNA extraction and quantitative real-time PCR analysis were performed as described^40^. Primer sequences are shown in Supplementary Table 10. Expression levels of mRNAs were normalized to the endogenous control 18S using ΔΔCt method.

### Region-capture Micro-C and data analysis

The assay and data analysis were performed as described with some modifications^22,80,81^. Briefly, Dovetail® Micro-C kit was used to create Micro-C libraries following manufacturer’s User Guide. Region capture Micro-C libraries were then generated with reagents from Dovetail® targeted enrichment panels. The panel of 80bp probes covers the rs28451064 genomic region and promoters of adjacent protein coding genes. The libraries were sequenced using the Illumina NovaSeq sequencer. The data was analyzed using Dovetail Micro-C data analysis pipeline and HiCcompare^82^.

### Chromatin immunoprecipitation (ChIP) and qPCR

ChIP was performed using a Magna ChIP™ A/G Chromatin Immunoprecipitation kit (Millipore) mostly following the vendor’s protocol. Briefly, differentiated iECs and iVSMCs were dissociated using Accutase (STEMCELL Technologies) and crosslinked with 1% formaldehyde for 10 min. Chromatins were sonicated for 55s using Covaris S2 following a sonication program optimized to generate fragments of 150–1000 bp. Chromatin fragments were immunoprecipitated with anti-H3K4me1 antibody (ab8895, abcam), anti-H3K27Ac antibody (MABE647, Millipore) or control IgG (ABIN101961, Antibodies Online). The pullout DNA was purified using spin columns provided with the Magna kit. Real-time PCR was performed to estimate the abundance of specific genomic segments, comparing the pullout DNA (output) to DNA samples used as the input for immunoprecipitation. Primer sequences are shown in Supplementary Table 8.

### Generation of SS-Δrs28451064^−/-^ rat

CRISPR/Cas9 methods were used to delete a noncoding genomic segment in the SS (SS/JrHsdMcwi) rat. The LiftOver^83^ tool at UCSC Genome Browser placed the non-coding syntenic region to the human rs28451064 at rn7 chr11: 31411381. spCas9 (QB3 MacroLab, University of California, Berkely) was mixed with sgRNAs and injected into fertilized SS/JrHsdMcwi strain rat embryos. A founder was identified harboring an 823-bp deletion (rn7 chr11:31,411,172-31,411,994) along with TGG insertion and was backcrossed to the parental strain to establish a breeding colony. This SS-Δrs28451064^−/-^ strain is registered as SS-Del(11p)4Mcwi (RGDID: 407987267). Heterozygous SS-Δrs28451064^+/-^ breeders were maintained on a purified 0.4% NaCl diet AIN-76A (Dyets, Inc).

### Telemetry measurement of blood pressure of SS-Δrs28451064^−/-^ rats

Continuous measurement of BP in conscious, freely moving rats was performed using radiotelemetry as described (Supplementary Fig. 1) ^43,84^. Briefly, a telemetry transmitter (model HD-S10, Data Systems International) was implanted in the right carotid artery when rats were 7 weeks of age. After one week of recovery, continuous BP monitoring was started. Following three days of stable baseline BP recording, the diet was switched to an AIN-76A diet containing 4% NaCl (high salt diet, HSD) for 14 days. Both male and female rats were studied.

### RNAScope

FFPE blocks from the kidney samples were sectioned into 5-μm sections directly mounted on Superfrost™ Plus Microscope Slides (Fisher). Slides were then processed following the recommended protocols by Advanced Cell Diagnostics for RNAscope™ Multiplex Fluorescent V2 Assay. Following processing, images were captured with fluorescence microscopy (BZ-X810, Keyence, IL) under a 40x objective.

### Differentiation of hiPSCs to podocytes

The hiPSCs were differentiated to podocytes using a protocol previously described^85^. Briefly, hiPSCs were dissociated with Accutase (STEMCELL Technologies) and seeded onto Matrigel (Corning)-coated p60 dishes at a density of ∼1 × 10^6^ cells/dish in mTeSR™ plus medium (STEMCELL Technologies) supplemented with 10 μM ROCK inhibitor Y-27632 (Selleckchem). The next day (day 0), posterior primitive streak differentiation was initiated by treating cells with 100 ng/mL Activin A (PeproTech) and 3 μM CHIR99021 (Selleckchem) in podocyte culture base media (BM): DMEM/F-12, GlutaMAX™ (Gibco), 1x B27 serum-free supplement (ThermoFisher), and 1% (v/v) of penicillin-streptomycin (Fisher). After a 48-hour incubation, medium was changed to intermediate mesoderm induction media (BM supplemented with 8 μM CHIR99021) and refreshed daily. On day 6, medium was changed to nephron progenitor differentiation medium: BM with 200 ng/mL FGF9 (PeproTech) and 0.180 USP/mL Heparin (Sigma Aldrich). After 2 days of daily medium change, day 7 cells were dissociated with Accutase and plated at a density of ∼5 × 10^4^ cells/cm^2^ in laminin-511 (PeproTech)-coated plates with podocyte induction media: BM supplemented with 100 ng/mL BMP7 (PeproTech), 100 ng/mL Activin A, 50 ng/mL VEGF (PeproTech), 3 μM CHIR99021, and 0.1 μM all-trans retinoic acid (STEMCELL Technologies). Media was substituted daily until podocyte maturation on day 12, where hiPSC-podocytes would be used for subsequent experiments.

### Cytoskeleton staining

Fluorescent staining of the actin cytoskeleton in iPSC-podocytes were performed with Alexa Fluor™ 488 Phalloidin (ThermoFisher) according to the manufacturer’s protocol. Briefly, cells were fixed on glass coverslips with 4% freshly made paraformaldehyde in DPBS. Following fixation, cells were permeabilized for 10 min in 1% Triton X-100/PBS, washed with PBS, and stained with Alexa Fluor 488-conjugated phalloidin according to the manufacturer’s protocol. One drop of SlowFade™ Glass Soft-set Antifade Mountant with DAPI (Invitrogen) was placed on each slide before mounting a coverslip. Images were captured with fluorescence microscopy (BZ-X810, Keyence, IL) under a 20x objective. The different staining patterns of phalloidin were grouped into 4 different classes as previously described^86,87^. Briefly, type A: > 90% cell area filled with stress fiber cables; type B: at least 2 thick cables running under nucleus, and the rest of cell area filled with fine fibers; type C: no thick cables, but some fibers present; type D: no visible stress fibers in the central area of the cell. Approximately 50 cells per well and 8 wells per treatment group were analyzed.

### Statistical analysis and data visualization

R (v4.2.2)^88^ was used for statistical analysis. R package ggplot2 (v3.5.0)^89^ was used for data visualization.

## AUTHOR CONTRIBUTIONS

QQ developed the data analytical strategy and analyzed and interpreted the data. YL led the omics assays. HX performed phenotyping and tissue collection. RP performed hiPSC and molecular experiments. LH performed podocyte experiments. JL contributed to study design, phenotyping, and tissue collection. PL contributed to the conception of the study and data analysis and interpretation. BT contributed to omics assays. VK and JH contributed to data analysis. KU contributed to phenotyping and tissue collection. MG and MAVA contributed to the development of the mutant rat. ASG, AWC, SR contributed to data interpretation. AMG led the development of the mutant rat and contributed to data interpretation. ML conceived and designed the study. QQ drafted the manuscript. HX, RP, LH, and ML contributed to the drafting of the manuscript. All authors edited and approved the manuscript.

## DATA AVAILABILITY

All data generated in this study, including cell type-specific gene expression changes, are accessible on the accompanying web portal (https://viz.datascience.arizona.edu/content/2d089f1d-2904-4888-b86f-70ca2fe7a297/), providing a resource for the research community.

