## Supplementary information for "A single-cell map of hypertension"

[1]Department of Physiology, University of Arizona College of Medicine – Tucson, Tucson, AZ, USA; [2]Molecular Systems Medicine Initiative, University of Arizona Health Sciences, Tucson, AZ, USA; [3]Department of Physiology, Medical College of Wisconsin, Milwaukee, WI, USA; [4]Division of Nephrology, Department of Medicine, University of Rochester, Rochester, NY, USA; [5]The Jackson Laboratory, Bar Harbor, ME, USA; [6]Versiti Blood Research Institute, Milwaukee, WI, USA; [7]Department of Pediatrics, Division of Hematology, Oncology, and Transplantation, Medical College of Wisconsin, Milwaukee, WI, USA; [8]Department of Cell Biology, Neurobiology, and Anatomy, Medical College of Wisconsin, Milwaukee, WI, USA.

*co-first authors.

[#]co-corresponding authors:

Qiongzi Qiu,

Mingyu Liang,

**Supplementary figure legends**

**Figure S1. Hypertension in Angiotensin II treated mice, Dahl salt sensitive rats on a high salt diet and SHR of 10 and 26 weeks old.** (a) Systolic blood pressure (SBP) of C57BL/6 mice treated with saline or angiotensin II (AngII) 490ng/kg/min. (b) Mean arterial pressure (MAP) of Sprague-Dawley (SD) rats or Dahl salt sensitive (SS) rats on a 0.4% NaCl baseline diet or 4% NaCl high salt (HS) diet. (c) MAP of spontaneously hypertensive rats (SHR) and age matched Wistar Kyoto normotensive rats (WKY). n = 8 per group. *, $p < 0.05$, **, $p < 0.01$, or ***, $p < 0.001$ by t-test.

**Figure S2. Cardiomyocyte hypertrophy induced by Angiotensin II treatment in C57BL/6 mice and in spontaneously hypertensive rat model.** (a) Quantification of cross-sectional area of cardiomyocytes and (b) representative left ventricle sections, magnification, ×200 in C57BL/6 mice treated with saline or AngII 490ng/kg/min. (c) Quantification of cross-sectional area of cardiomyocytes and (d) representative left ventricle sections, magnification, ×200 in SHR and age matched WKY. n = 8 per group. **, $p < 0.01$, ***, $p < 0.001$ by one-way ANOVA.

**Figure S3. Fibrosis area in left ventricle is increased in Angiotensin II treated C57BL/6 mice and Dahl salt sensitive rats fed a 4% NaCl diet.** (a) Quantification by Masson Trichrome staining and (b) representative heart sections, magnification, ×200 in C57BL/6 mice treated with saline or AngII 490ng/kg/min. (c) Quantification by Masson Trichrome staining and (d) representative heart sections, magnification, ×200 in SD rats or SS rats on a 0.4% or 4% NaCl diet. n = 8 per group. *, $p < 0.05$ by one-way ANOVA.

**Figure S4. Wall thickness ratio of 3$^{rd}$ order mesenteric artery is increased in spontaneously hypertensive rats.** (a) Wall thickness ratio of 3$^{rd}$ order of mesenteric artery determined by artery media to inner lumen ratio and (b) representative cross-sectional mesenteric artery, magnification, ×200 in in SHR and age matched WKY. (c) Quantification of wall thickness ratio of 3$^{rd}$ order of mesenteric artery and (d) representative cross-sectional mesenteric artery, magnification, ×200 in SD rats or SS rats on a 0.4% or 4% NaCl diet. n = 8 per group. ***, $p < 0.001$ by one-way ANOVA.

**Figure S5. Kidney fibrosis is increased in Dahl salt sensitive rats and spontaneously hypertensive rats.** (a) Quantification by Masson Trichrome staining and (b) representative kidney sections illustrating renal cortex fibrosis, magnification, ×200 in SD rats and SS rats fed a 0.4% NaCl (LS) or 4% NaCl (HS) diet. (c) Quantification by Masson Trichrome staining and (d) representative kidney sections of outer medulla (OM), magnification, ×200. (e) Quantification by Masson Trichrome staining and (f) representative kidney sections illustrating renal cortex fibrosis, magnification, ×200 in SHR and age WKY normotensive rats. (g) Quantification by Masson Trichrome staining and (h) representative kidney sections of outer medulla (OM), magnification, ×200. n = 7 or 8 per group. *, $p < 0.05$, **, $p < 0.01$, ***, $p < 0.001$ by t-test.

**Figure S6. Urinary albumin in Angiotensin II treated mice, Dahl salt sensitive rats on a high salt diet and SHR of 10- and 26-week-old.** Urinary albumin to creatinine ratio (UACR) in (a) C57BL/6 mice treated with saline or AngII 490ng/kg/min, (b) SD rats or SS rats on a 0.4% NaCl baseline diet or 4% NaCl high salt (HS) diet, and (c) SHR and age matched WKY. n = 8 per group. *, $p < 0.05$, ***, $p < 0.001$ by t-test.

**Figure S7. Cell type annotation.** (a) Dot plot displaying the expression levels of selected marker genes across identified cell types in multiple models and tissues. (b) Heatmap of corresponding scores between the final annotation of integrated immune clusters in each model and the ImmGen cell type reference, with largest scores across reference cell types highlighted by asterisks. The AngII graph includes data from AngII-treated C57BL/6 mice. The salt-sensitive graph includes data from SS rats and SD rats on high-salt diets. The spontaneous graph includes data from SHR rats and WKY rats of different ages. Tissue abbreviations: HYP (hypothalamus), MCA (middle cerebral artery), LV (left ventricle), LK (left kidney), and MSA (3rd mesenteric artery).

**Figure S8. Stacked bar plot displaying the relative proportions of each major cell type across tissues and strains under different conditions.**

**Figure S9. Comparative analysis of gene expression and overlapping differential expressed genes (DEG) across hypertension models and studies.** (a) UMAP plots showing *Cd74* expression changes across conditions in three models. (b) Proportion of DEGs identified in one or more than one tissues and strains that overlap with human blood pressure-relevant genes reported in two studies (Keaton JM et al.[1] and Mishra MK et al.[2]).

**Figure S10. Differential gene expression and pathway enrichment across cell types in hypothalamus.** (a) Dot plot representing log2 fold changes of genes selected from Fig. 2a, across all cell types in each comparison group that yielded significant results. (b) Top up- and down-regulated genes in astrocytes, oligodendrocyte precursor cells (OPCs), and activated microglia in SS rats on high-salt diets. (c) Expression changes of selected genes specifically in microglia of SS rats on high-salt diets. (d) Dot plot showing the number of enriched pathways corresponding to cell-type-specific DEGs in the hypothalamus across comparison groups, grouped by occurrence across models.

**Figure S11. Circle plots illustrating ligand-receptor interactions from pathways in Fig. 2e in SS rats on high-salt diet and SHR rat of different ages, with edge width representing communication strength.**

**Figure S12. snMultiome-seq based analysis of marker gene expression, motif enrichment and TF-gene interactions in kidney endothelial cell subtypes.** (a) Dot plot showing marker gene expression in LK EC subtypes. (b) Top 10 motifs enriched by peaks associated with marker genes of each EC subtype in salt-sensitive (left) and spontaneous (right) models. (c) Dot plot showing TF motifs specifically enriched in *Mecom*+ endothelial cells (ECs) in both the salt-sensitive (x-axis) and spontaneous (y-axis) models, as measured by the motif activity deviation z-score difference between *Mecom*+ ECs and the remaining ECs. Horizontal and vertical dashed lines indicate a z-score difference of 0.3. TF motifs with a z-score difference greater than 0.3 in both models are highlighted in red. (d) Prioritization of gene targets for *Hoxb4* in salt-sensitive model. The x-axis represents the Pearson correlation between TF motif activity and integrated gene expression across endothelial cells. The y-axis shows the TF linkage score, calculated as the sum of scaled motif scores for all linked peaks based on peak-to-gene link correlation. The color of the points reflects the hypergeometric enrichment of the TF motif in linked peaks for each gene. The horizontal dashed line marks the 80th percentile of the linkage score, while the vertical dashed lines represent motif-to-gene expression correlations of -0.25 and 0.25, respectively.

**Figure S13. Overlap of DEGs and SNP-related genes across traits and SNP-eQTL associations in BP-relevant traits.** (a) Proportion of overlap between DEGs and SNP-related genes across traits, grouped by BP-relevant and BP-irrelevant traits, with DEGs categorized by different standards in each column. (b-g) Relationship between selected SNPs and eQTL gene expression: rs7302981-G with *CERS5* (b), rs841216-G with *MGRN1* (c), rs7255-C with *LDAH* (d), rs622076-A with *C4A* (e), rs2279499-A with *ST7L* (f), and rs460105-T with *VPS9D1* (g). Each panel includes: the effect size (95% CI) of the specific SNP allele on BP-relevant traits (left), box plots showing the association between SNP genotypes and gene expression from GTEx data in a relevant tissue (middle), and the expression of the gene in specific cell types in the corresponding tissue under different conditions from our snRNA-seq data (right).

**Figure S14. Association of genes with blood pressure traits.** (a) Scatter plot showing the association of genes with multiple blood pressure traits. The y-axis represents the cumulative significance of related SNPs, while the x-axis shows the weighted expression change of these genes in selected cell types during hypertension. The top 10 genes with the highest cumulative significance are highlighted. Cell types were selected based on Fig. 4e. (b) Expression levels of the top 10 genes with the highest cumulative significance in hypothalamus endothelial cells across conditions in the AngII model (left), alongside their top 5 associated SNPs (right).

**Figure S15. SNP-related gene based cell type cluster identification and SNP-cell type cluster association.** (a) Consensus similarity heatmap of cell types across strains, conditions, and tissues, based on a presence-absence matrix of differentially expressed SNP-related genes across various models and conditions. (b-d) SNP-cell type cluster associations via linked genes supported by multiple lines of evidence: rs179993 in Cluster 4 (b), rs11838776 in Cluster 4 (c), and rs77924615 in Cluster 2 (d). Each panel includes: GWAS association of the SNP with all BP-relevant traits (left), SNP-gene association supported by different evidence (middle), and gene expression changes across multiple cell types within the specific cluster (right).

**Figure S16. Pipeline for prioritizing BP traits-associated SNP-gene pairs (left). Scatter plot showing the expression changes of filtered genes in hypertensive and normotensive strains using weighted fold change, categorized by the trait associated with the corresponding SNP (right).**

**Figure S17. Expression of genes neighboring rs28451064**. (a) Dot plot showing the expression levels of selected genes across cell types in each tissue (y axis) and treatment in each model (x axis). (b) Differential expression results of selected genes across treatments in each model, based Wilcoxon test. Only genes with a Bonferroni-adjusted p-value less than 0.05 were shown, and highlighted with diamond shapes.

**Figure S18. Comparative analysis and generation of a rat model for rs28451064.** (a) Comparative map alignment between human and rat. The genes and order around rs28451064 are syntenic. (b) CRISPR-SpCas9 sgRNA targets flanking a ~800-bp region in rat semiconserved at the sequence level with the human rs28451064 locus. An 823-bp deletion in chromosome 11 was generated. (Note: The corresponding LiftOver coordinates from rn7 to rn6 were used to show vertebrate sequence conservation in this 823-bp region.) (c) Sanger sequence confirmation of the 823-bp deletion.

**Figure S19. Deletion of the rs28451064 orthologous region does not change mean arterial pressure (MAP), systolic blood pressure (SBP), diastolic blood pressure (DBP), or heart rate in SS rats.** (a) Data with male and female rats combined. N=12 and 14. (b) Male rats. N= 5 and 5. (c) Female rats. N=7 and 9.

**Figure S20. Deletion of the rs28451064 orthologous region decreases pulse pressure in both male and female SS rats on a 4% NaCl diet.** (a) Male rats. N= 5 and 5. (b) Female rats. N=7 and 9. #, $p<0.05$ for WT vs. delta-rs064; *, $p<0.05$ vs. WT; two-way repeated measure ANOVA followed by Holm-Sidak test.

**Figure S21. Generation of isogenic hiPSCs containing homozygous rs28451064-G (low DBP allele, or larger PP allele) or rs28451064-A (high DBP allele, or smaller PP allele).** (a) Schematic of the two-step genome editing process. (b) Deletion of DNA segment containing rs28451064 locus. 39b – original iPSC cell line, X32 - SNP rs28451064 deleted cell line. (c) Reconstitution of SNP rs28451064 locus containing either homozygous larger PP allele or smaller PP allele. 39b – original iPSC cell line, X8 - rs28451064 deleted cell line, y series – reconstituted rs28451064 larger PP allele cell line, z series – reconstituted rs28451064 smaller PP cell line.

**Figure S22. Sanger sequencing confirmation of hiPSC editing.** (a) Sanger sequencing confirming deletion of a 120 bp region containing rs28451064. (b) Sanger sequencing confirming reconstitution of either homozygous larger PP (top row, three clones) or smaller PP (bottom row, three clones) allele of rs28451064. Larger PP allele cell lines – y8, y11 and y14. Smaller PP allele cell lines – z2, z11 and z14.

**Figure S23. Quality assessment of isogenic hiPSCs containing homozygous rs28451064-G (larger PP allele) or rs28451064-A (smaller PP allele).** (a) Proliferation assay. Equal number of cells were seeded on day 0 in 6cm dish for reconstituted clones. Cells were collected on day 4 and total cell number for reconstituted iPSCs were compared with original iPSC cell line, 39b. Larger PP allele cell lines – y8, y11 and y14. Smaller PP allele cell lines – z2, z11 and z14. (b) Mycoplasma contamination screening for reconstituted iPSCs. x22 is another mycoplasma negative iPSC cell line. Original iPSC cell line - 39b. (c) Representative karyograms for three Large PP (y8, y11, y14) and three smaller PP (z2, z11 and z14) clones. Chromosomes of 20 proliferating cells were counted except for clone Z8 where 10 proliferating cells were counted and fully analyzed using G-banding. Three cells were karyotyped. These cells had a modal number of 46 chromosomes. The sex chromosome constitution indicates normal female. No consistent abnormalities were observed in the chromosomal number or banding patterns.

**Figure S24. Differentiation of edited hiPSCs to endothelial cells (iECs) and vascular smooth muscle cells (iVSMCs).** (a) Differentiation protocol adapted from Patsch et al., Nat Cell Biol, 2015[3]. (b) Representative DIC images of iEC and iVSMC differentiation on day 1, day 4, day 6 and day 11. Scale bar – 200 µm. (c) Expression of EC marker CD31 and VSMC marker ACTA2 in iECs and iVSMCs. N = 66 for iEC and N = 72 for iVSMC.

**Figure S25. Dot plot showing the expression levels of *Npr3* across cell types in each tissue (y axis) and treatment in each model (x axis) in our snRNA-seq data.**

**Figure S26. *Npr3* is expressed in podocytes.** Representative images from multiplex RNAScope showing the spatial distribution and co-localization of *Nphs1* (red; a podocyte marker) and *Npr3* (green) transcripts in kidney sections from SS rats. The top left panel displays *Npr3* transcripts, the middle left panel shows *Nphs1* transcripts, and the bottom left panel presents a merged image indicating areas of co-localization (yellow). The large right panel provides a zoomed-in view, where white arrowheads highlight regions of strong co-localization between *Nphs1* and *Npr3*, demonstrating the expression of *Npr3* in podocytes. Scale bars = 100 µm.

**Figure S27. Deletion of the rs1173771 haplotype orthologous region decreased NPR3 expression in podocytes and increased the sensitivity of SS rats to hypertension-induced albuminuria.** (a) Diagram depicting the rat genomic region orthologous to the human rs1173771 linkage disequilibrium (LD) region. A 30.4 kbp region orthologous to the 17.4 kbp human rs1173771 LD region was deleted from the genome of Dahl salt sensitive (SS) rats to generate SS-Δrs1173771LD rats[4]. The transcription start site of the closest protein-coding gene, *NPR3*, is approximately 126 kbp from the haplotype region in human genome and ~76 kbp in rat genome. (b) The increase of mean arterial pressure (ΔMAP) after diet switch to 4% NaCl, relative to baseline MAP on a 0.4% NaCl diet, was significantly attenuated in male SS-Δrs1173771LD rats (Δ771LD) compared with wild type (WT) littermate SS rats. The graph was re-plotted from data reported in Xue, et al, bioRxiv 2024[4]. N = 8 per group. #, $p < 0.05$ by two-way RM ANOVA, *, $p < 0.05$ by Holm-Sidak test. (c) Urinary albumin to creatinine ratio (UACR) in SS-Δrs1173771LD$^{-/-}$ rats was not different than WT littermates on the 0.4% NaCl baseline diet or 4% NaCl high salt (HS) diet. N = 8. (d) Changes of UACR from the baseline 0.4% NaCl diet to HS for 14 days were not different between SS-Δrs1173771LD$^{-/-}$ rats and WT littermates. N = 8. (e) *Npr3* expression in isolated glomeruli was decreased in SS-Δrs1173771LD$^{-/-}$ rats, based on qPCR. N = 8; *, $p < 0.05$, unpaired t-test. (f) *Npr3* expression in podocytes was decreased in SS-Δrs1173771LD$^{-/-}$ rats, based on RNAScope. Co-localization of *Nphs1*, a podocyte-specific marker, with *Npr3* was used to assess the expression of *Npr3* in podocytes. Co-localization analysis was performed with ImageJ using split-channel thresholded binary masks and image calculator "multiply" function to locate only pixels present in both channels. 4 rats per group and 18 glomeruli per rat were examined. *** $p < 0.001$, unpaired t-test.

**Figure S28. C-type natriuretic peptide (CNP), acting through natriuretic peptide receptor C (NPRC), protects podocytes.** (a) hiPSC-derived podocytes express podocyte marker *NPHS1*. The expression of *NPHS1*, normalized to 18S rRNA, is elevated by approximately 200-fold in hiPSC-derived podocytes compared to undifferentiated hiPSCs. ***, $p < 0.001$, unpaired t-test. (b) Representative images of different types of phalloidin staining patterns observed in hiPSC-derived podocytes. Cells were treated with TNFα or AngII alone and in combination with CNP or CNP + AP811. Phalloidin staining was used to assess actin cytoskeleton reorganization of the podocytes in response to the treatments. White arrows highlight representative stress fibers. Scale bars = 20 µm. Type A: >90% of cell area filled with thick stress fibers. Type C: no thick cables, but some fibers present. (c) Quantification of phalloidin staining in hiPSC-derived podocytes following a 24-hour treatment of 20 ng/mL TNFα with two additional interventions. TNFα significantly increased the proportion of Type A cells and decreased the proportion of Type C cells. Co-treatment with 100 nM CNP attenuated the effect of TNFα. The addition of 100 nM AP811, an antagonist of NPRC, reversed the protective effect of CNP. N = 6-8. * = $p < 0.05$, ** = $p < 0.01$, **** = $p < 0.0001$, two-way ANOVA followed by Holm-Sidak test. (d) AngII treatment (1 µM) produced a similar increase in stress fibers, with CNP co-treatment reducing stress fibers and AP811 reversing that effect. N = 6-8. ** = $p < 0.01$, **** = $p < 0.0001$, two-way ANOVA followed by Holm-Sidak test.

**Supplementary tables (submitted as an Excel file)**

Supplementary Table 1: Summary of animal samples used in snRNA-seq, snMultiome-seq, Stereo-seq, and bulk mRNA-seq.

Supplementary Table 2: QC metrics for snRNA-seq and snMultiome-seq.

Supplementary Table 3: Major cell type annotations across datasets.

Supplementary Table 4: EC subtype annotations across datasets.

Supplementary Table 5: Cell type-specific DEGs identified in all treatment-control comparisons across strains.

Supplementary Table 6: Consistent relationships among allelic effect of SNPs on BP, allelic effect on eQTL gene expression in humans, and DEG in specific cell types in our animal models.

Supplementary Table 7: Sequences of guide RNAs for rs28451064 editing.

Supplementary Table 8: PCR and qPCR primer sequences for hiPSC and hiPSC derived samples.

Supplementary Table 9: ssDNA donor fragment for rs28451064 alleles.

Supplementary Table 10: qPCR primer sequences for rat samples.

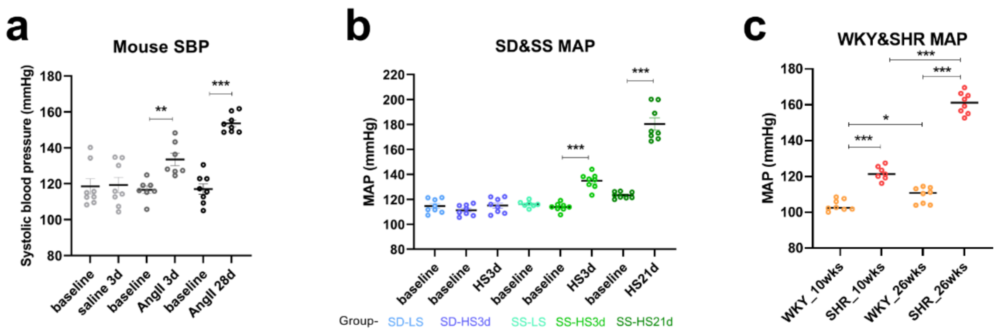


**Figure S1. Hypertension in Angiotensin II treated mice, Dahl salt sensitive rats on a high salt diet and SHR of 10 and 26 weeks old.** (a) Systolic blood pressure (SBP) of C57BL/6 mice treated with saline or angiotensin II (AngII) 490ng/kg/min. (b) Mean arterial pressure (MAP) of Sprague-Dawley (SD) rats or Dahl salt sensitive (SS) rats on a 0.4% NaCl baseline diet or 4% NaCl high salt (HS) diet. (c) MAP of spontaneously hypertensive rats (SHR) and age matched Wistar Kyoto normotensive rats (WKY). n = 8 per group. *, p < 0.05, **, p < 0.01, or ***, p < 0.001 by t-test.

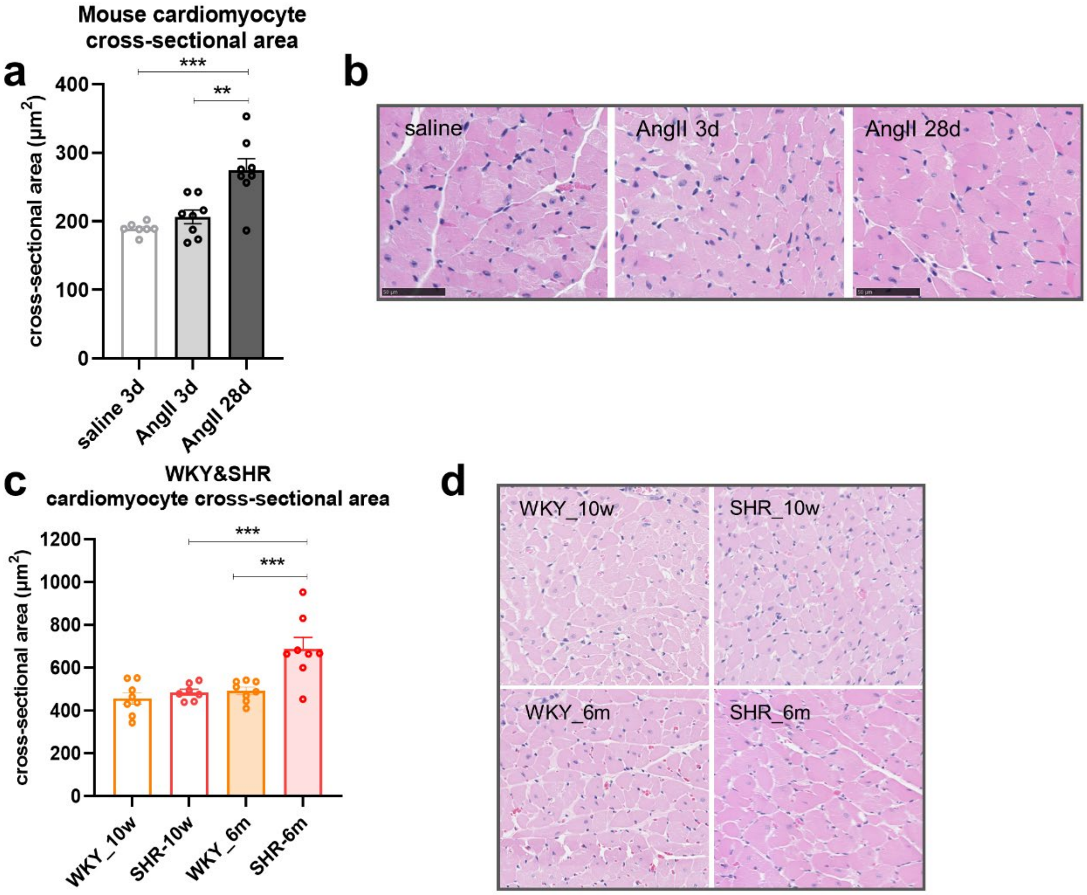


**Figure S2. Cardiomyocyte hypertrophy induced by Angiotensin II treatment in C57BL/6 mice and in spontaneously hypertensive rat model.** (a) Quantification of cross-sectional area of cardiomyocytes and (b) representative left ventricle sections, magnification, ×200 in C57BL/6 mice treated with saline or AngII 490ng/kg/min. (c) Quantification of cross-sectional area of cardiomyocytes and (d) representative left ventricle sections, magnification, ×200 in SHR and age matched WKY. n = 8 per group. **, $p < 0.01$, ***, $p < 0.001$ by one-way ANOVA.

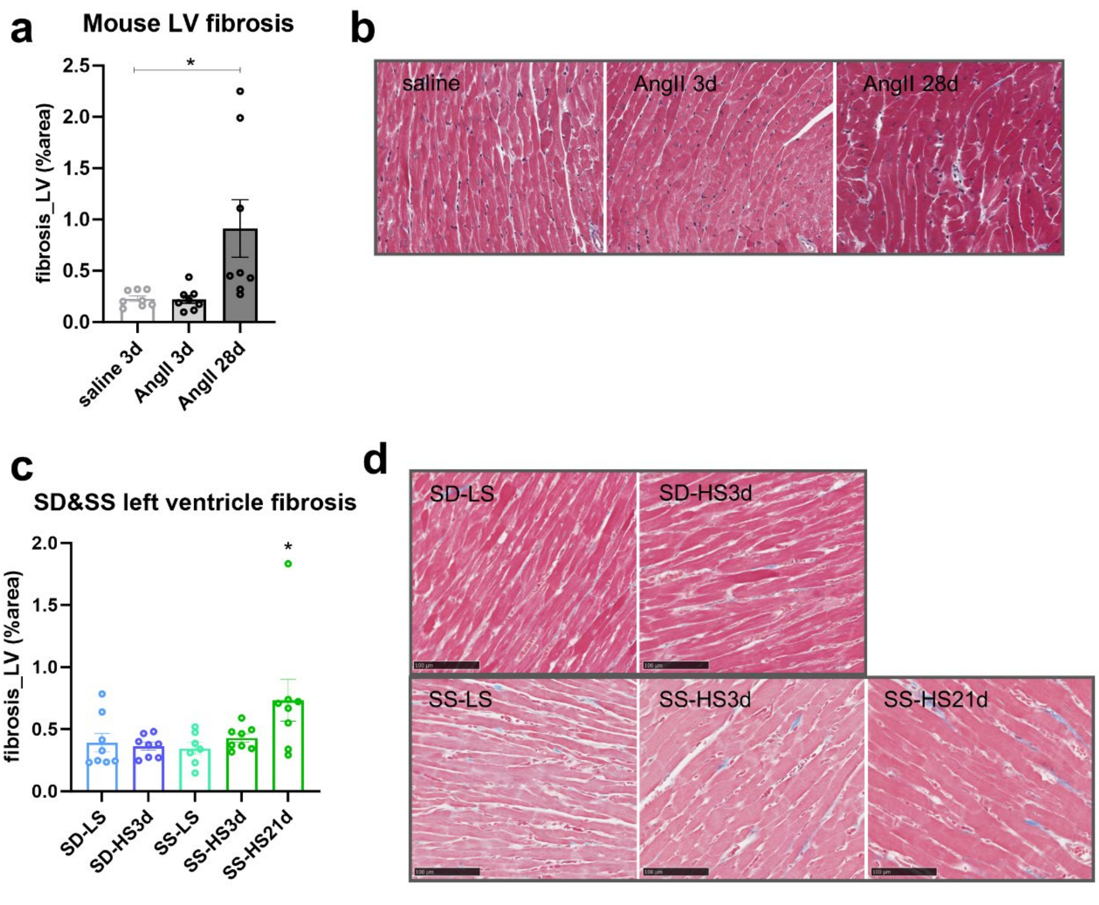


**Figure S3. Fibrosis area in left ventricle is increased in Angiotensin II treated C57BL/6 mice and Dahl salt sensitive rats fed a 4% NaCl diet.** (a) Quantification by Masson Trichrome staining and (b) representative heart sections, magnification, ×200 in C57BL/6 mice treated with saline or AngII 490ng/kg/min. (c) Quantification by Masson Trichrome staining and (d) representative heart sections, magnification, ×200 in SD rats or SS rats on a 0.4% or 4% NaCl diet. n = 8 per group. *, $p < 0.05$ by one-way ANOVA.

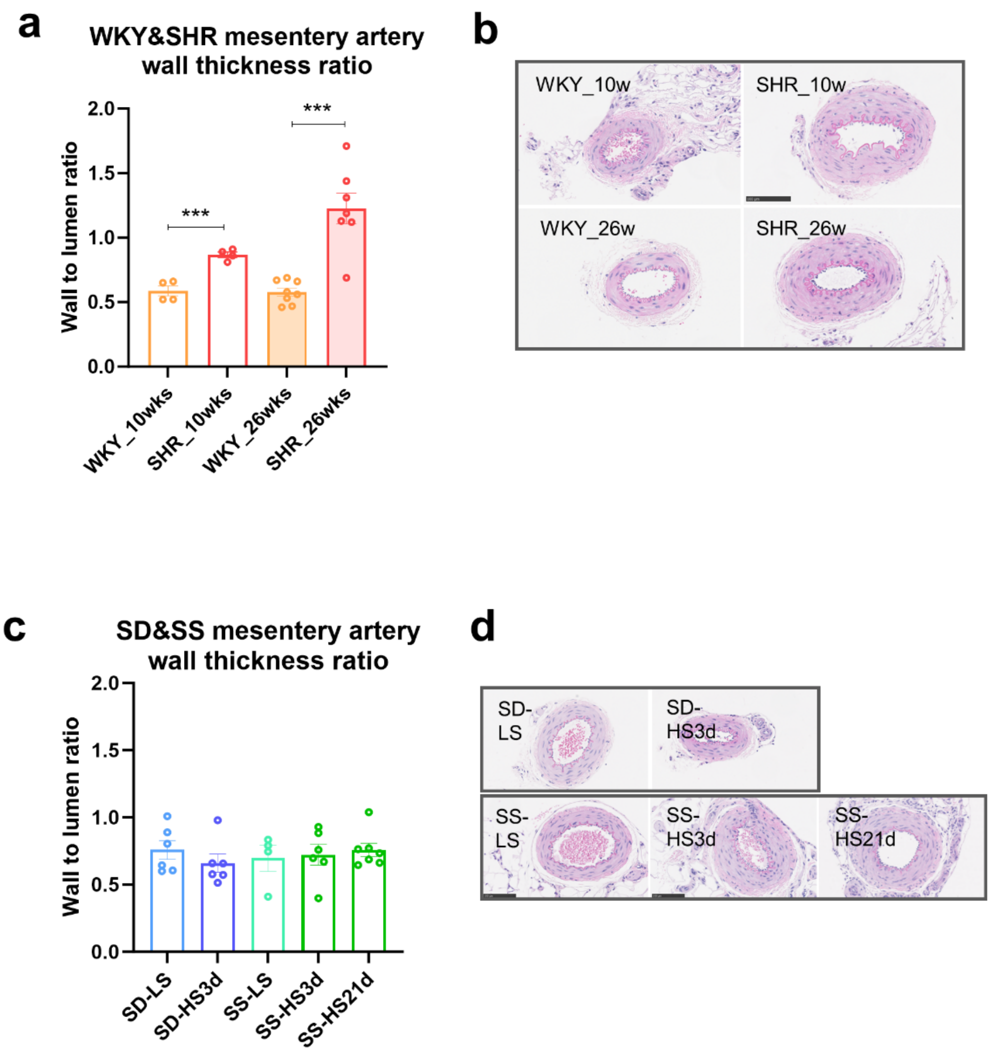


**Figure S4. Wall thickness ratio of 3rd order mesenteric artery is increased in spontaneously hypertensive rats.** (a) Wall thickness ratio of 3rd order of mesenteric artery determined by artery media to inner lumen ratio and (b) representative cross-sectional mesenteric artery, magnification, ×200 in in SHR and age matched WKY. (c) Quantification of wall thickness ratio of 3rd order of mesenteric artery and (d) representative cross-sectional mesenteric artery, magnification, ×200 in SD rats or SS rats on a 0.4% or 4% NaCl diet. n = 8 per group. ***, $p < 0.001$ by one-way ANOVA.

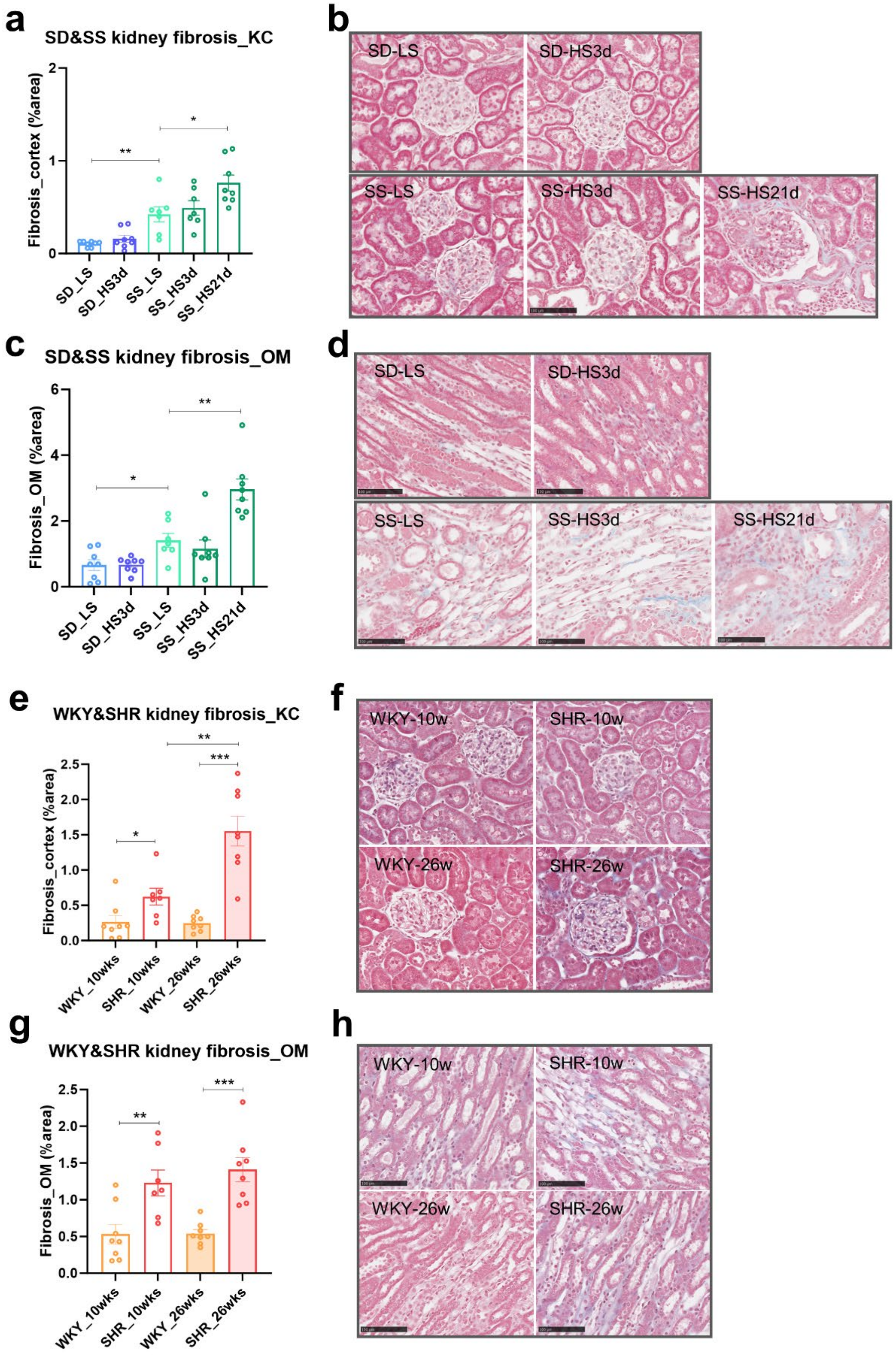


**Figure S5. Kidney fibrosis is increased in Dahl salt sensitive rats and spontaneously hypertensive rats.**

(a) Quantification by Masson Trichrome staining and (b) representative kidney sections illustrating renal cortex fibrosis, magnification, ×200 in SD rats and SS rats fed a 0.4% NaCl (LS) or 4% NaCl (HS) diet. (c) Quantification by Masson Trichrome staining and (d) representative kidney sections of outer medulla (OM), magnification, ×200. (e) Quantification by Masson Trichrome staining and (f) representative kidney sections illustrating renal cortex fibrosis, magnification, ×200 in SHR and age WKY normotensive rats. (g) Quantification by Masson Trichrome staining and (h) representative kidney sections of outer medulla (OM), magnification, ×200. n = 7 or 8 per group. *, $p < 0.05$, **, $p < 0.01$, ***, $p < 0.001$ by t-test.

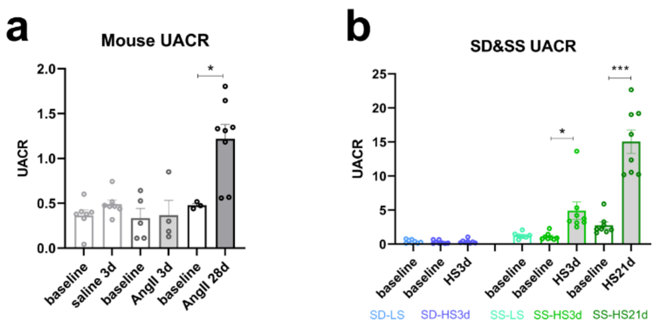


**Figure S6. Urinary albumin in Angiotensin II treated mice, Dahl salt sensitive rats on a high salt diet and SHR of 10- and 26-week-old.** Urinary albumin to creatinine ratio (UACR) in (a) C57BL/6 mice treated with saline or AngII 490ng/kg/min, (b) SD rats or SS rats on a 0.4% NaCl baseline diet or 4% NaCl high salt (HS) diet, and (c) SHR and age matched WKY. n = 8 per group. *, $p < 0.05$, ***, $p < 0.001$ by t-test.

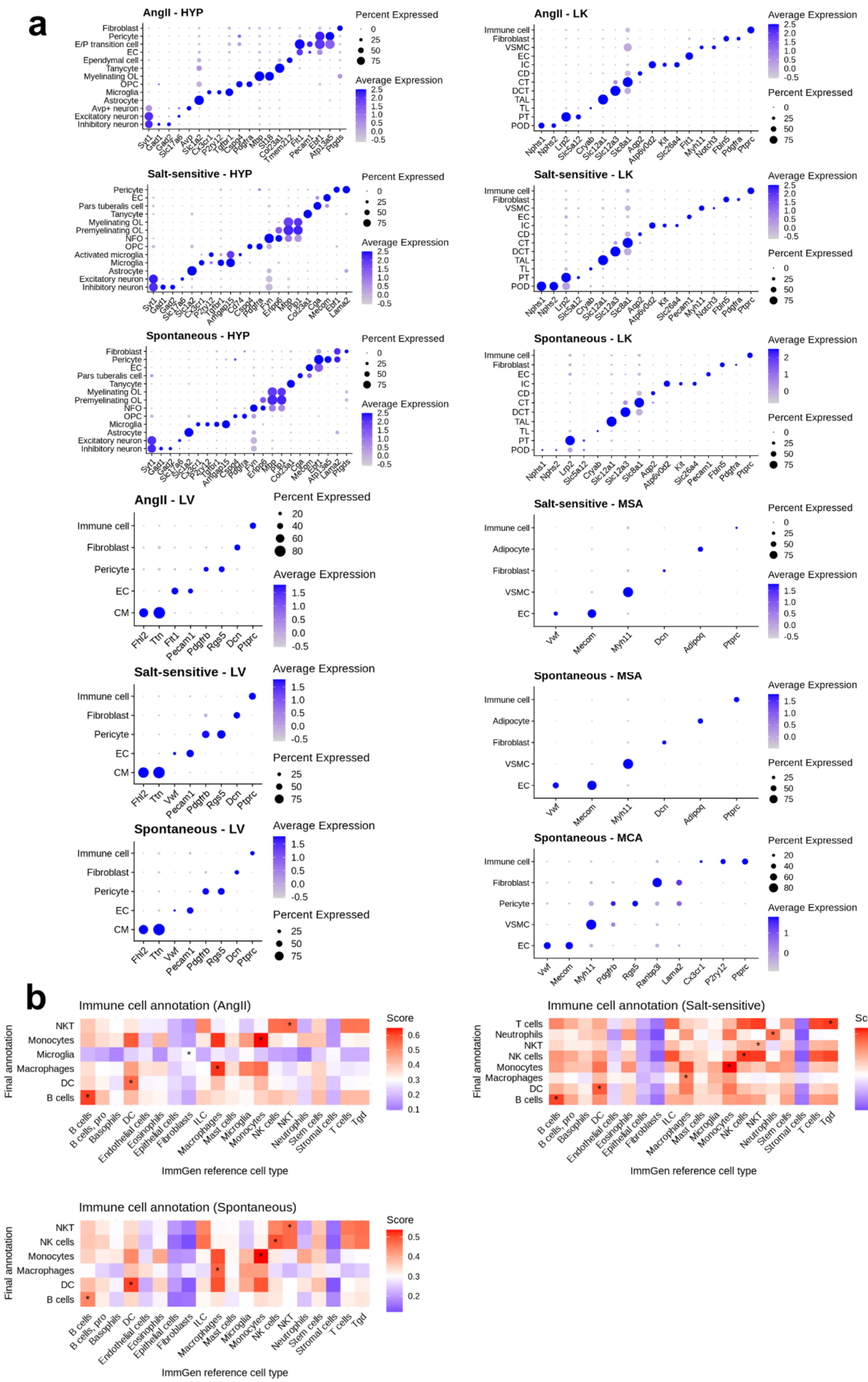
a
AngII - HYP
Percent Expressed
Average Expression
AngII - LK
Salt-sensitive - HYP
Salt-sensitive - LK
Spontaneous - HYP
Spontaneous - LK
AngII - LV
Salt-sensitive - MSA
Salt-sensitive - LV
Spontaneous - MSA
Spontaneous - LV
Spontaneous - MCA
Immune cell
Fibroblast
Pericyte
EC
CM
VSMC
Adipocyte
Fhl2
Ttn
Vwf
Pecam1
Pdgfrb
Rgs5
Dcn
Ptprc
Mecom
Myh11
Adipoq
b
Immune cell annotation (AngII)
Immune cell annotation (Salt-sensitive)
Immune cell annotation (Spontaneous)
Score
Final annotation
ImmGen reference cell type
NKT
Monocytes
Microglia
Macrophages
DC
B cells
T cells
Neutrophils
NK cells

**Figure S7. Cell type annotation.** (a) Dot plot displaying the expression levels of selected marker genes across identified cell types in multiple models and tissues. (b) Heatmap of corresponding scores between the final annotation of integrated immune clusters in each model and the ImmGen cell type reference, with largest scores across reference cell types highlighted by asterisks. The AngII graph includes data from AngII-treated C57BL/6 mice. The salt-sensitive graph includes data from SS rats and SD rats on high-salt diets. The spontaneous graph includes data from SHR rats and WKY rats of different ages. Tissue abbreviations: HYP (hypothalamus), MCA (middle cerebral artery), LV (left ventricle), LK (left kidney), and MSA (3rd mesenteric artery).

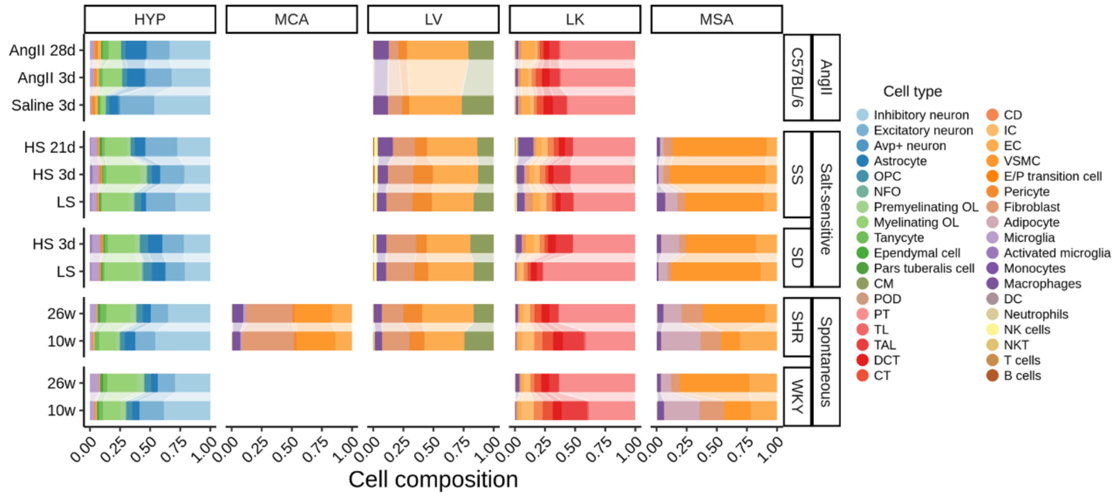


**Figure S8. Stacked bar plot displaying the relative proportions of each major cell type across tissues and strains under different conditions.**

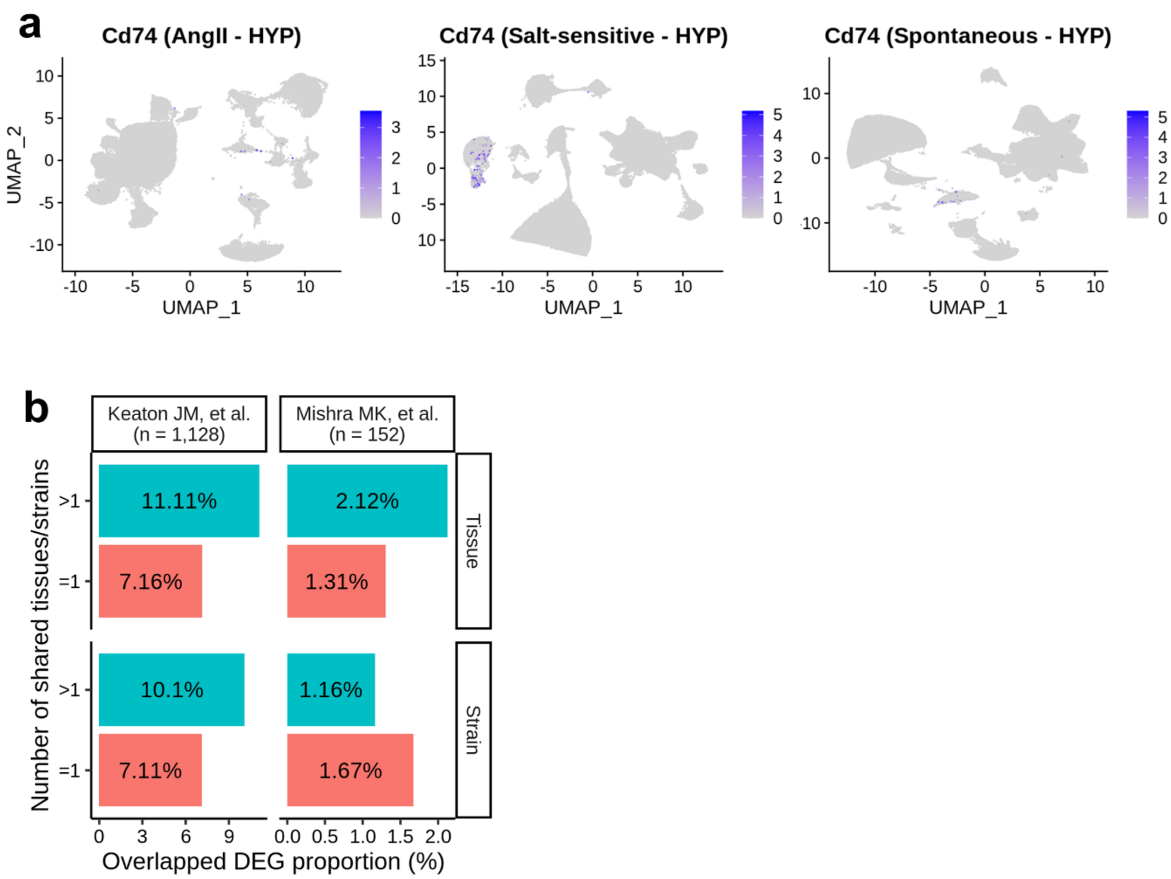


**Figure S9. Comparative analysis of gene expression and overlapping differential expressed genes (DEG) across hypertension models and studies.** (a) UMAP plots showing *Cd74* expression changes across conditions in three models. (b) Proportion of DEGs identified in one or more than one tissues and strains that overlap with human blood pressure-relevant genes reported in two studies (Keaton JM et al.[1] and Mishra MK et al.[2]).

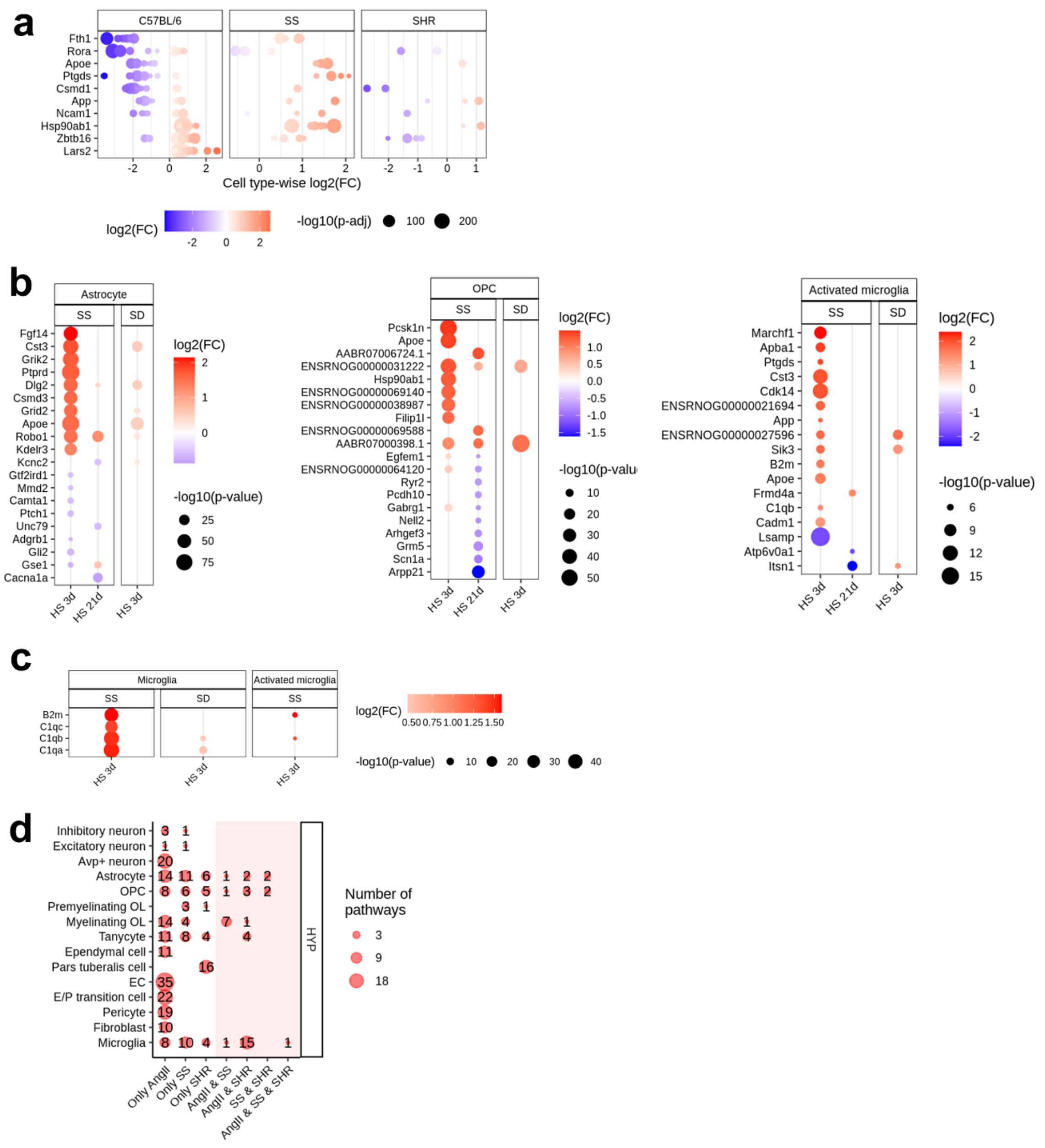


**Figure S10. Differential gene expression and pathway enrichment across cell types in hypothalamus.** (a) Dot plot representing log2 fold changes of genes selected from Fig. 2a, across all cell types in each comparison group that yielded significant results. (b) Top up- and down-regulated genes in astrocytes, oligodendrocyte precursor cells (OPCs), and activated microglia in SS rats on high-salt diets. (c) Expression changes of selected genes specifically in microglia of SS rats on high-salt diets. (d) Dot plot showing the number of enriched pathways corresponding to cell-type-specific DEGs in the hypothalamus across comparison groups, grouped by occurrence across models.

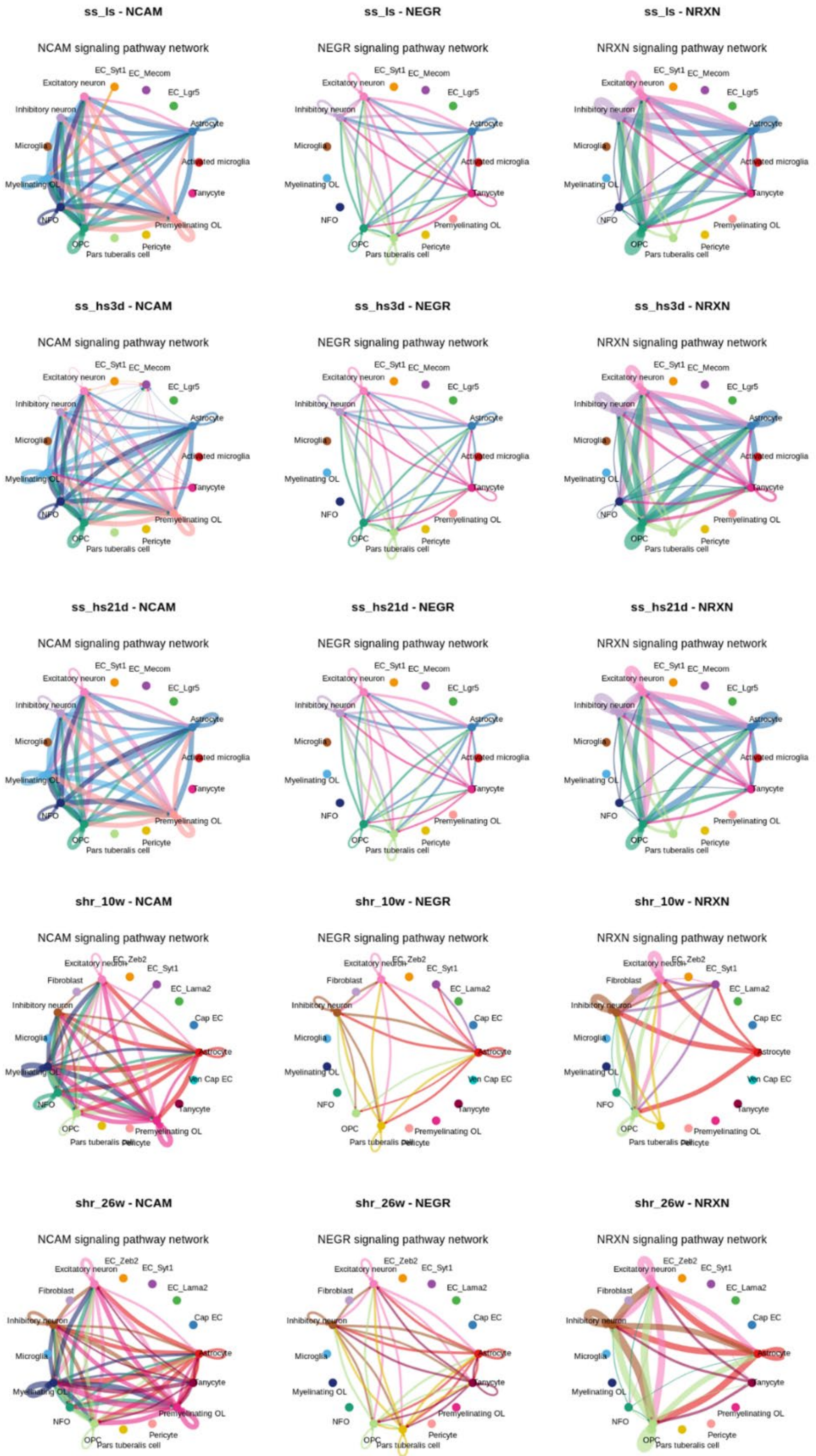


**Figure S11. Circle plots illustrating ligand-receptor interactions from pathways in Fig. 2e in SS rats on high-salt diet and SHR rat of different ages, with edge width representing communication strength.**

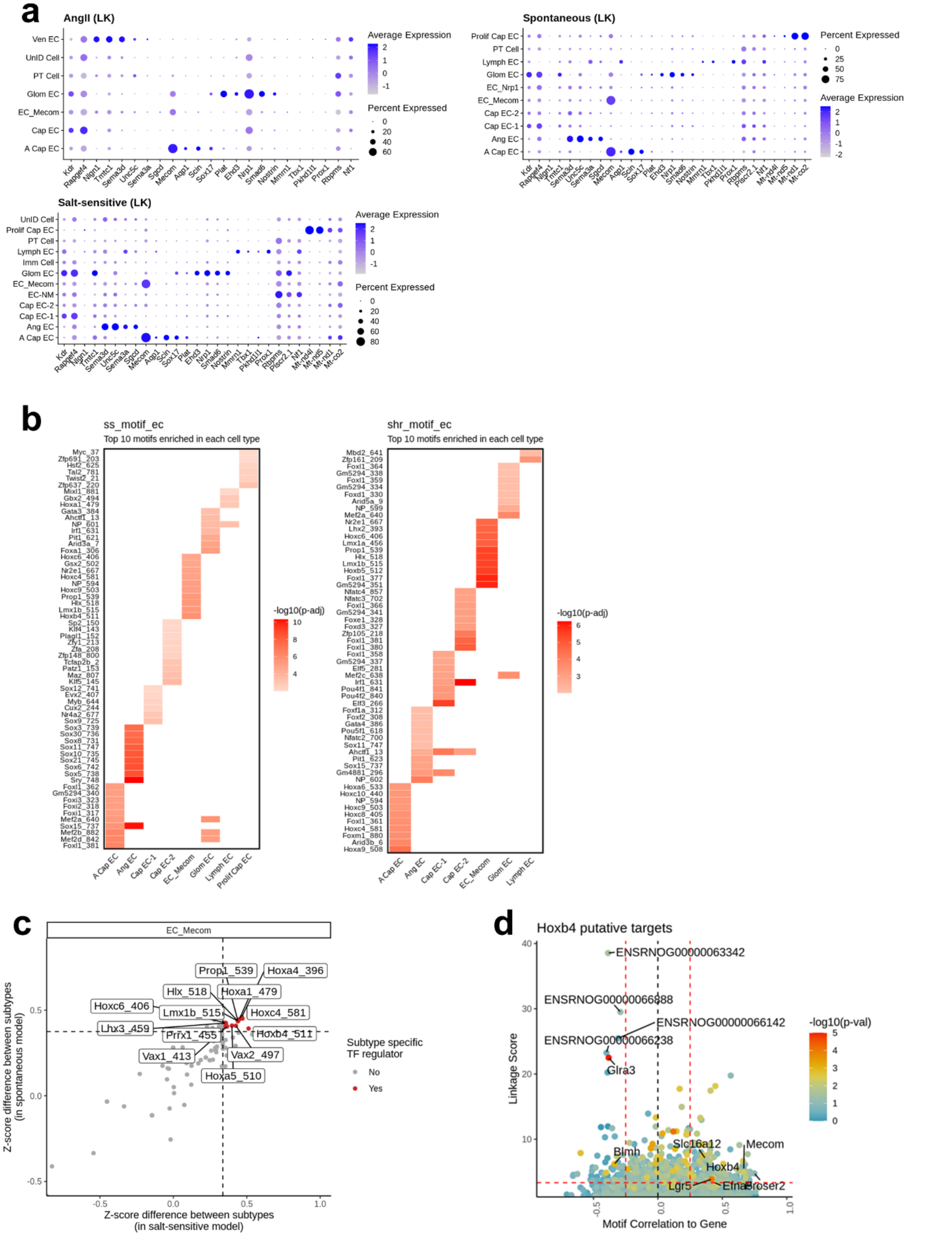


**Figure S12. snMultiome-seq based analysis of marker gene expression, motif enrichment and TF-gene interactions in kidney endothelial cell subtypes.** (a) Dot plot showing marker gene expression in LK EC subtypes. (b) Top 10 motifs enriched by peaks associated with marker genes of each EC subtype in salt-sensitive (left) and spontaneous (right) models. (c) Dot plot showing TF motifs specifically enriched in *Mecom*+ endothelial cells (ECs) in both the salt-sensitive (x-axis) and spontaneous (y-axis) models, as measured by the motif activity deviation z-score difference between *Mecom*+ ECs and the remaining ECs. Horizontal and vertical dashed lines indicate a z-score difference of 0.3. TF motifs with a z-score difference greater than 0.3 in both models are highlighted in red. (d) Prioritization of gene targets for *Hoxb4* in salt-sensitive model. The x-axis represents the Pearson correlation between TF motif activity and integrated gene expression across endothelial cells. The y-axis shows the TF linkage score, calculated as the sum of scaled motif scores for all linked peaks based on peak-to-gene link correlation. The color of the points reflects the hypergeometric enrichment of the TF motif in linked peaks for each gene. The horizontal dashed line marks the 80th percentile of the linkage score, while the vertical dashed lines represent motif-to-gene expression correlations of -0.25 and 0.25, respectively.

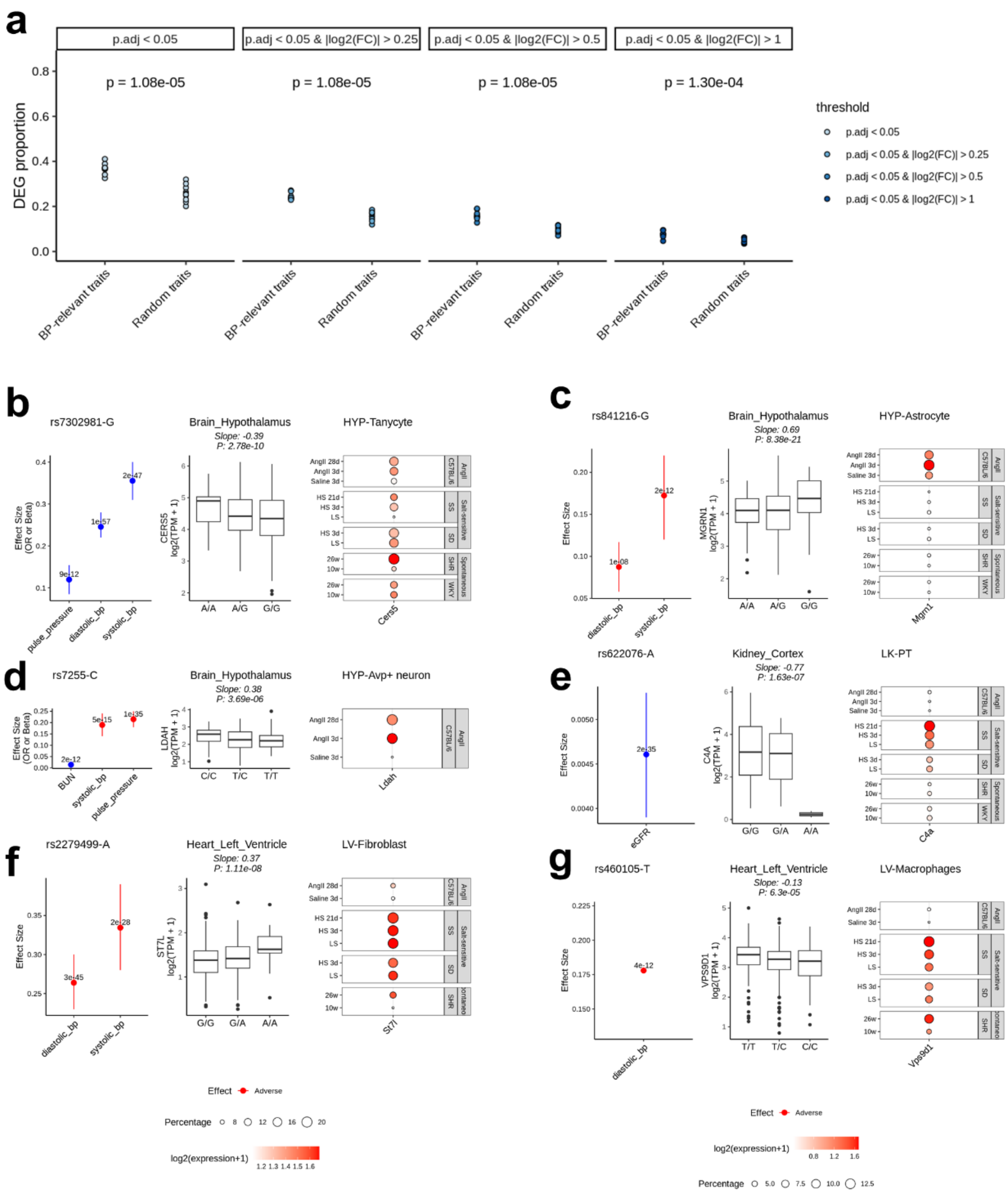


**Figure S13. Overlap of DEGs and SNP-related genes across traits and SNP-eQTL associations in BP-relevant traits.** (a) Proportion of overlap between DEGs and SNP-related genes across traits, grouped by BP-relevant and BP-irrelevant traits, with DEGs categorized by different standards in each column. (b-g) Relationship between selected SNPs and eQTL gene expression: rs7302981-G with *CERS5* (b), rs841216-G with *MGRN1* (c), rs7255-C with *LDAH* (d), rs622076-A with *C4A* (e), rs2279499-A with *ST7L* (f), and rs460105-T with *VPS9D1* (g). Each panel includes: the effect size (95% CI) of the specific SNP allele on BP-relevant traits (left), box plots showing the association between SNP genotypes and gene expression from GTEx data in a relevant tissue (middle), and the expression of the gene in specific cell types in the corresponding tissue under different conditions from our snRNA-seq data (right).

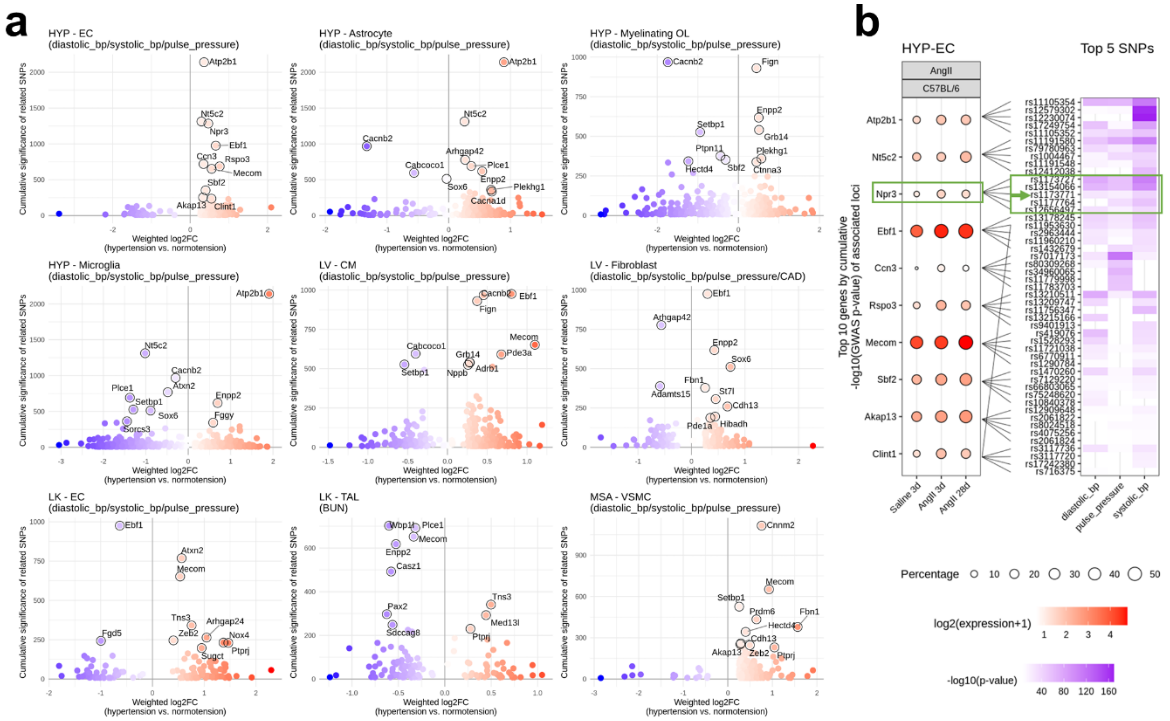

**Figure S14. Association of genes with blood pressure traits.** (a) Scatter plot showing the association of genes with multiple blood pressure traits. The y-axis represents the cumulative significance of related SNPs, while the x-axis shows the weighted expression change of these genes in selected cell types during hypertension. The top 10 genes with the highest cumulative significance are highlighted. Cell types were selected based on Fig. 4e. (b) Expression levels of the top 10 genes with the highest cumulative significance in hypothalamus endothelial cells across conditions in the AngII model (left), alongside their top 5 associated SNPs (right).

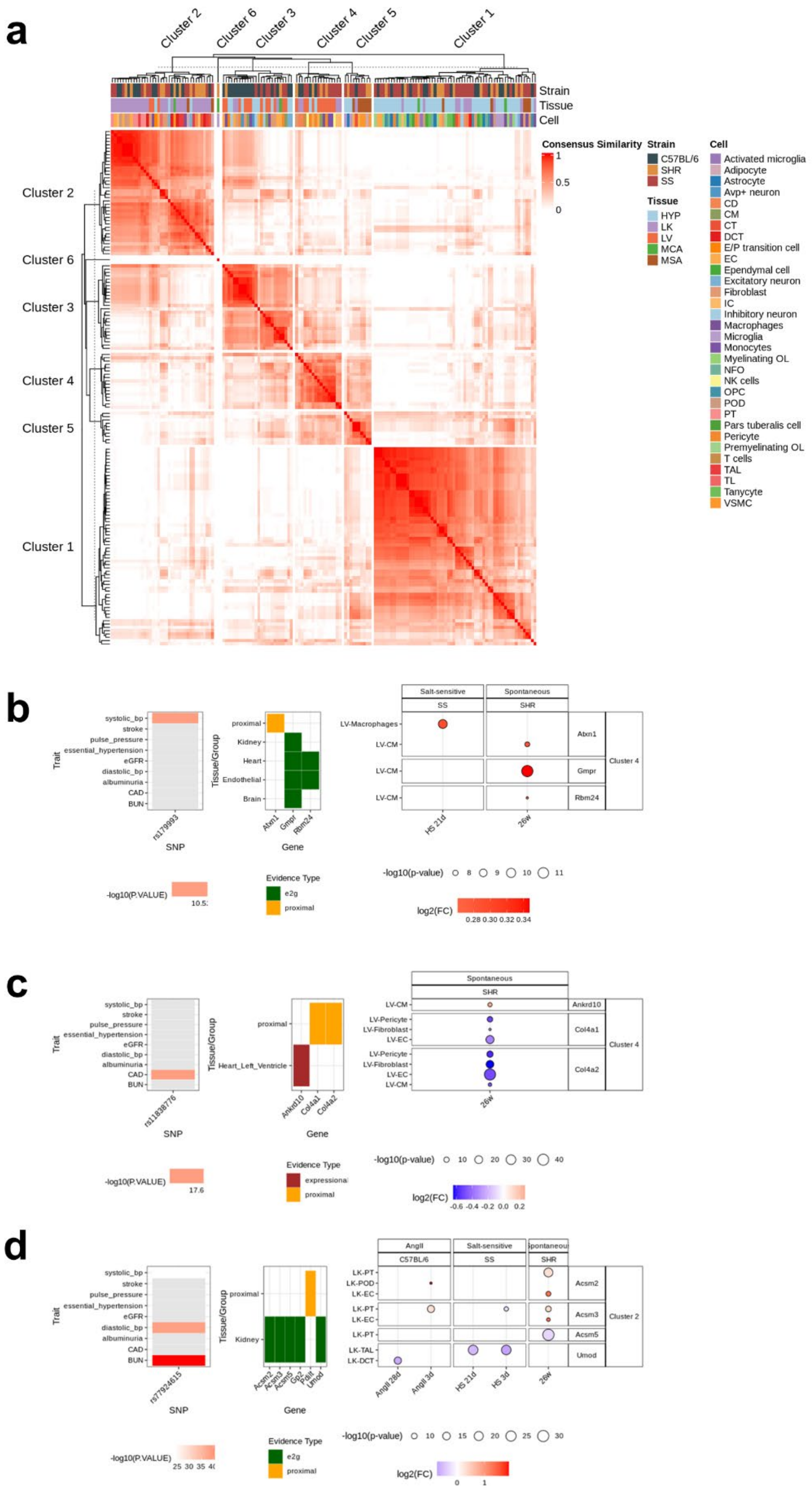

a
Cluster 2
Cluster 6
Cluster 3
Cluster 4
Cluster 5
Cluster 1
Strain
Tissue
Cell
Consensus Similarity
1
0.5
0
Strain
C57BL/6
SHR
SS
Tissue
HYP
LK
LV
MCA
MSA
Cell
Activated microglia
Adipocyte
Astrocyte
Avp+ neuron
CD
CM
CT
DCT
E/P transition cell
EC
Ependymal cell
Excitatory neuron
Fibroblast
IC
Inhibitory neuron
Macrophages
Microglia
Monocytes
Myelinating OL
NFO
NK cells
OPC
POD
PT
Pars tuberalis cell
Pericyte
Premyelinating OL
T cells
TAL
TL
Tanycyte
VSMC
b
Trait
systolic_bp
stroke
pulse_pressure
essential_hypertension
eGFR
diastolic_bp
albuminuria
CAD
BUN
rs179993
SNP
Tissue/Group
proximal
Kidney
Heart
Endothelial
Brain
Atxn1
Gmpr
Rbm24
Gene
Salt-sensitive
SS
Spontaneous
SHR
LV-Macrophages
LV-CM
Atxn1
Gmpr
Rbm24
Cluster 4
HS 21d
26w
-log10(P.VALUE)
10.5
Evidence Type
e2g
proximal
-log10(p-value)
8
9
10
11
log2(FC)
0.28 0.30 0.32 0.34
c
rs11838776
Heart_Left_Ventricle
Ankrd10
Col4a1
Col4a2
LV-Pericyte
LV-Fibroblast
LV-EC
17.6
expressional
-0.6 -0.4 -0.2 0.0 0.2
10
20
30
40
d
rs7924615
AngII
C57BL/6
LK-PT
LK-POD
LK-EC
LK-TAL
LK-DCT
Acsm2
Acsm3
Acsm5
Gls2
Pdilt
Umod
Cluster 2
AngII 28d
AngII 3d
HS 3d
25 30 35 40
15

**Figure S15. SNP-related gene based cell type cluster identification and SNP-cell type cluster association.** (a) Consensus similarity heatmap of cell types across strains, conditions, and tissues, based on a presence-absence matrix of differentially expressed SNP-related genes across various models and conditions. (b-d) SNP-cell type cluster associations via linked genes supported by multiple lines of evidence: rs179993 in Cluster 4 (b), rs11838776 in Cluster 4 (c), and rs77924615 in Cluster 2 (d). Each panel includes: GWAS association of the SNP with all BP-relevant traits (left), SNP-gene association supported by different evidence (middle), and gene expression changes across multiple cell types within the specific cluster (right).

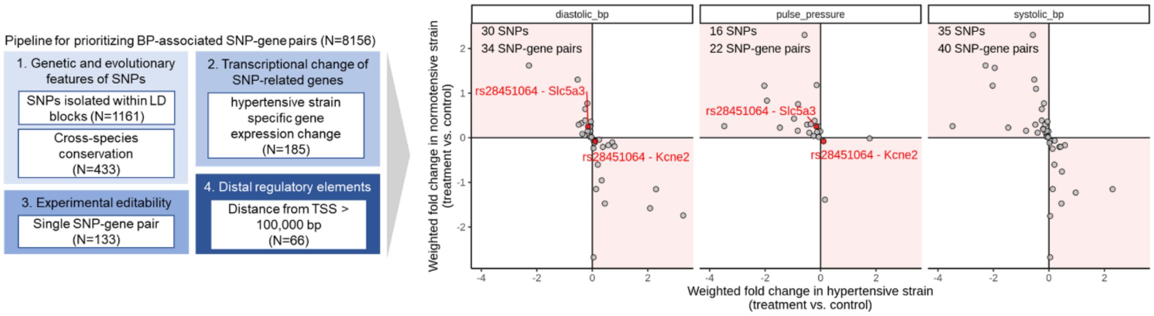


**Figure S16. Pipeline for prioritizing BP traits-associated SNP-gene pairs (left). Scatter plot showing the expression changes of filtered genes in hypertensive and normotensive strains using weighted fold change, categorized by the trait associated with the corresponding SNP (right).**

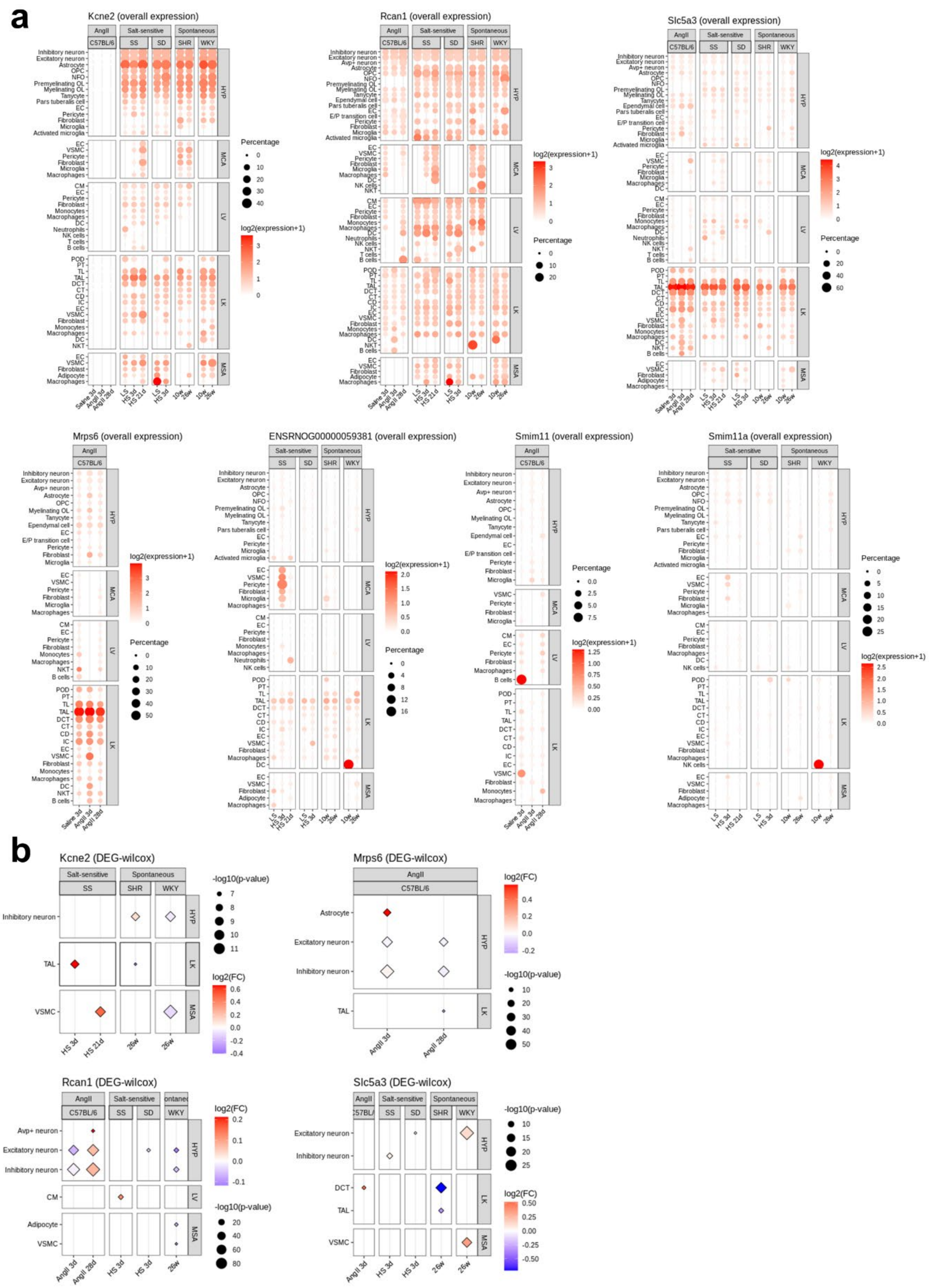
a
Kcne2 (overall expression)
Rcan1 (overall expression)
Slc5a3 (overall expression)
Mrps6 (overall expression)
ENSRNOG00000059381 (overall expression)
Smim11 (overall expression)
Smim11a (overall expression)
AngII
C57BL/6
Salt-sensitive
SS
SD
Spontaneous
SHR
WKY
HYP
MCA
LV
LK
MSA
Percentage
log2(expression+1)
Saline 3d
AngII 3d
AngII 28d
LS
HS 3d
HS 21d
10w
26w
b
Kcne2 (DEG-wilcox)
Mrps6 (DEG-wilcox)
Rcan1 (DEG-wilcox)
Slc5a3 (DEG-wilcox)
-log10(p-value)
log2(FC)
Inhibitory neuron
Excitatory neuron
Avp+ neuron
Astrocyte
TAL
DCT
VSMC
CM
Adipocyte

**Figure S17. Expression of genes neighboring rs28451064**. (a) Dot plot showing the expression levels of selected genes across cell types in each tissue (y axis) and treatment in each model (x axis). (b) Differential expression results of selected genes across treatments in each model, based Wilcoxon test. Only genes with a Bonferroni-adjusted p-value less than 0.05 were shown, and highlighted with diamond shapes.

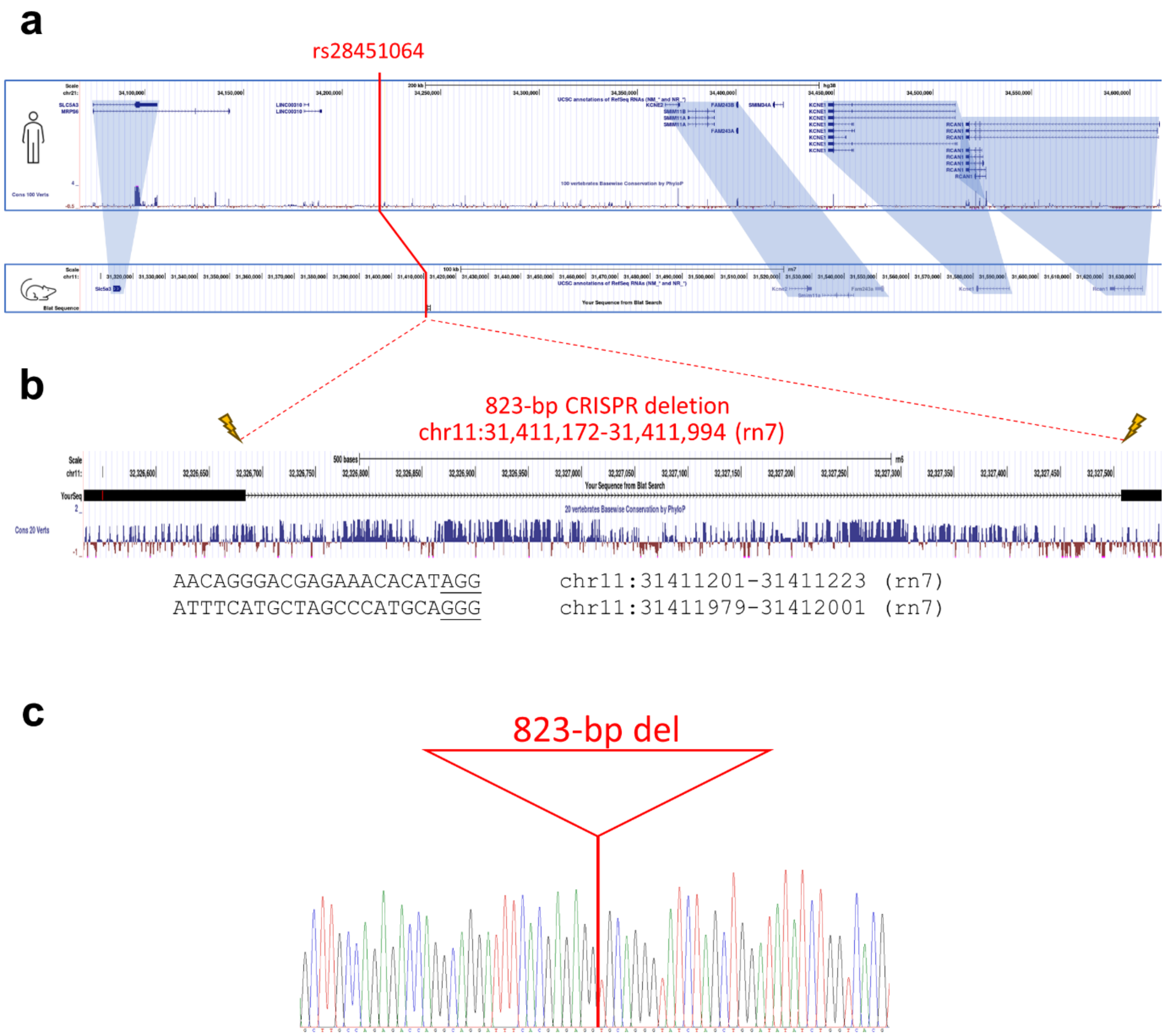


**Figure S18. Comparative analysis and generation of a rat model for rs28451064.** (a) Comparative map alignment between human and rat. The genes and order around rs28451064 are syntenic. (b) CRISPR-SpCas9 sgRNA targets flanking a ~800-bp region in rat semiconserved at the sequence level with the human rs28451064 locus. An 823-bp deletion in chromosome 11 was generated. (Note: The corresponding LiftOver coordinates from rn7 to rn6 were used to show vertebrate sequence conservation in this 823-bp region.) (c) Sanger sequence confirmation of the 823-bp deletion.

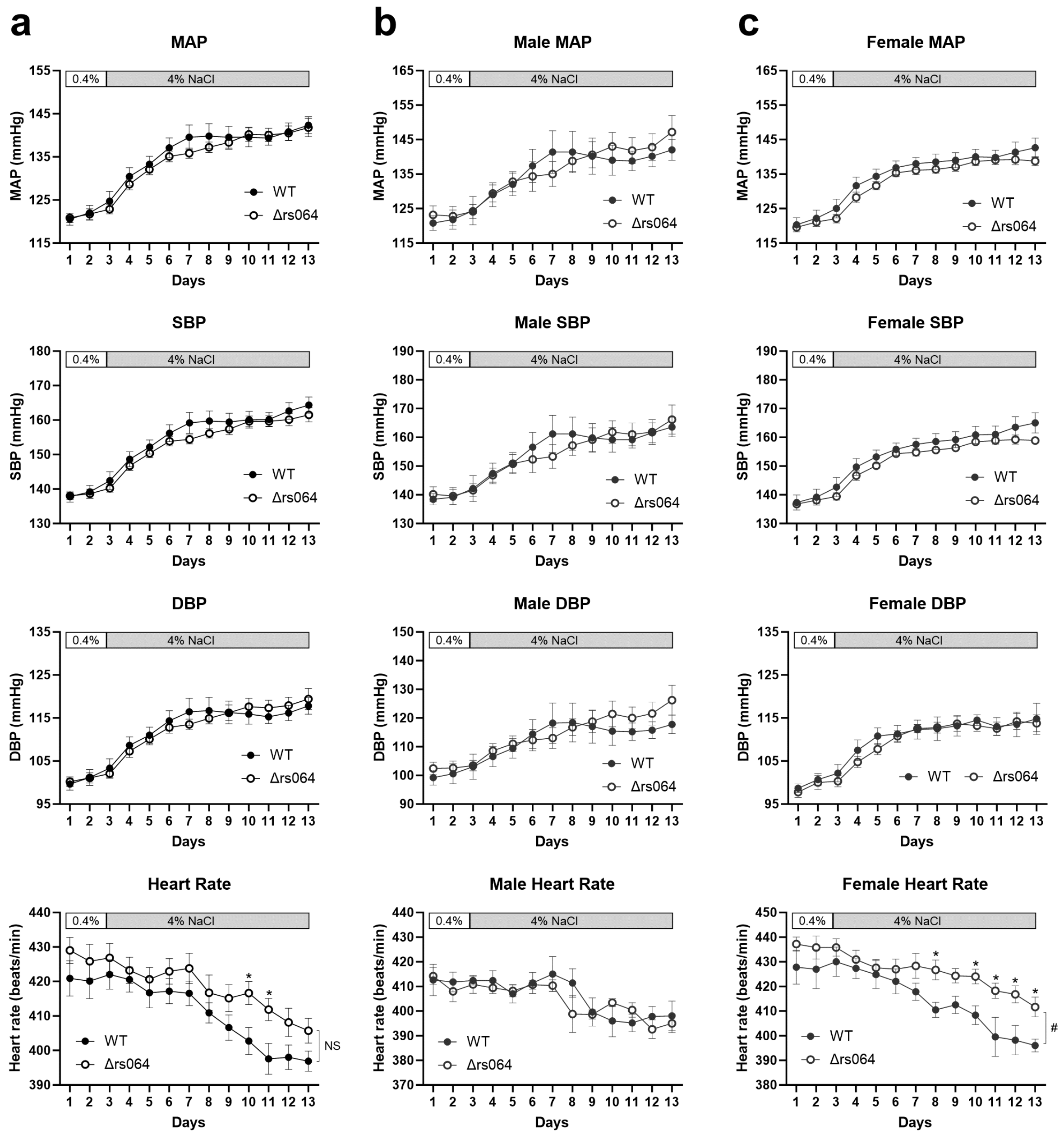


**Figure S19. Deletion of the rs28451064 orthologous region does not change mean arterial pressure (MAP), systolic blood pressure (SBP), diastolic blood pressure (DBP), or heart rate in SS rats.** (a) Data with male and female rats combined. N=12 and 14. (b) Male rats. N= 5 and 5. (c) Female rats. N=7 and 9.

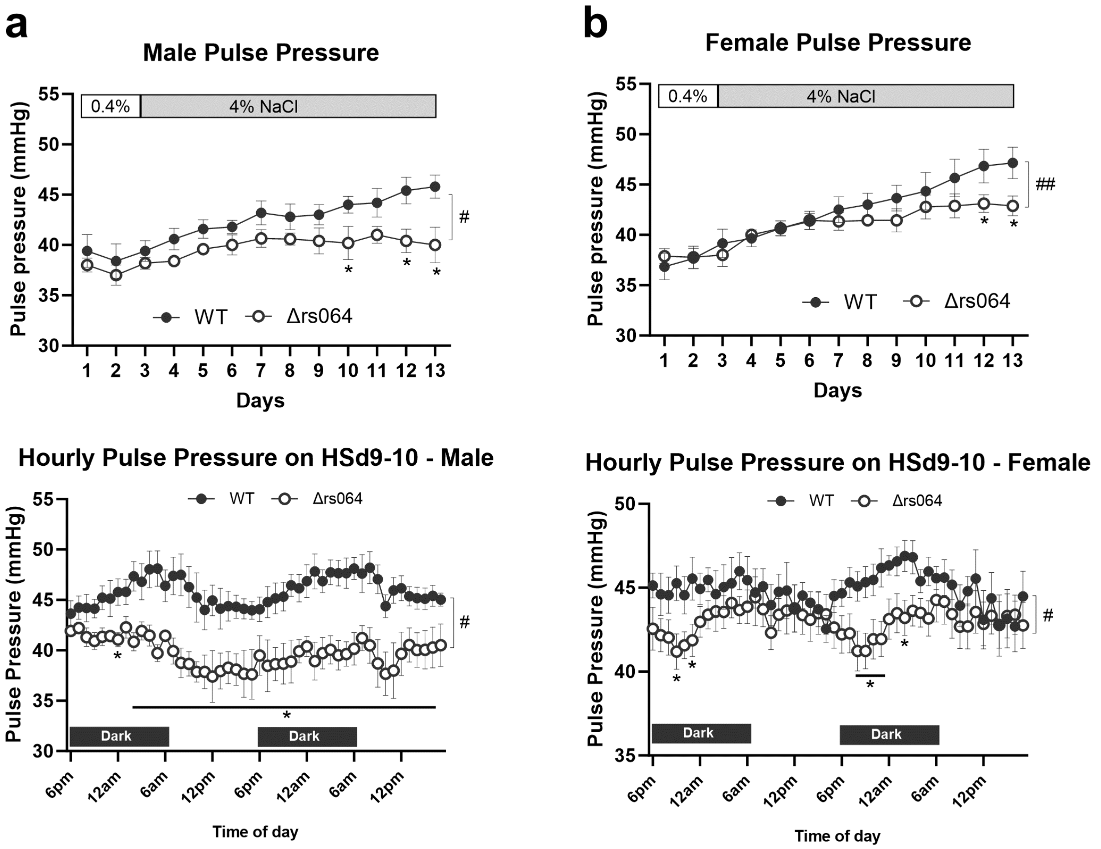


**Figure S20. Deletion of the rs28451064 orthologous region decreases pulse pressure in both male and female SS rats on a 4% NaCl diet.** (a) Male rats. N= 5 and 5. (b) Female rats. N=7 and 9. #, $p<0.05$ for WT vs. delta-rs064; *, $p<0.05$ vs. WT; two-way repeated measure ANOVA followed by Holm-Sidak test.

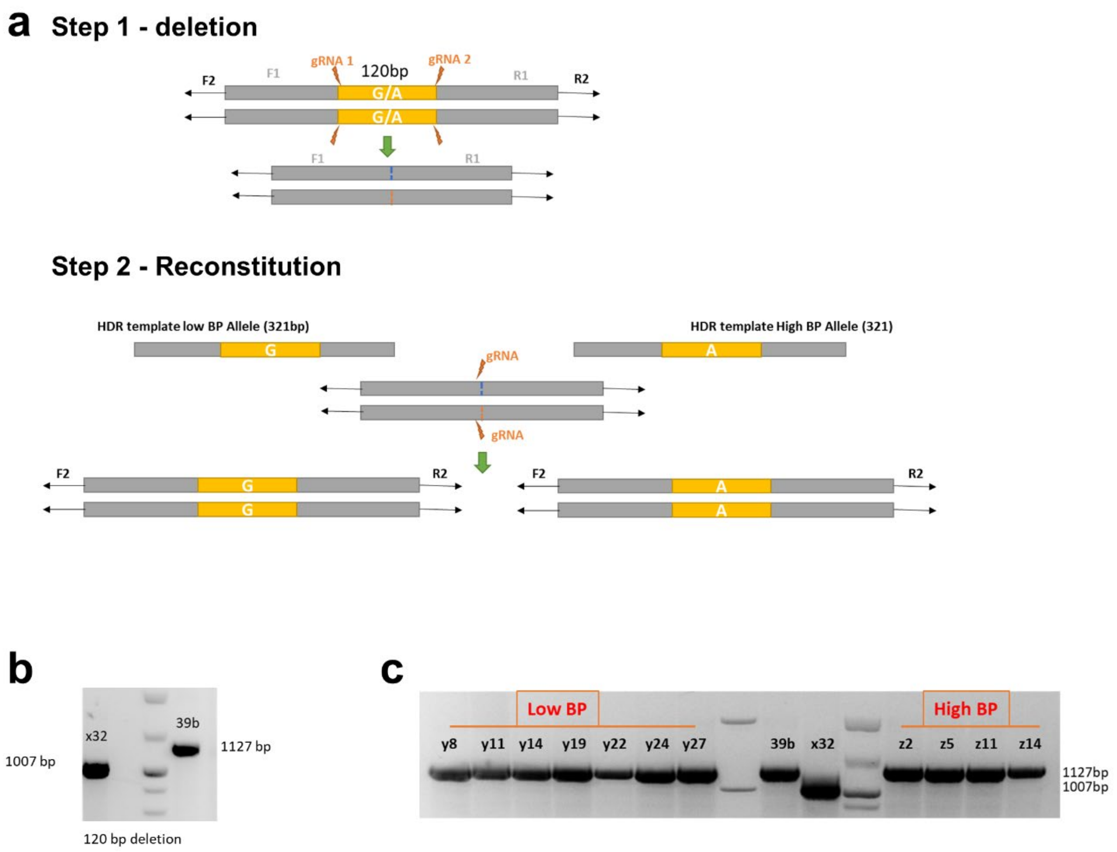


**Figure S21. Generation of isogenic hiPSCs containing homozygous rs28451064-G (low DBP allele, or larger PP allele) or rs28451064-A (high DBP allele, or smaller PP allele).** (a) Schematic of the two-step genome editing process. (b) Deletion of DNA segment containing rs28451064 locus. 39b – original iPSC cell line, X32 - SNP rs28451064 deleted cell line. (c) Reconstitution of SNP rs28451064 locus containing either homozygous larger PP allele or smaller PP allele. 39b – original iPSC cell line, X8 - rs28451064 deleted cell line, y series – reconstituted rs28451064 larger PP allele cell line, z series – reconstituted rs28451064 smaller PP cell line.

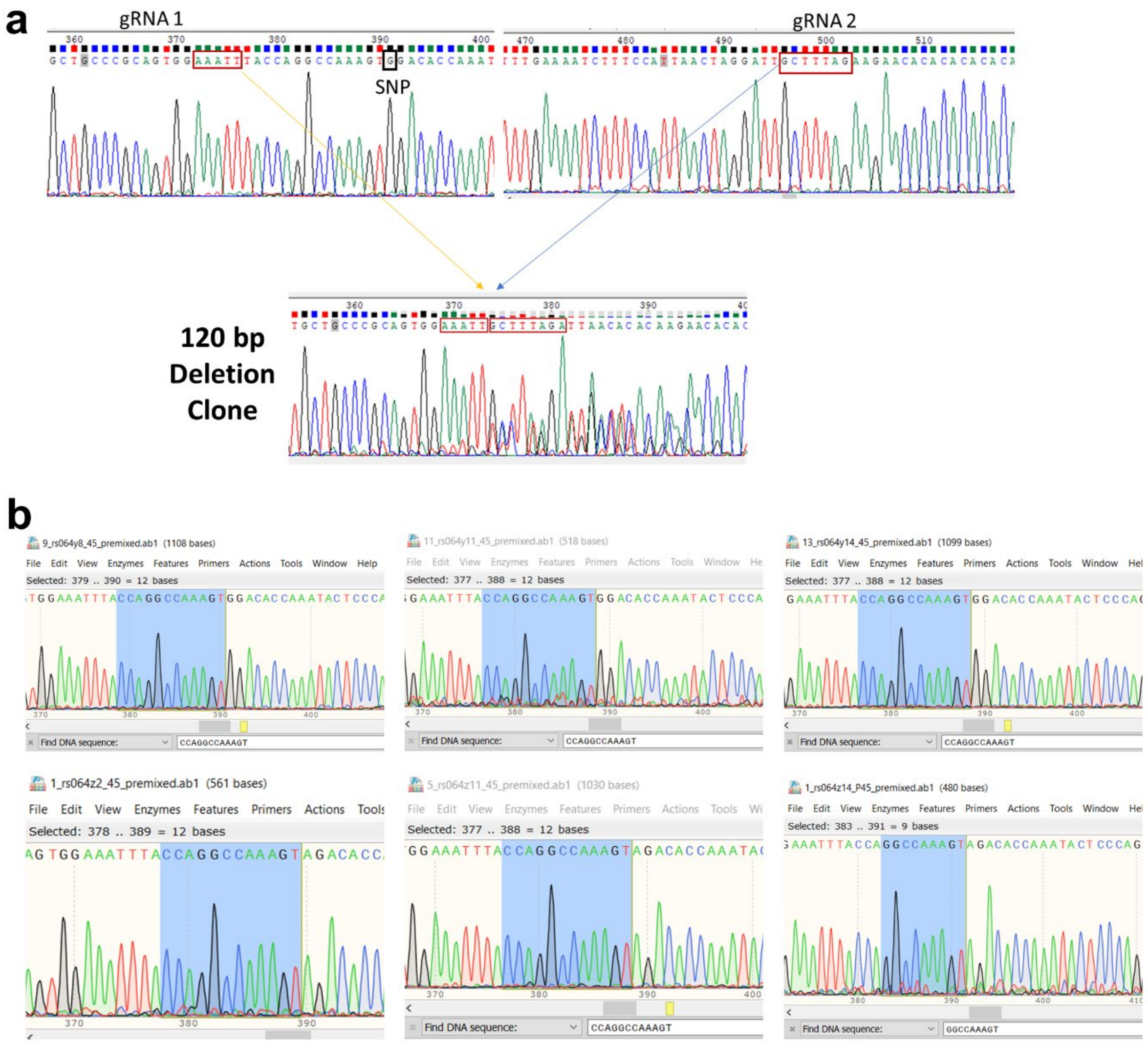


**Figure S22. Sanger sequencing confirmation of hiPSC editing.** (a) Sanger sequencing confirming deletion of a 120 bp region containing rs28451064. (b) Sanger sequencing confirming reconstitution of either homozygous larger PP (top row, three clones) or smaller PP (bottom row, three clones) allele of rs28451064. Larger PP allele cell lines – y8, y11 and y14. Smaller PP allele cell lines – z2, z11 and z14.

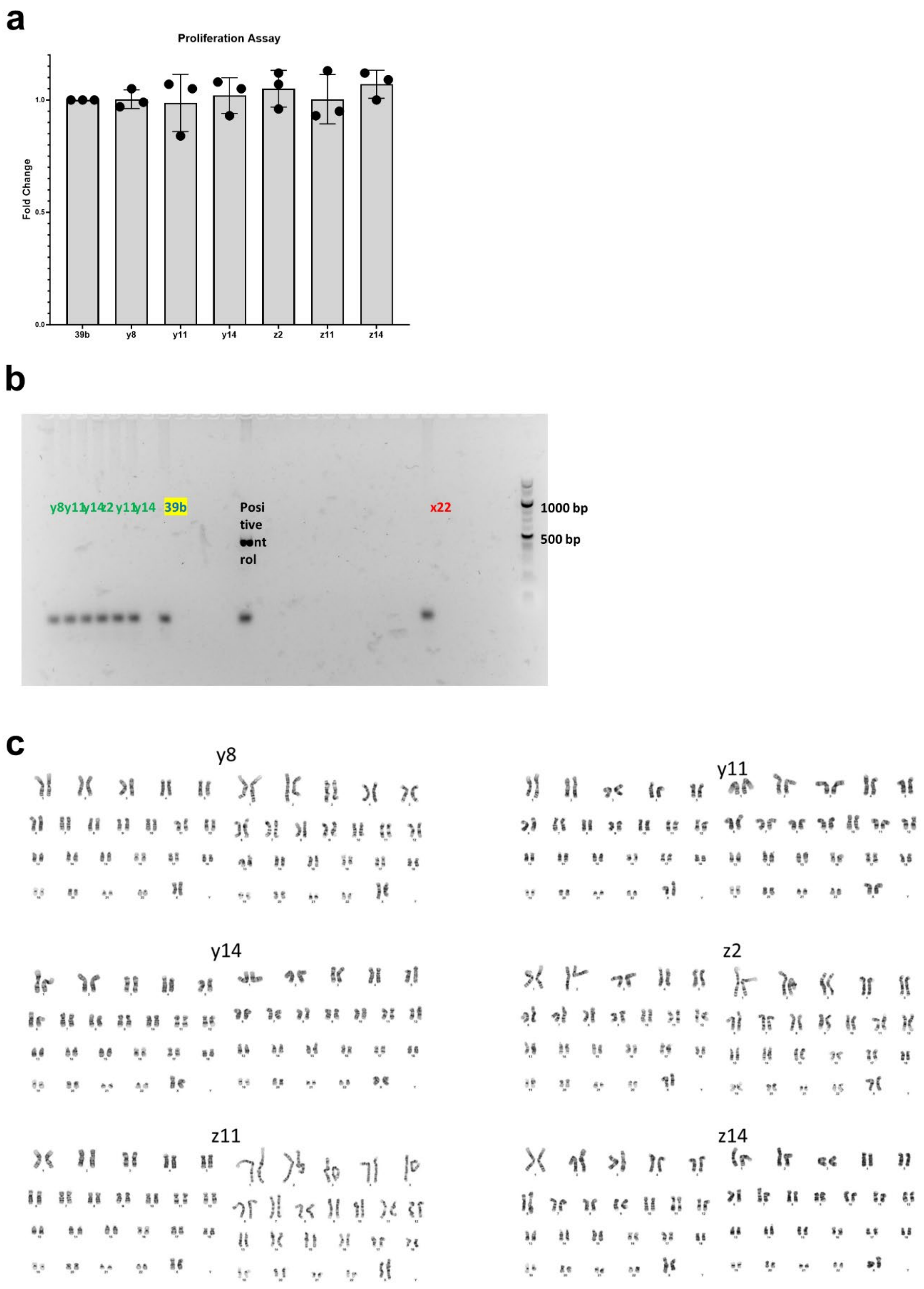


**Figure S23. Quality assessment of isogenic hiPSCs containing homozygous rs28451064-G (larger PP allele) or rs28451064-A (smaller PP allele).** (a) Proliferation assay. Equal number of cells were seeded on day 0 in 6cm dish for reconstituted clones. Cells were collected on day 4 and total cell number for reconstituted iPSCs were compared with original iPSC cell line, 39b. Larger PP allele cell lines – y8, y11 and y14. Smaller PP allele cell lines – z2, z11 and z14. (b) Mycoplasma contamination screening for reconstituted iPSCs. x22 is another mycoplasma negative iPSC cell line. Original iPSC cell line - 39b. (c) Representative karyograms for three Large PP (y8, y11, y14) and three smaller PP (z2, z11 and z14) clones. Chromosomes of 20 proliferating cells were counted except for clone Z8 where 10 proliferating cells were counted and fully analyzed using G-banding. Three cells were karyotyped. These cells had a modal number of 46 chromosomes. The sex chromosome constitution indicates normal female. No consistent abnormalities were observed in the chromosomal number or banding patterns.

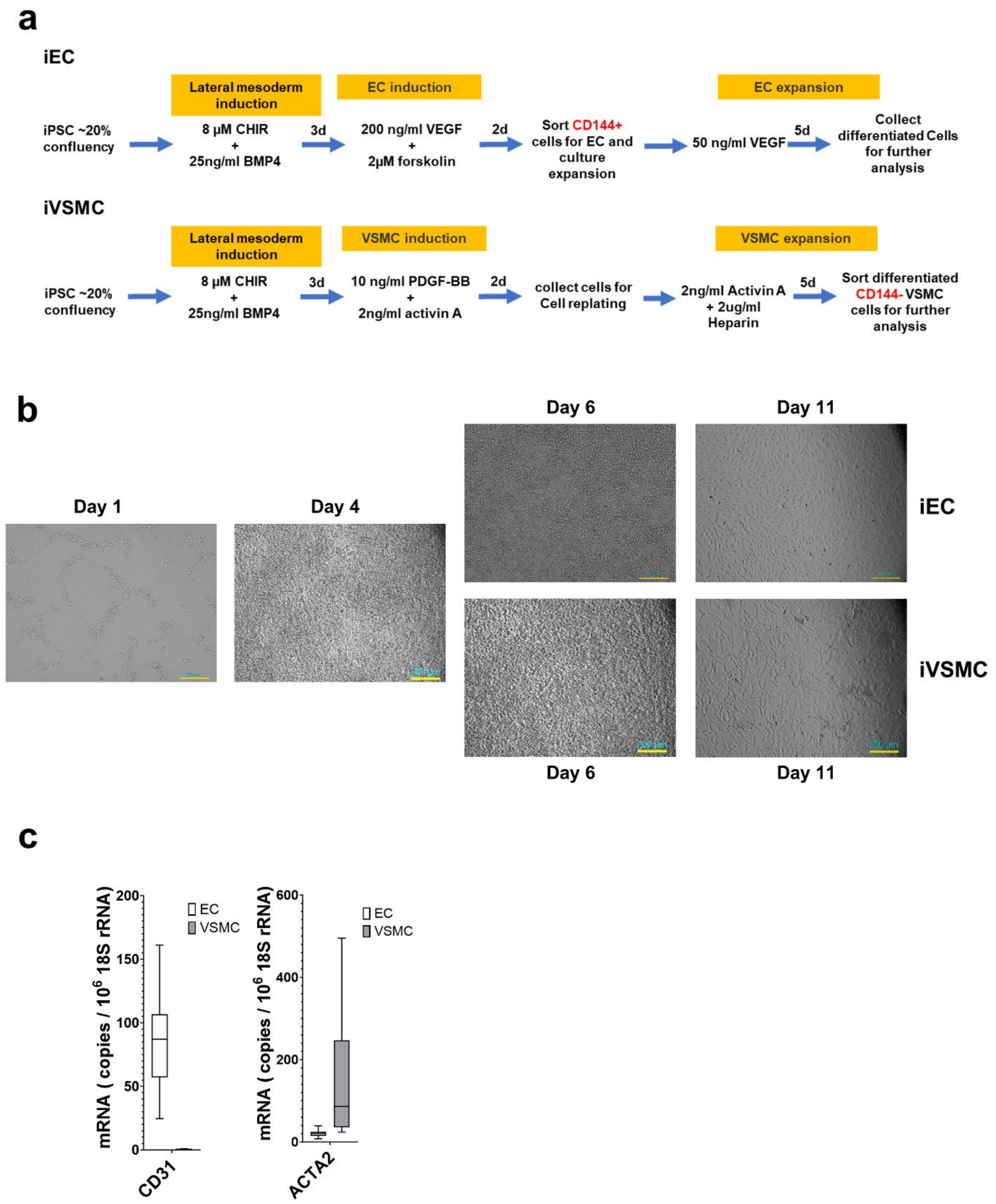


**Figure S24. Differentiation of edited hiPSCs to endothelial cells (iECs) and vascular smooth muscle cells (iVSMCs).** (a) Differentiation protocol adapted from Patsch et al., Nat Cell Biol, 2015[3]. (b) Representative DIC images of iEC and iVSMC differentiation on day 1, day 4, day 6 and day 11. Scale bar – 200 μm. (c)

Expression of EC marker CD31 and VSMC marker ACTA2 in iECs and iVSMCs. N = 66 for iEC and N = 72 for iVSMC.

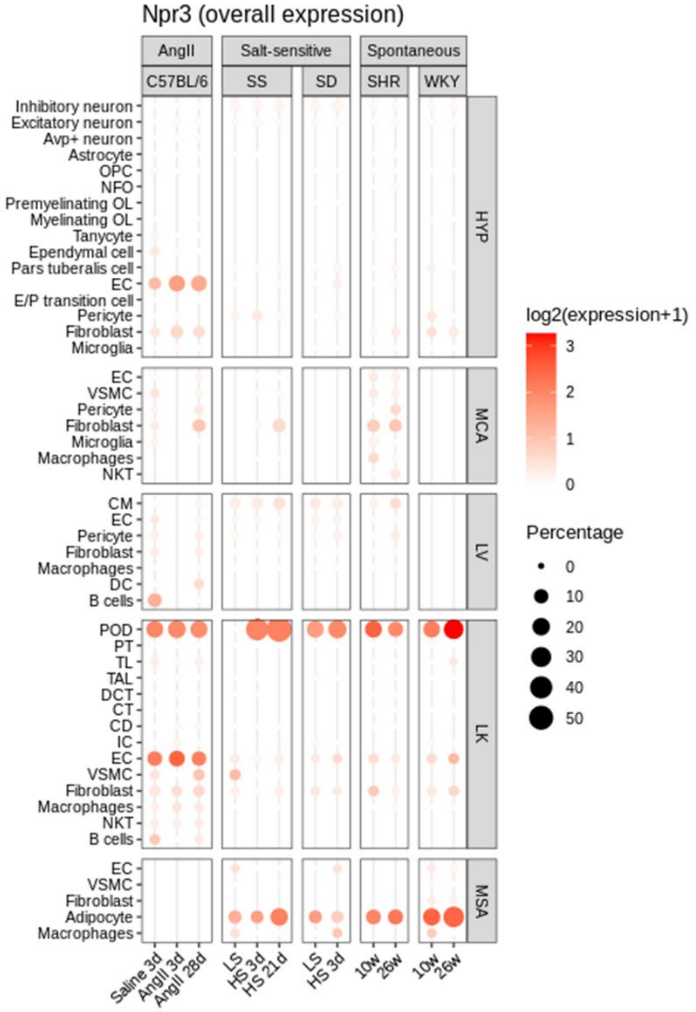


**Figure S25. Dot plot showing the expression levels of *Npr3* across cell types in each tissue (y axis) and treatment in each model (x axis) in our snRNA-seq data.**

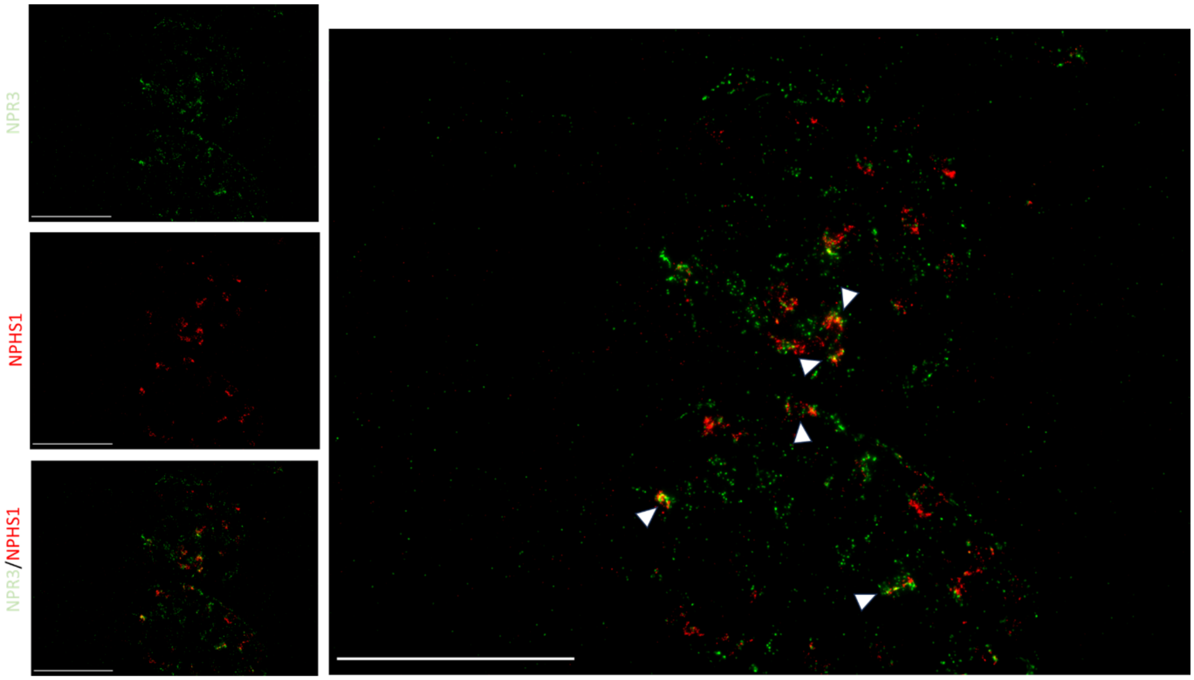


**Figure S26. *Npr3* is expressed in podocytes.** Representative images from multiplex RNAScope showing the spatial distribution and co-localization of *Nphs1* (red; a podocyte marker) and *Npr3* (green) transcripts in kidney sections from SS rats. The top left panel displays *Npr3* transcripts, the middle left panel shows *Nphs1* transcripts, and the bottom left panel presents a merged image indicating areas of co-localization (yellow). The large right panel provides a zoomed-in view, where white arrowheads highlight regions of strong co-localization between *Nphs1* and *Npr3*, demonstrating the expression of *Npr3* in podocytes. Scale bars = 100 µm.

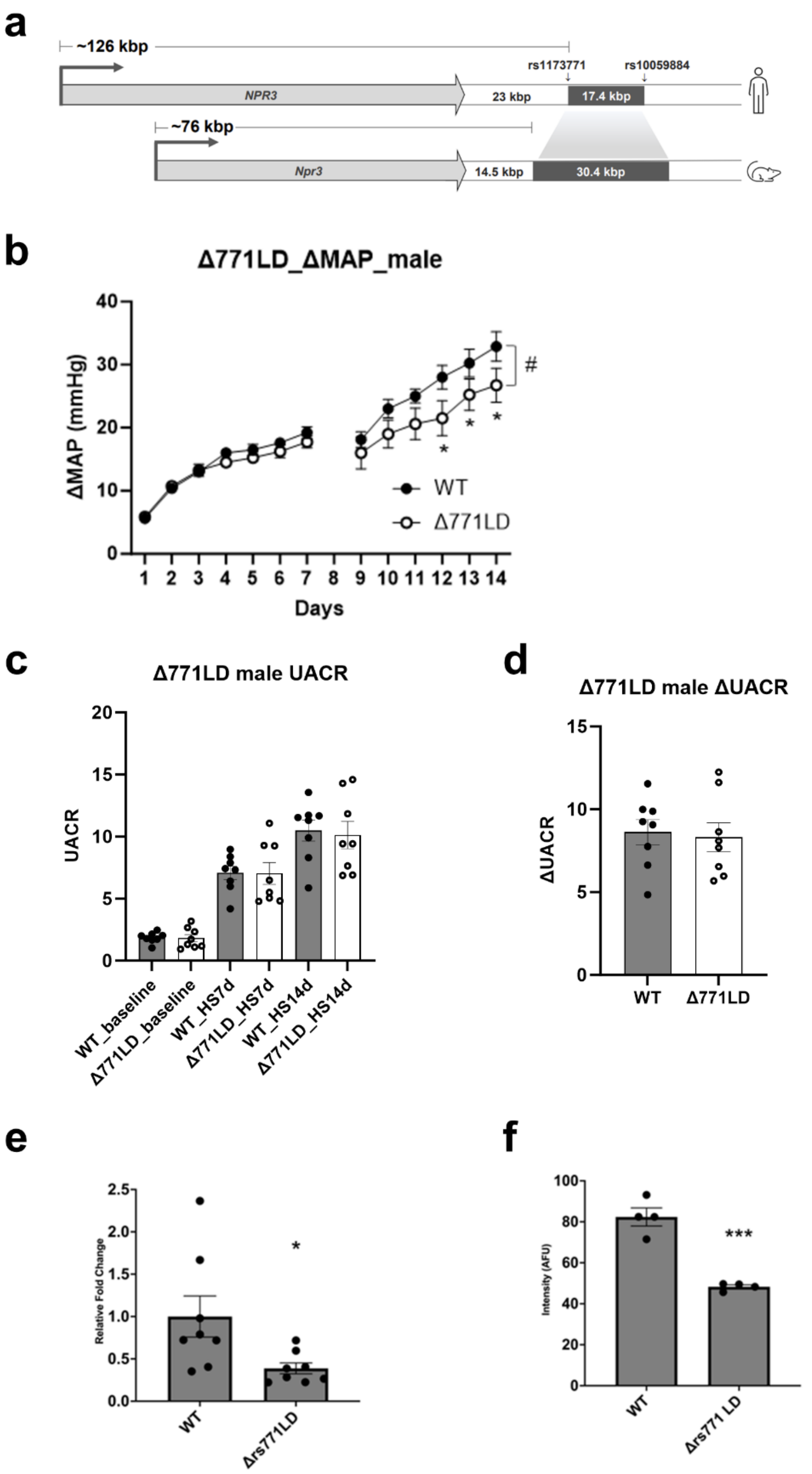


**Figure S27. Deletion of the rs1173771 haplotype orthologous region decreased NPR3 expression in podocytes and increased the sensitivity of SS rats to hypertension-induced albuminuria.** (a) Diagram depicting the rat genomic region orthologous to the human rs1173771 linkage disequilibrium (LD)

region. A 30.4 kbp region orthologous to the 17.4 kbp human rs1173771 LD region was deleted from the genome of Dahl salt sensitive (SS) rats to generate SS-Δrs1173771LD rats[4]. The transcription start site of the closest protein-coding gene, *NPR3*, is approximately 126 kbp from the haplotype region in human genome and ~76 kbp in rat genome. (b) The increase of mean arterial pressure (ΔMAP) after diet switch to 4% NaCl, relative to baseline MAP on a 0.4% NaCl diet, was significantly attenuated in male SS-Δrs1173771LD rats (Δ771LD) compared with wild type (WT) littermate SS rats. The graph was re-plotted from data reported in Xue, et al, bioRxiv 2024[4]. N = 8 per group. #, $p < 0.05$ by two-way RM ANOVA, *, $p < 0.05$ by Holm-Sidak test. (c) Urinary albumin to creatinine ratio (UACR) in SS-Δrs1173771LD$^{-/-}$ rats was not different than WT littermates on the 0.4% NaCl baseline diet or 4% NaCl high salt (HS) diet. N = 8. (d) Changes of UACR from the baseline 0.4% NaCl diet to HS for 14 days were not different between SS-Δrs1173771LD$^{-/-}$ rats and WT littermates. N = 8. (e) *Npr3* expression in isolated glomeruli was decreased in SS-Δrs1173771LD$^{-/-}$ rats, based on qPCR. N = 8; *, $p < 0.05$, unpaired t-test. (f) *Npr3* expression in podocytes was decreased in SS-Δrs1173771LD$^{-/-}$ rats, based on RNAScope. Co-localization of *Nphs1*, a podocyte-specific marker, with *Npr3* was used to assess the expression of *Npr3* in podocytes. Co-localization analysis was performed with ImageJ using split-channel thresholded binary masks and image calculator "multiply" function to locate only pixels present in both channels. 4 rats per group and 18 glomeruli per rat were examined. *** $p < 0.001$, unpaired t-test.

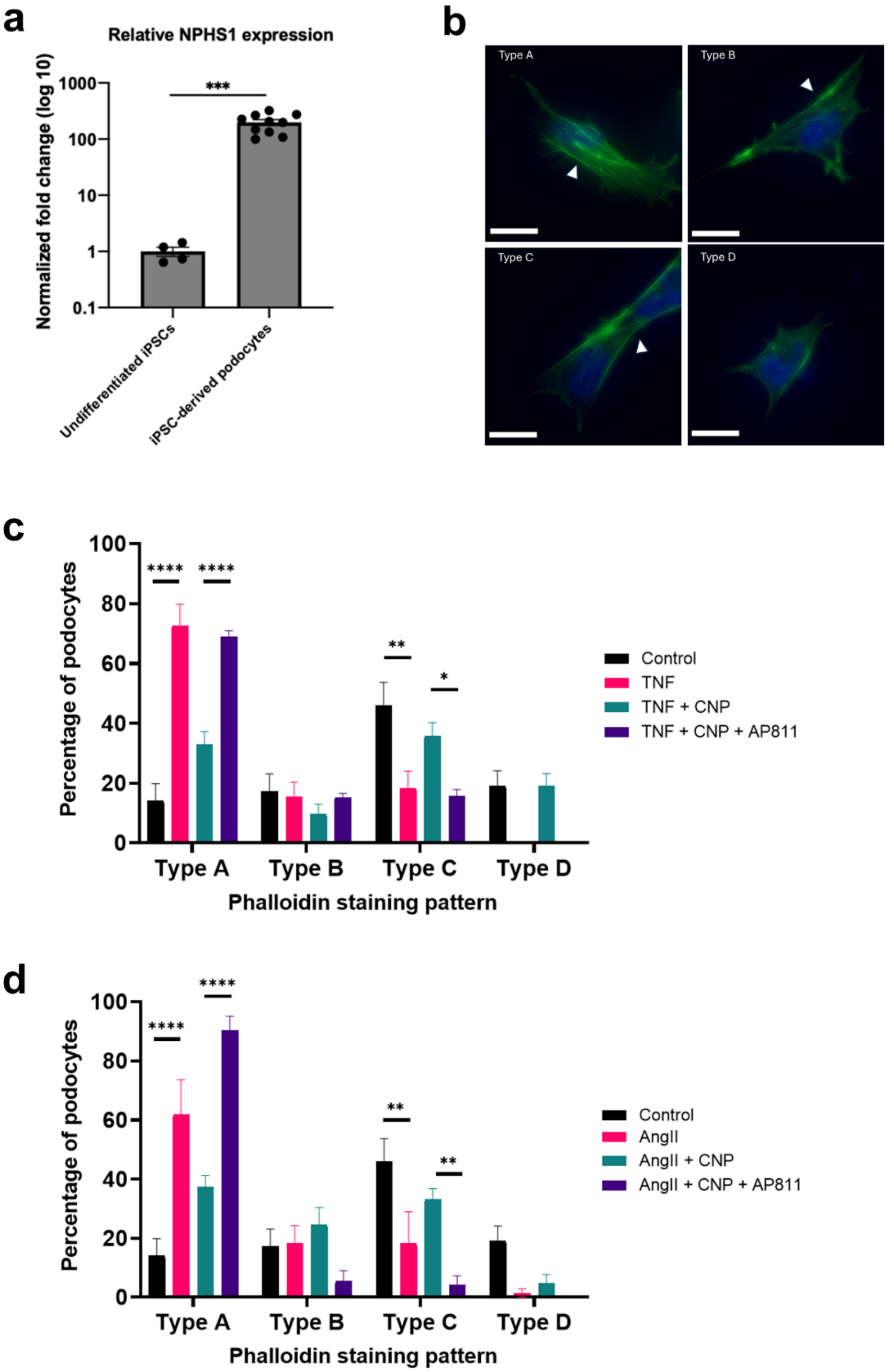


**Figure S28. C-type natriuretic peptide (CNP), acting through natriuretic peptide receptor C (NPRC), protects podocytes.** (a) hiPSC-derived podocytes express podocyte marker *NPHS1*. The expression of *NPHS1*, normalized to 18S rRNA, is elevated by approximately 200-fold in hiPSC-derived podocytes compared to undifferentiated hiPSCs. ***, $p < 0.001$, unpaired t-test. (b) Representative images of different types of phalloidin staining patterns observed in hiPSC-derived podocytes. Cells were treated with TNFα or AngII alone and in combination with CNP or CNP + AP811. Phalloidin staining was used to assess actin cytoskeleton reorganization of the podocytes in response to the treatments. White arrows highlight representative stress fibers. Scale bars = 20 µm. Type A: >90% of cell area filled with thick stress fibers. Type C: no thick cables, but some fibers present. (c) Quantification of phalloidin staining in hiPSC-derived podocytes following a 24-hour treatment of 20 ng/mL TNFα with two additional interventions. TNFα significantly increased the proportion of Type A cells and decreased the proportion of Type C cells. Co-treatment with 100 nM CNP attenuated the effect of TNFα. The addition of 100 nM AP811, an antagonist of NPRC, reversed the protective effect of CNP. N = 6-8. * = $p < 0.05$, ** = $p < 0.01$, **** = $p < 0.0001$, two-way ANOVA followed by Holm-Sidak test. (d) AngII treatment (1 µM) produced a similar increase in stress fibers, with CNP co-treatment reducing stress fibers and AP811 reversing that effect. N = 6-8. ** = $p < 0.01$, **** = $p < 0.0001$, two-way ANOVA followed by Holm-Sidak test.
